# A Composition-Dependent Molecular Clutch Between T Cell Signaling Condensates and Actin

**DOI:** 10.1101/316414

**Authors:** J.A. Ditlev, A.R. Vega, D.V. Köster, X. Su, T. Tani, A.M. Lakoduk, R.D. Vale, S. Mayor, K. Jaqaman, M.K. Rosen

## Abstract

Biomolecular condensates play important roles in eukaryotic cells by concentrating molecules into foci without a surrounding membrane. During T cell activation, biomolecular condensates form at the immunological synapse (IS) through multivalency-driven phase separation of the adaptor protein LAT and its binding partners Grb2, Sos1, SLP-76, Nck and WASP. These condensates move radially at the IS, traversing a radially-oriented and then a concentric actin network. To understand the persistent radial movement, we biochemically reconstituted LAT condensates with mobile actomyosin filaments. We found that basic regions of Nck and N-WASP promote strong association and co-movement of LAT condensates with actin. Condensates lacking these components were instead propelled by steric interactions. In cells, LAT condensates lost Nck while traversing the boundary between the two actin networks, and condensates engineered to constitutively bind actin moved aberrantly. We propose that Nck and WASP form a clutch between LAT condensates and actin, and changes in composition enable condensate movement by distinct actin networks in different regions of the IS.

## Introduction

Biomolecular condensates are compartments in eukaryotic cells that concentrate macromolecules without an encapsulating membrane (Banani et al., 2017; Shin and Brangwynne, 2017). Numerous condensates are found in the cytoplasm and nucleoplasm, where they are involved in processes ranging from mRNA storage and degradation to DNA repair and ribosome biogenesis (Brangwynne et al., 2011; Feric et al., 2016; Luo et al., 2018; Protter and Parker, 2016). They are also found at membranes, where they control the organization, and likely the activity, of many signaling receptors (Banjade and Rosen, 2014; Su et al., 2016; Zeng et al., 2018).

Condensates are thought to form through phase separation driven by multivalent interactions between molecules containing multiple binding elements (Banani et al., 2017; Banjade and Rosen, 2014; Li et al., 2012). Recent models suggest that a limited collection of proteins and/or RNA molecules forms the essential phase separating scaffold of particular condensates (Banani et al., 2016; Ditlev et al., 2018; Langdon et al., 2018). These molecules then recruit larger numbers of client proteins to complete the structure. Condensate composition is thus determined by the specificity of interactions among scaffolds and between scaffolds and clients. The concentrations, interactions, and dynamics of scaffolds and clients are believed to dictate the biochemical activities, and consequent cellular functions, of individual condensates (Banani et al., 2017; Holehouse and Pappu, 2018; Stroberg and Schnell, 2018). The composition of many condensates is known to change in response to signals, implying regulated changes in activity (Chen et al., 2008; Dellaire et al., 2006; Markmiller et al., 2018; Salsman et al., 2017; Youn et al., 2018). However, the relationships between composition and biochemical/cellular functions are not well understood in most cases.

During activation of a T cell by an antigen presenting cell (APC), condensates organized around the transmembrane adaptor protein LAT (Linker for Activation of T cells) play important roles in downstream signaling and resultant T cell activation (Balagopalan et al., 2013; Houtman et al., 2006; Kumari et al., 2015). These condensates are located at the interface between the T cell and APC, known as the immunological synapse (IS) (Bunnell et al., 2002). We recently provided evidence that these structures form through multivalency-driven phase separation of LAT and its intracellular binding partners Grb2, Sos1, SLP-76, Nck, and WASP (Su et al., 2016). In this system LAT, Grb2 and Sos1 appear to act as the key scaffolds, which recruit SLP-76, Nck, and WASP as clients. During T cell activation, LAT condensates that initially appear in the periphery of the IS are moved radially to the IS center by two different actin cytoskeletal networks, a peripheral dendritic meshwork and more central circular arcs (DeMond et al., 2008; Kaizuka et al., 2007; Mossman et al., 2005; Yu et al., 2010). This translocation is essential for proper T cell responses to antigen presenting cells (Babich et al., 2012; Ilani et al., 2009; Kumari et al., 2012; Yu et al., 2012).

It has remained unknown how LAT condensates engage actin to move across the IS and whether their composition plays a role in this process. To address these questions, we analyzed interactions between LAT condensates and actin in reconstituted biochemical systems and in cells using quantitative fluorescence microscopy and statistical analysis. Our *in vitro* studies revealed that LAT condensates bind actin filaments in a composition-dependent fashion, primarily through interactions of Nck and N-WASP. In cells, we observed that Nck dissipates from LAT condensates as they traverse the IS. This loss coincided spatially with the change in actin architecture from dendritic network to circular arcs. LAT condensates containing a mutant Grb2, which caused them to constitutively bind actin filaments independently of Nck, exhibited aberrant movement across the IS. These data suggest a model in which LAT condensates engage actin differently depending on the density of Nck and WASP proteins in them, such that switching between compositions and actin-binding modes enables them to move radially via the two actin networks at the IS.

## Results

### LAT condensates reconstituted *in vitro* within actin networks move in a composition-dependent manner

We previously reconstituted LAT condensates on supported lipid bilayers (SLBs) through addition of various T cell signaling proteins to membrane-attached phospho-LAT (pLAT) (Su et al., 2016). In separate work, we also reconstituted membrane-associated contractile actomyosin networks by attaching actin to SLBs via the membrane-anchored actin binding domain of ezrin (eABD) in the presence of myosin II and capping protein (Köster et al., 2016). To examine interactions of LAT condensates with actin networks, here we combined these two systems into a single assay (Figure 1A). We attached polyhistidine-tagged phospho-LAT (pLAT) and polyhistidine-tagged eABD to Ni-NTA functionalized lipids within the SLB. We induced LAT phase separation into condensates by adding an increasing subset of binding partners, in the order Grb2, Sos1, phospho-SLP-76 (pSLP-76), Nck, and, finally, N-WASP, as previously described (Su et al., 2016). Hereafter we use the nomenclature pLAT ➔ X to indicate condensates containing pLAT and all binding partners up to X (e.g. if X is Nck, then the condensates would contain pLAT, Grb2, Sos1, pSLP-76, and Nck). Note that in T cells, the main WASP family protein at the IS is WASP (Kumari et al., 2014), which acts as a constitutive complex with WASP Interacting Protein (WIP) (Anton et al., 2002; Ramesh et al., 1997). However, due to difficulties expressing recombinant full length WASP, here we used full length N-WASP fused N-terminally to the N-WASP binding fragment (residues 451-485) of WIP. WASP and N-WASP are 48 % identical in overall amino acid sequence, and 56 % identical in the regions that appear to be most important in the biochemical context of phase separation (Basic and Proline-rich, see below and Su et al. (Su et al., 2016)). Further, the known binding interactions and regulation of WASP and N-WASP are highly analogous (Padrick and Rosen, 2010). Thus, while different in an organismic context (Jain and Thanabalu, 2015; Snapper et al., 2001, 1998), the essential biochemical behaviors of WASP are very likely to be reflected in N-WASP. We fused N-WASP to the N-WASP binding fragment of WIP in order to stabilize the EVH1 domain of the protein (Peterson et al., 2007; Volkman et al., 2002), to create a single, well-expressed polypeptide. Our use of full length N-WASP here represents a step closer to the natural signaling system than our previous reconstitution, which used only a C-terminal fragment, enabling us to better capture essential features of cellular LAT condensates while still maintaining manageable complexity.

**Figure 1.**
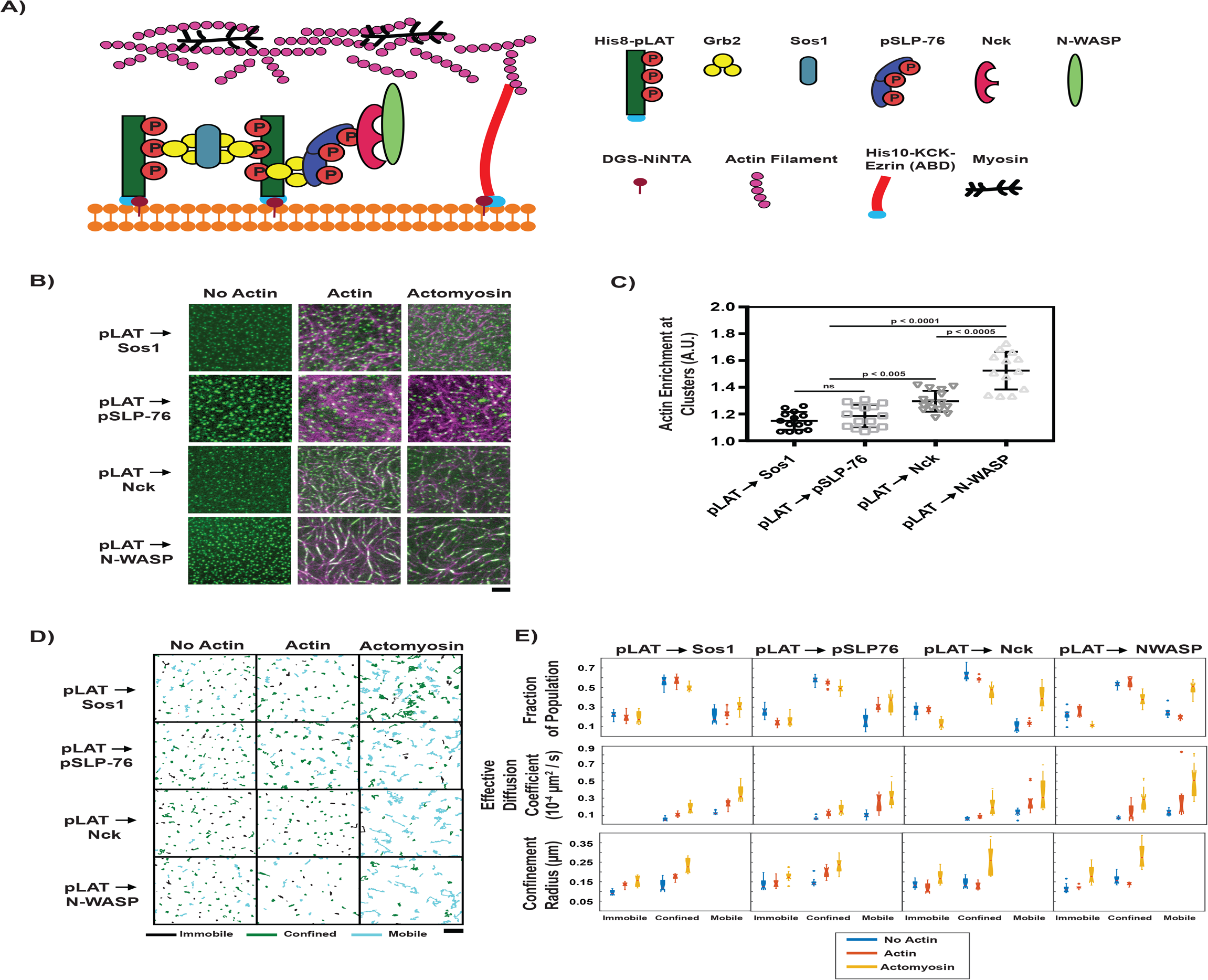
Composition regulates condensate movement with actin networks. **(A)** Schematic of biochemical reconstitution in solutions containing LAT and eABD attached to SLBs and LAT phase separation agents, actin filaments, and myosin filaments in solution. **(B)** TIRF microscopy images of LAT condensates of the indicated compositions (see text for nomenclature) without actin (left column), with actin-only networks (middle column), or with actomyosin networks (right column). pLAT-Alexa488 is green, rhodamine-actin is magenta. Scale bar = 5 μm. **(C)** Actin enrichment at condensates in reconstitution assays using actomyosin, the ratio of actin fluorescence intensity within condensates to actin intensity outside condensates (see Methods). Shown are the individual data points and their mean +/- s.d. from N = 15 fields of view from 3 independent experiments (with 5 FOV per experiment). P-values are for indicated distribution comparisons via Wilcoxon rank-sum test with Bonferroni correction. **(D)** Example plots of condensate tracks colored based on their movement type, as classified by Moment Scaling Spectrum analysis. Black = immobile, green = confined, and cyan = mobile. Scale bar = 5 μm. **(E)** (top) Fraction of the indicated condensate mobility types in the absence of actin (blue), in the presence an actin-only network (red), or in the presence of an active actomyosin network (gold). (middle) Effective diffusion coefficients of the condensates categorized and colored as in top row (not relevant for immobile condensates). (bottom) Confinement radius of the condensates categorized and colored as in top row (not relevant for mobile condensates). Each measurement is shown as a boxplot of the distribution of values from analyzing N = 15 fields of view from 3 independent experiments (with 5 FOV per experiment). Boxplot description: red central mark shows median; box edges show 25th and 75th percentiles; dashed whiskers extend to the most extreme data points not designated as “outliers”; and notch emanating from median indicates the 95% confidence interval around the median, shown for visual aid.

For experiments involving actin, we added polymerized actin filaments that bound to the SLB via anchored eABD. To induce actin filament movement we added muscle myosin II and ATP, as previously described (Köster et al., 2016). While T cells express only non-muscle myosin II, the muscle isoform is functionally similar, differing largely in making somewhat longer filaments (800 nm vs 300 nm average length under similar conditions (Vicente-Manzanares et al., 2009)), and is much easier to purify from tissues in biochemical quantities. Since most movement of T cell receptor condensates (which typically coincide with LAT condensates) is blocked by inhibition of myosin (Yi et al., 2012), more complex reconstitutions including actin filament assembly and disassembly dynamics were not warranted in our work here (Blanchoin et al., 2000; Didry et al., 1998; Shekhar and Carlier, 2017). Thus, our reconstituted system should retain key qualitative behaviors of T cell actomyosin, while remaining experimentally practical.

We induced LAT condensate formation without actin, with actin alone, or with active actomyosin networks (Figure 1B, Videos 1, 2, and 3). We immediately observed that condensates containing Nck or Nck and N-WASP associated with and wet actin filaments in both actin networks alone and actomyosin networks, while those lacking these proteins remained distributed across the SLB (Figure 1B). As a corollary, co-localization analysis (see Supplemental Methods) showed that actin enrichment in condensates increased significantly in the presence of Nck and N-WASP (Figure 1C).

In all conditions, we automatically detected and tracked the condensates at the SLB for 15 minutes, and then classified their movement on the SLB using Moment Scaling Spectrum analysis ((Jaqaman et al., 2011, 2008; Vega et al., 2018); see Materials and Methods) (Figure 1 – figure supplement 1). This analysis revealed that LAT condensates were either immobile, confined (i.e. moved within a confinement region), or mobile (i.e. moved without apparent restrictions, in a manner akin to free diffusion). In the absence of actin, 80% of condensates showed little movement, regardless of composition; they were either immobile or confined, in an area of radius 100-150 nm (Figure 1D, E, Video 1). In the presence of actin alone (i.e. no myosin), condensate movement varied with composition. pLAT ➔ Sos1 and pLAT ➔ pSLP76 showed an increase in apparent diffusion coefficient, the mobile fraction, and/or confinement radius (Figure 1D, E, Video 2), while pLAT ➔ N-WASP had a tendency to align with actin filaments and showed a decrease in the mobile fraction and confinement radius (Figure 1D, E, Video 2). pLAT ➔ Nck exhibited an intermediate behavior, with a slight increase in the fraction of mobile condensates and their apparent diffusion coefficient, but at the same time a decrease in the confinement radius of confined and immobile condensates (Figure 1D, E, Video 2). pLAT ➔ Nck also tended to align with actin filaments, although to a lesser degree than pLAT ➔ N-WASP (Figure 1B, C). Lastly, in the presence of active actomyosin networks, condensates of all compositions exhibited an overall increase in mobility (larger mobile fraction, apparent diffusion coefficient, and/or confinement radius). The increase for pLAT ➔ Sos1 and pLAT ➔ pSLP76 was subtle, larger for pLAT ➔ Nck, and largest for pLAT ➔ N-WASP. The influence of actomyosin on pLAT➔ N-WASP condensates was generally in the opposite direction of the influence of actin alone (Figure 1D, E, Video 3). These data suggest that condensates containing Nck or Nck and N-WASP, which wet filaments and show a large differential in behavior between actin alone and actomyosin conditions, can be viewed distinctly from condensates containing Sos1 or Sos1 and pSLP76, which do not wet filaments and show small differences between the two types of actin networks.

#### LAT condensates *in vitro* containing Nck and N-WASP move with contracting actomyosin networks with high fidelity

To delineate the effect of composition on the ability of LAT condensates to move with actin filaments, we devised an *in vitro* system where the actin filaments moved in a directional manner. This system enabled us to clearly distinguish between condensate that move with the actin filaments (because they would also exhibit directional movement) and those that do not. For this, we performed experiments where an SLB-associated actin network was induced to contract into asters by addition of myosin II filaments in the presence of low concentrations of salt and ATP (see Supplemental Methods). These experiments were especially susceptible to photo-damage caused by light exposure (Figure 2 – figure supplement 1), thus imaging conditions were set to minimize light exposure and the end point of all time-lapse images was compared with adjacent, non-imaged regions of the SLB. These actomyosin contraction experiments were performed with pLAT ➔ Sos1 and pLAT ➔ N-WASP condensates. In the beginning of such experiments, we found that pLAT ➔ Sos1 condensates were randomly distributed on the membrane while pLAT ➔ N-WASP condensates were aligned along the actin filaments as observed above (Figure 2A). In both cases the filament network started to contract immediately upon myosin II addition and formed stable asters within 2 minutes (Video 4). As shown in Figure 2A, at the end of the contraction, most of the pLAT ➔ Sos1 condensates remained scattered across the SLB, while virtually all of the pLAT ➔ N-WASP condensates had moved with the actin into asters. To quantify these behaviors, we examined the speed and direction of condensate movement and actin movement during actomyosin network contraction using Spatio-Temporal Image Correlation Spectroscopy (STICS) (Ashdown et al., 2014) (Figure 2B). We found that the speed of pLAT ➔ N-WASP condensates correlated well with the speed of actin, while the speed of pLAT ➔ Sos1 condensates did not (Figure 2C). Additionally, the distribution of angles between the vectors of pLAT ➔ N-WASP condensate movement and proximal actin movement showed clear preference for smaller angles, indicating a high degree of co-movement. In contrast, the angle distribution for pLAT ➔ Sos1 showed only a slight preference for smaller angles, which was marginally significant (Figure 2D). Together, the steady state (Figure 1) and contraction (Figure 2) experiments and analyses show that LAT condensates are influenced by actin network dynamics in a composition-dependent fashion. Condensates containing Nck or N-WASP bind to and move with actomyosin filaments. In contrast, condensates lacking these proteins do not bind filaments appreciably, and are likely moved by non-specific steric contacts. Thus, Nck and N-WASP function as a molecular clutch between LAT condensates and actin.

**Figure 2.**
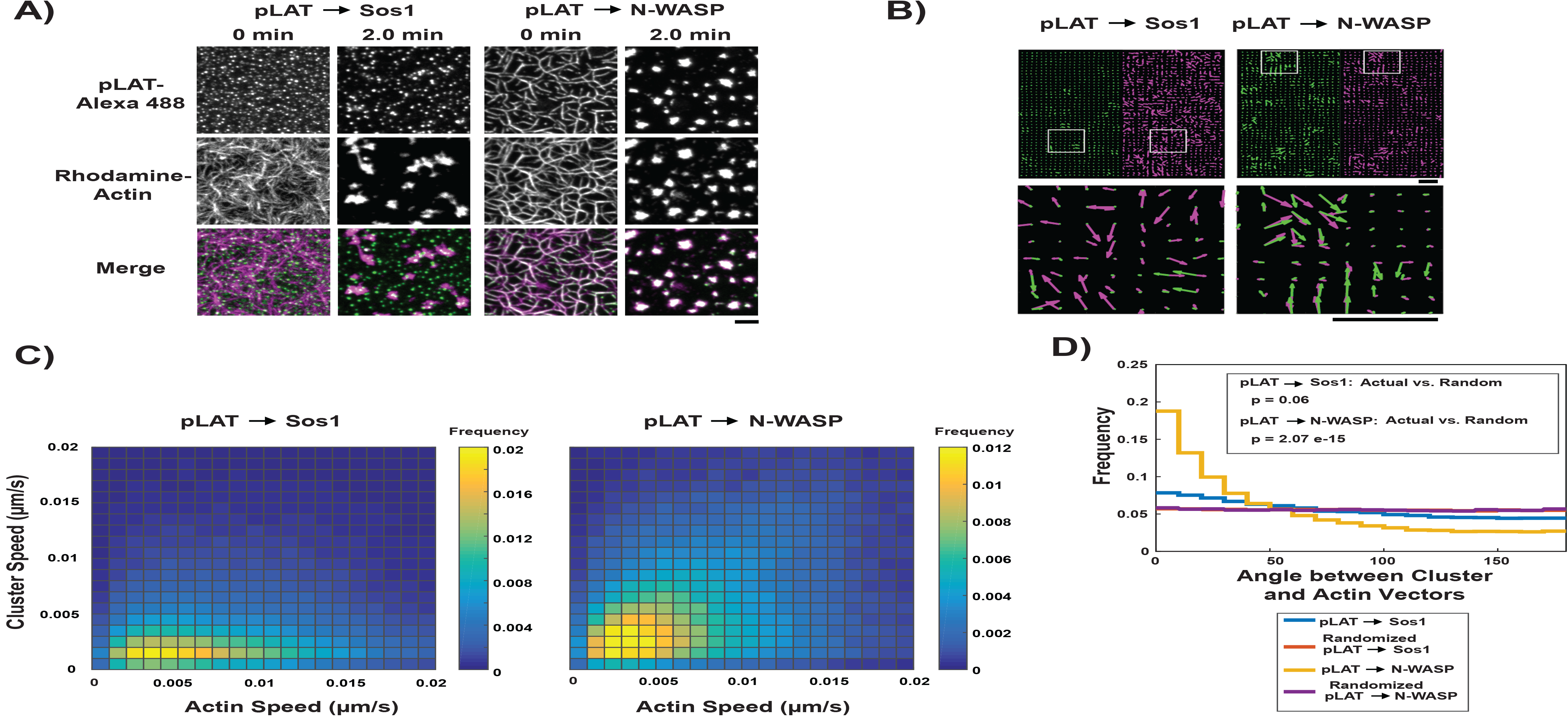
pLAT ➔ N-WASP condensates bind to and move with moving actin filaments. **(A)** TIRF microscopy images of pLAT ➔ Sos1 condensates (left two columns) and pLAT ➔ N-WASP condensates (right two columns) formed in an actin network before (t = 0 min) and after (t = 2 min) addition of myosin II. pLAT condensates are green and actin is magenta in merge. Scale bar = 5 μm. **(B-D)** STICS analysis of actin and condensate movement. **(B)** Representative map of actin (magenta) and pLAT condensate (green) vector fields. Lower panels show magnification of box regions in upper panels. **(C)** Condensate speed vs. actin speed at same position. Condensate composition indicated above each heat map. Heat map indicates frequency in each bin, i.e. counts in each bin normalized by total number of counts. **(D)** Distribution of the angle between actin and condensate movement vectors for pLAT ➔ Sos1 (blue), pLAT ➔ N-WASP (gold), randomized pLAT ➔ Sos1 (red) and randomized pLAT ➔ N-WASP (purple) (see Methods for randomization). P-values are for indicated distribution comparisons via Kolmogorov-Smirnov test. Data in (C) and (D) are pooled from 15 fields of view from 3 independent experiments (5 FOV per experiment).

### Basic regions of Nck and N-WASP couple LAT condensates to actin filaments

The data above suggest that Nck and N-WASP mediate binding of LAT condensates to actin filaments. To test this, we quantified the recruitment of preformed actin filaments to SLBs by LAT condensates of different compositions in the absence of eABD. As shown in Figure 3A, only condensates containing Nck or Nck and N-WASP recruited substantial amounts of actin filaments. In both cases, condensates were deformed and elongated along filaments, as observed above.

**Figure 3.**
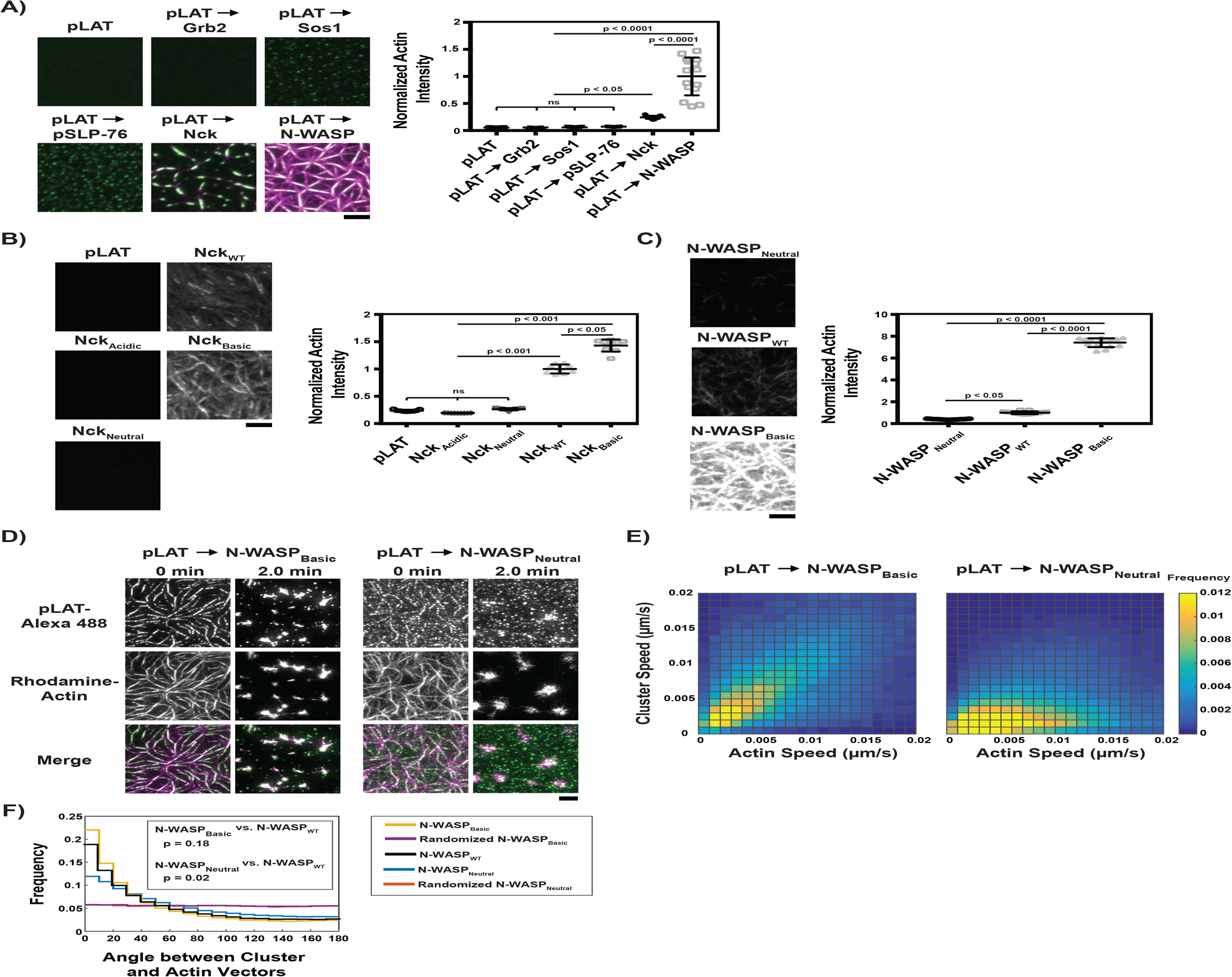
Basic regions of Nck and N-WASP mediate interaction of LAT condensates with actin filaments. **(A)** (Left) TIRF microscopy images of rhodamine-actin (magenta) recruited to SLBs by the indicated LAT condensate compositions (green). Scale bar = 5 μm. All panels use the same intensity range. (Right) Normalized average rhodamine-actin fluorescence intensity on SLBs. Shown are the individual data points and their mean +/- s.d. N = 15 fields of view from 3 independent experiments (5 FOV per experiment). **(B)** (Left) TIRF microscopy images of rhodamine-actin recruited to SLBs by His-tagged Nck variants. Scale bar = 5 μm. All panels use the same intensity range. (Right) Normalized average rhodamine-actin fluorescence intensity on SLBs. Shown are the individual data points and their mean +/- s.d. N = 15 fields of view from 3 independent experiments (5 FOV per experiment). **(C)** (Left) TIRF microscopy images of rhodamine-actin recruited to SLBs by His-tagged N-WASP variants. All panels use the same intensity range. (Right) Normalized average rhodamine-actin fluorescence intensity on SLBs. Shown are the individual data points and their mean +/- s.d. N = 15 fields of view from 3 independent experiments (5 FOV per experiment). **(D)** (Left) TIRF microscopy images of pLAT ➔ N-WASP_Basic_ condensates (left two columns) and pLAT ➔ N-WASP_Neutral_ condensates (right two columns) formed in an actin network before (t = 0 min) and after (t = 2 min) addition of myosin II. LAT condensates are green and actin is magenta in merge. Scale bar = 5 μm. **(E)** Condensate speed vs. actin speed at same position from STICS analysis. Condensate composition indicated above each heat map. Heat map indicates frequency in each bin (as in Figure 2C). N = 15 fields of view from 3 independent experiments (5 FOV per experiments). **(F)** Distribution of the angle between actin and condensate movement vectors for pLAT ➔ N-WASP_Basic_ (gold), randomized pLAT ➔ N-WASP_Basic_ (purple), pLAT ➔ N-WASP_WT_ (black, same data as in Figure 2), pLAT ➔ N-WASP_Neutral_ (blue), and randomized pLAT ➔ N-WASP_Neutral_ (red). N = 15 fields of view from 3 independent experiments (5 FOV per experiment). P-values are for indicated distribution comparisons via Kolmogorov-Smirnov test.

We next performed co-sedimentation assays to determine whether Nck or N-WASP could bind actin filaments in solution or whether efficient binding required the proteins to be arrayed on a two-dimensional surface. Nck did not appreciably co-sediment with actin filaments. Consistent with previous reports, N-WASP did bind filaments (Co et al., 2007), although not to the same degree as α-actinin, a high-affinity actin filament-binding protein (Figure 3 – figure supplement 1). To identify the elements of Nck that mediate LAT condensate binding to actin filaments, we attached polyhistidine-tagged fragments of Nck to SLBs and measured their ability to recruit actin filaments from solution. Nck is composed of three SH3 domains and an SH2 domain, connected by flexible linkers of 25-42 residues (Banjade et al., 2015). Of these seven elements, two contain dense basic patches, one contains dense acidic patches, and the remainder are relatively free of dense charge patches. As detailed in Figure 3 – figure supplement 2, Nck fragments containing an excess of basic elements recruited actin to the membrane, where greater excess resulted in more efficient recruitment, while neutral or acidic fragments did not. Similarly, mutating one of the basic elements (the linker between the first and second SH3 domains, L1) to neutralize it (Nck_Neutral_) or to make it acidic (Nck_Acidic_) greatly impaired actin recruitment, while making it more basic (Nck_Basic_) enhanced actin recruitment (Figure 3B, Figure 3 – figure supplement 3). These data indicate that basic regions of Nck likely contribute to binding of LAT condensates to actin filaments.

Like Nck, N-WASP also has a central basic region (amino acid residues 186-200). We thus asked whether this region and/or the two C-terminal WH2 motifs contribute to the coupling of LAT condensates to actin filaments (Bieling et al., 2018; Co et al., 2007). We generated pLAT ➔ N-WASP condensates with N-WASP fragments consisting of the basic-proline elements (BP), basic-proline + VCA (BPVCA) elements, and basic-proline + VCA_mut_ (BPVCA_mut_), which contains mutations to the WH2 motifs in the VCA region that impair filament binding (Co et al., 2007). All three types of condensates strongly recruited actin, indicating that WH2-actin interactions are not needed for actin filament recruitment in the context of LAT condensates (Figure 3 – figure supplement 4). To further examine the basic region, we generated three variants of His_6_-tagged full length N-WASP (fused N-terminally to the WASP-binding region of WIP to afford stability; Figure 3 – figure supplement 4): N-WASP containing a doubled basic region (N-WASP_Basic_), wild-type N-WASP (N-WASP_WT_) and N-WASP containing a neutral linker instead of the basic region (N-WASP_Neutral_). As shown in Figure 3C and Figure 3 – figure supplement 5, these variants recruited actin in the order N-WASP_Basic_ >> N-WASP_WT_ > N-WASP_Neutral_. Thus, for both Nck and N-WASP, the degree of positive charge in basic elements strongly correlates with the ability of the proteins and their LAT condensates to recruit actin filaments to SLBs.

To test whether these basic region-mediated interactions are necessary to couple LAT condensate movement to actin movement (Figure 2), we performed actin contraction assays with condensates containing N-WASP_Basic_ or N-WASP_Neutral_. Before myosin II addition, pLAT-> N-WASP_Basic_ condensates aligned almost perfectly with actin filaments, while pLAT-> N-WASP_Neutral_ condensates only partially aligned with actin filaments, consistent with the notion that the basic region mediates binding to actin filaments (Figure 3D). After actin network contraction, pLAT ➔ N-WASP_Basic_ condensates colocalized with actin asters to a similar degree as pLAT ➔ N-WASP_WT_, while pLAT ➔ N-WASP_Neutral_ did not (Figure 3D vs. Figure 2A, Video 5 vs. Video 4). STICS analysis revealed that the correlation of pLAT ➔ N-WASP_Basic_ movement with local actin movement was slightly better than that of pLAT à N-WASP_WT_, while the correlation of pLAT ➔ N-WASP_Neutral_ was worse (Figure 3E, F). The remaining correlation for pLAT ➔ N-WASP_Neutral_ is most likely due to the presence of Nck, which also contributes to condensate binding to actin. Together these data demonstrate that regions of Nck and N-WASP that contain dense basic patches can mediate the clutch-like behaviors of the proteins by directly interacting with actin filaments proportionally to the degree of positive charge, and that these interactions are necessary for LAT condensates to faithfully move with actin.

### The composition of LAT condensates changes as they traverse the IS

In Jurkat T cells activated by OKT3, an antibody that binds to the CD3ε subunit of the TCR, and ICAM-1 bound to an SLB, LAT condensates that form at the periphery of the IS move to its center over ~5 minutes as the cell-SLB contact matures. To investigate whether the composition-dependent interactions observed in our biochemical data have consequences for LAT condensate behavior in cells, we examined the composition of LAT condensates as they moved in the plane of the plasma membrane during activation of live Jurkat T cells. We used Jurkat T cells because they retain many features of primary T cells relevant to movement and signaling from LAT condensates, but are easier to manipulate and analyze. In both cell types, LAT condensate formation is well-documented (Balagopalan et al., 2013; Lin et al., 1999; Su et al., 2016; Yokosuka et al., 2005), condensate movement across the IS is correlated with actin flow (DeMond et al., 2008; Kaizuka et al., 2007; Murugesan et al., 2016; Yi et al., 2012), and proximal biochemical signaling from LAT through SLP-76 is similar (Bartelt et al., 2009). However, the IS between Jurkat T cells and supported lipid bilayers is larger than that of primary T cells (Murugesan et al., 2016), and LAT condensates do not initiate actin polymerization to the same degree in Jurkat T cells as in primary T cells, which allows us to analyze the ability of condensates to couple to dynamic actin networks without accounting for their own self-generated polymerized actin (Kumari et al., 2015).

We co-expressed LAT-mCitrine with Grb2-mCherry or LAT-mCherry with Nck-sfGFP in Jurkat T cells. These cells bound to SLBs coated with mobile ICAM-1 and OKT3 producing an IS mimic with the SLB. Of note, in contrast to Jurkat T cells adhered to immobile substrates, which extend long lamellipodia (Babich et al., 2012), Jurkat T cells adhering to fluid SLBs here extend short lamellipodia at the IS periphery. We used total internal reflection fluorescence (TIRF) microscopy to capture images of activated cells every 5 seconds for up to 5 minutes. We then automatically detected and tracked LAT condensates from their formation at the periphery of the IS to their coalescence with the central supramolecular activation complex (cSMAC) at the synapse center (Jaqaman et al., 2008) (Figure 4 – figure supplement 1A, Figure 4 – figure supplement 2, Video 6), monitoring the fluorescence intensity of LAT and Grb2 or Nck in the condensates. In order to follow the LAT condensates neutrally, i.e. while blind to their Grb2 or Nck content, we detected and tracked them based on the LAT channel, and then read out the Grb2 or Nck intensities at the LAT condensate locations (“master/slave” channel analysis, as in (Loerke et al., 2011)). In addition to affording neutrality, this scheme ensured accurate LAT condensate detection and tracking because of the overall better signal in the LAT channel, especially when compared to the Nck channel, which tended to have high background due to soluble Nck molecules not bound to LAT condensates. Furthermore, to overcome the stochasticity and noise inherent to live-cell image data, we aligned tracks (in space or time, as described below) and averaged them to uncover underlying overall trends. For meaningful alignment and averaging, we filtered tracks by their duration, extent of directed movement, initial and final positions (to ensure that they traversed a sufficient radial distance across the IS), and initial Grb2 or Nck intensity to ensure that changes in intensity could be measured accurately (see Materials and Methods for more details).

This analysis revealed that Grb2 colocalized with LAT condensates at the edge of the synapse (Figure 4A, Video 7) and its fluorescence intensity in the condensates was maintained throughout the trajectory to the center of the synapse (Figure 4B, C). In contrast, while Nck also colocalized with LAT condensates at the edge of the synapse (Figure 4D, Video 8), its fluorescence diminished relative to LAT during the trajectory. This intensity decrease began at 0.6-0.7 of the distance from the center of the synapse to the synapse edge (referred to hereafter as “normalized radial position,” equal to zero at the synapse center and one at the synapse edge; see Materials and Methods) (Figure 4C, E). The emergence of this pattern from averaging 125 tracks from 25 cells suggests that spatial position is a key determinant of Nck residence in LAT condensates. In contrast, when the condensates were aligned according to their appearance time (Figure 4 – figure supplement 1B), the median Nck intensity steadily decreased over the course of the trajectory across the IS without any characteristic start time (beyond t = 0). The decrease in Nck intensity is not the result of photobleaching (Figure 4 – figure supplement 3). This suggests that in this experimental setting, position plays a more instructive role than time in determining the residence of Nck in condensates (and presumably the residence of WASP, which is recruited to LAT condensates via Nck). However, since time and space are coupled, our data do not rule out a role for time in this process, as has previously been observed in experimental conditions where LAT condensates were immobilized (Barda-Saad et al., 2005).

**Figure 4.**
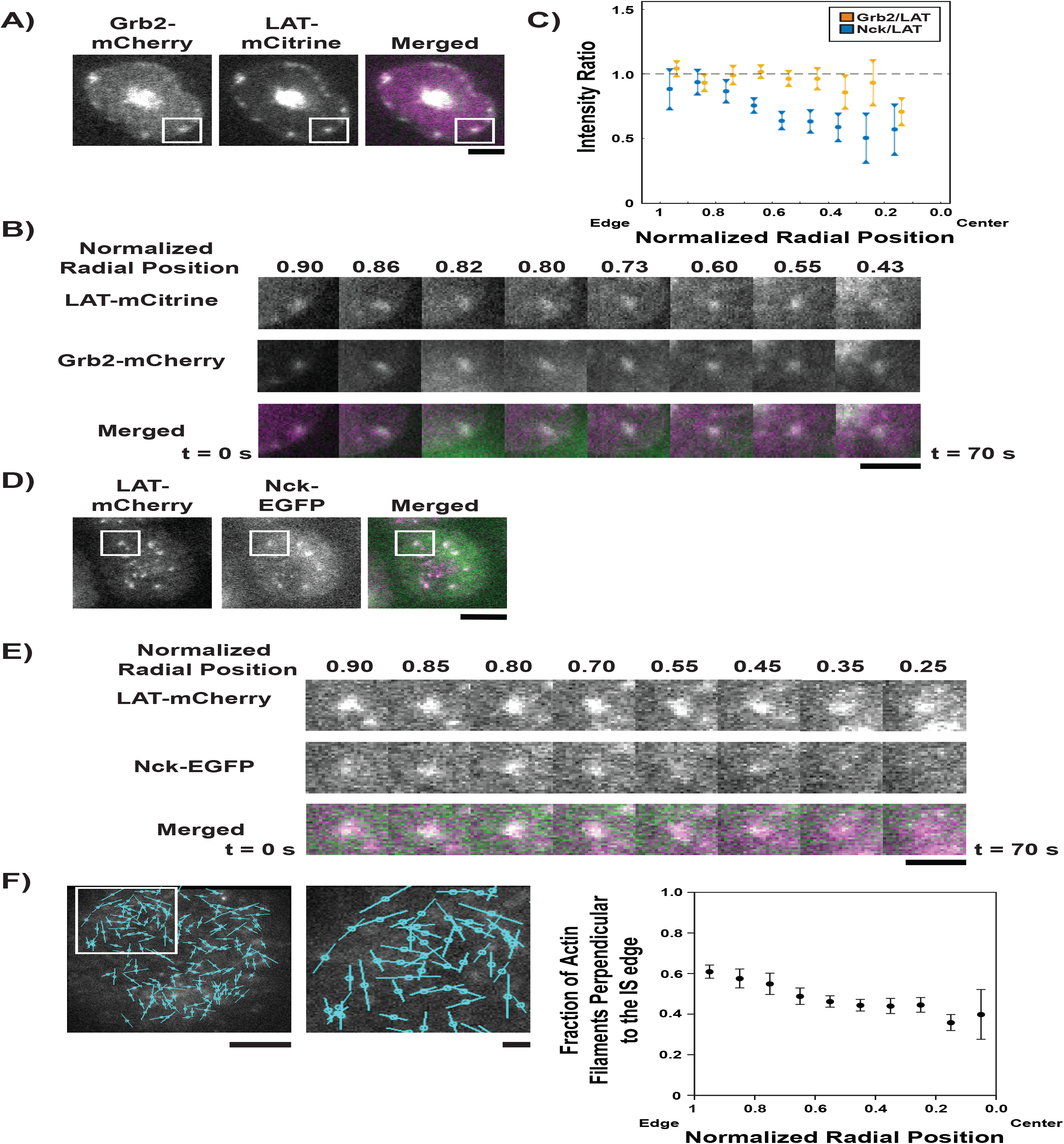
LAT cluster composition changes as clusters move across the IS.

### Nck dissipation coincides in space with the switch in actin architecture from dendritic network to concentric arcs

Translocation of LAT condensates from the synapse periphery to the center of the IS is driven by motion of the actin cytoskeleton (DeMond et al., 2008; Kaizuka et al., 2007; Mossman et al., 2005; Yu et al., 2010). Recent work has shown that two actin networks are generated at the IS in activated T cells (Murugesan et al., 2016; Yi et al., 2012). In the outer ~1/3 of the synapse, the Arp2/3 complex generates a dendritic actin meshwork, where the filaments are on average directed radially, perpendicular to the synapse edge. In the medial region closer to the cSMAC, this meshwork is largely replaced by formin-generated concentric actin arcs that are directed parallel to the synapse edge (Figure 4 – figure supplement 4) (Hammer 3rd and Burkhardt, 2013; Yi et al., 2012). Both filament networks move through the action of myosin motors as the cell-cell conjugate matures; however, the nature of this movement is different in the two cases. The outer dendritic network moves in a direction perpendicular to the edge of the synapse in a process termed retrograde flow (DeMond et al., 2008; Kaizuka et al., 2007; Mossman et al., 2005; Yu et al., 2010), analogous to actin flow observed at the leading edge of migrating cells (Ponti et al., 2005, 2004). In contrast, the inner concentric arcs sweep toward the center of the synapse in a telescoping manner and appear to have components of motion both perpendicular and parallel to the synapse edge (Murugesan et al., 2016).

The normalized radial position at which LAT condensates started to lose Nck is similar to the position where the actin network has been reported to change from dendritic architecture to arc architecture (Hammer 3rd and Burkhardt, 2013; Murugesan et al., 2016; Yi et al., 2012). To corroborate this in our cells, we incubated Jurkat T cells with the dye SiR-Actin, which binds to actin filaments in a defined orientation (perpendicular to the filament orientation; Figure 4 – figure supplement 5) (Nordenfelt et al., 2017). This dye enabled us to use instantaneous polarization TIRF microscopy (Mehta et al., 2016) to evaluate the orientation of actin filaments at the IS. We found that SiR-actin at concentrations higher than 50nM blocked actin flow at the IS, but speckle labeling of actin networks with 10 nM SiR-actin produced normal actin flow toward the center of the IS. Since the actin networks were stained as sparsely distributed speckles of SiR-actin, our analysis procedure does not involve (or require) identification of individual filaments in the images and then measuring their orientations. Rather, by observing the fluorescence polarization orientation of single speckles randomly bound to actin filaments, we can build a spatial map of filament orientations across the IS. We found that actin filaments in the outer 30% of the synapse were generally oriented perpendicular to the synapse edge, while those closer to the center of the synapse were parallel to the synapse edge (Figure 4F), in good agreement with earlier super-resolution work (Murugesan et al., 2016). The change in actin filament polarization occurred at 0.6-0.7 of the distance from the center of the synapse, which correlates well with the position at which Nck dissipated from LAT condensates. This position is unrelated to the overall three-dimensional geometry of the cell, indicating that actin architecture at the IS, as observed via TIRF microscopy, is not determined by the position of the cell body above the membrane surface (Figure 4 – figure supplement 6).

### Constitutive engagement of LAT condensates with actin leads to their aberrant movement across the IS

Our combined biochemical and cellular data thus far indicate that Nck and WASP mediate LAT condensate engagement with actin, and that LAT condensates lose these proteins as they move from the dendritic actin meshwork in the outer part of the synapse to the contractile arcs network closer to the synapse center. The biochemical data suggest that this change in composition should allow LAT condensates to interact differently with the two actin networks. We hypothesized that this switch in interaction might be necessary for the proper radial movement of LAT condensates at the IS, given the different orientation (perpendicular vs. parallel to the synapse edge) and movement (retrograde flow vs. telescoping motion) of filaments in the two actin networks at the IS. To test this hypothesis, we altered the adhesion of LAT condensates to the actin filament network by fusing Grb2, which remains in the condensates throughout their trajectories (Figure 4A-C), with the doubled basic region of N-WASP (Grb2_Basic_). In biochemical assays, this generated LAT condensates that bound actin filaments in the absence of Nck or WASP. The pLAT-Grb2_basic_ complex recruited actin filaments to SLBs, while pLAT alone or the pLAT-Grb2 complex did not (Figure 5A). In actomyosin contraction assays, condensates of pLAT/Grb2_Basic_/Sos1 initially wet filaments and then localized to actin asters after myosin II-induced contraction to a greater degree than pLAT/Grb2/Sos1, although to a lesser degree than pLAT ➔ N-WASP (Figure 5B, Video 9 vs. Video 4). Similarly, during actomyosin network contraction, the movement of pLAT/Grb2_Basic_/Sos1 condensates was correlated more strongly with actin movement than condensates containing Grb2 (Figure 5 – figure supplement 1), but less strongly than pLAT ➔ N-WASP condensates (Figure 2C, D). Together, these data demonstrate that the double basic motif of N-WASP, when added to Grb2, can act as a molecular clutch coupling LAT condensates to actin.

**Figure 5.**
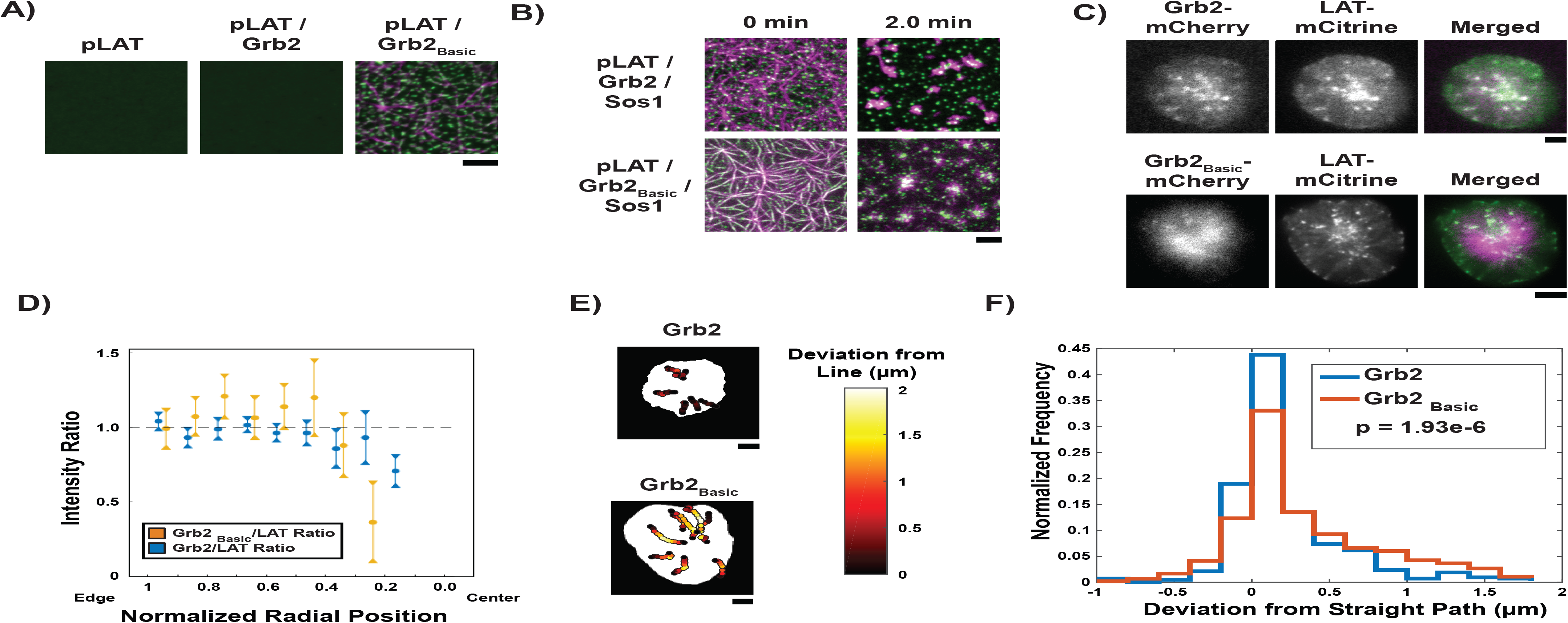
Grb2 fused to a basic molecular clutch can couple LAT condensates to actin. **(A)** TIRF microscopy images of rhodamine-actin recruited to SLBs by His-tagged pLAT or condensates of pLAT ➔ Grb2 or pLAT ➔ Grb2_Basic_. Scale bar = 5 μm. **(B)** TIRF microscopy images of pLAT ➔ Sos1 condensates containing Grb2_WT_ (top row; data from Figure 2) or Grb2_Basic_ (bottom row) formed in an actin network before (t = 0 min) and after (t = 2 min) addition of myosin II. Actin shown in magenta and LAT condensates in green. Scale bar = 5 μm. **(C)** TIRF microscopy image of Jurkat T cell expressing Grb2-mCherry (top, magenta in merge) or Grb2_Basic_-mCherry (bottom, magenta in merge) and LAT-mCitrine (green in merge) activated on an SLB coated with OKT3 and ICAM-1. Scale Bar = 5 μm. **(D)** Fluorescence intensity ratios of Grb2 / LAT (blue, data from Figure 4C) or Grb2_Basic_ / LAT in condensates (gold) at different normalized radial positions. Measurements were made at identical relative locations but data are slightly offset in the graph for visual clarity. Plot displays median and notches from boxplot for 95% confidence interval from N = 44 condensates from 12 cells expressing Grb2_Basic_-mCherry and LAT-mCitrine from 7 independent experiments and 82 condensates from 11 cells expressing Grb2-mCherry and LAT-mCitrine from 5 independent experiments (same cells as in Figure 4C). Only tracks in which the mean Grb2 or Grb2_Basic_ intensity was greater than 1 standard deviation above background during the first three measurements were used to generate this plot. **(E)** Trajectories of LAT condensates in Jurkat T cells expressing Grb2-mCherry (top) or Grb2_Basic_-mCherry (bottom) recorded over 2 to 5 minutes of imaging. Trajectories are color-coded as indicated in the legend at right according to deviation from a straight line between the estimated starting point of actin engagement and just before entering the cSMAC (see Methods). Scale bar = 5 μm. **(F)** Distribution of deviations from a straight line for condensates in Jurkat T cells expressing Grb2-mCherry (red) or Grb2_Basic_-mCherry (blue). N = 44 condensates from 12 cells expressing Grb2_Basic_-mCherry and LAT-mCitrine from 7 independent experiments and 82 condensates from 11 cells expressing Grb2-mCherry and LAT-mCitrine from independent experiments (same cells as in Figure 4C). P-value is for comparing the two distributions via a Kolmogorov-Smirnov test. Only tracks in which the mean Grb2 or Grb2_Basic_ intensity was greater than 1 standard deviation above background during the first three measurements were used to generate this plot.

We next asked whether expression of Grb2_Basic_ in Jurkat T cells would perturb the radial movement of LAT condensates due to their constitutively engaged clutch, including in the medial region of the IS where they encounter actin arcs. This was quantified as deviation from a straight path between the start of persistent inward radial movement and coalescence with the cSMAC (see Supplemental Methods for more details). For this, cells expressing Grb2_Basic_-mCherry were activated on SLBs as above. Similar to cells expressing Grb2-mCherry, LAT condensates that formed at the periphery of the IS retained Grb2_Basic_-mCherry throughout their trajectories to the cSMAC (Figure 5C, D). However, evaluation of condensate trajectories revealed that they deviated from a straight path significantly more than condensates in cells expressing Grb2-mCherry (Figure 5E, F, Video 10 vs. Video 7). This behavior is consistent with abnormally high adhesion of condensates containing Grb2_Basic_ to actin filaments, even after Nck has presumably dissipated, leading to trajectories that reflected more the telescoping, circular component of the contractile actin arc motion.

### Formin activity is necessary for LAT condensate composition change

Finally, we asked whether the transition from the dendritic actin architecture to the contractile arc architecture might play a role in changing the composition of LAT condensates at this location in the IS. Previous data showed that the contractile arcs are generated by the formin mDia1 and could be eliminated by cell treatment with the formin inhibitor, SMIFH2 (Murugesan et al., 2016). We found that in contrast to control cells (treated with DMSO), where Nck dissipated normally from LAT condensates (Figure 6A and B, Figure 6 – figure supplement 1, Video 11), cells treated with SMIFH2 for five minutes prior to imaging displayed LAT condensates with virtually constant Nck intensities throughout their trajectories from the periphery to the cSMAC (Figure 6A and C, Figure 6 – figure supplement 1, Video 12). Thus, the activity of formin proteins, and/or perhaps the actin arcs that they generate, act to alter the composition of LAT condensates, likely altering their downstream signaling activities in the central region of the IS. We note that the SMIFH2 data further support the notion that in unperturbed cells, space, rather than time, is the key determinant of Nck residence in condensates (assuming that formin does not also create a temporal signal). Our combined data suggest that the two actin networks in activated Jurkat T cells not only spatially organize the immunological synapse by moving LAT condensates, but may also contribute to creation of specific signaling zones.

**Figure 6.**
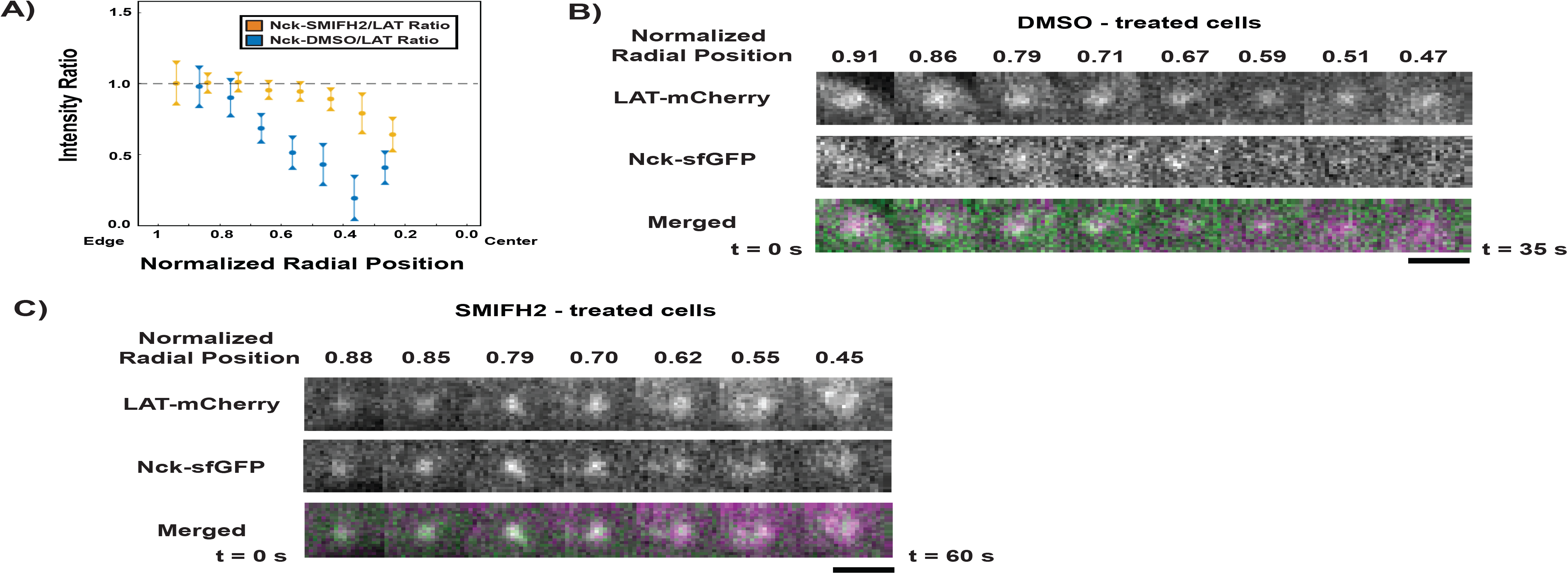
Formin activity is necessary for Nck dissipation from LAT condensates. **(A)** Fluorescence intensity ratios of Nck / LAT in condensates in Jurkat T cells treated with DMSO (blue) or the formin inhibitor, SMIFH2, (gold) at different normalized radial positions. Measurements were made at identical relative locations but data are slightly offset in the graph for visual clarity. Plot displays median and notches from boxplot for 95% confidence interval from N = 43 condensates from 11 DMSO-treated cells from 5 individual experiments and 102 condensates from 14 SMIFH2-treated cells from 5 individual experiments. Only tracks in which the mean Nck intensity was greater than 1 standard deviation above background during the first three measurements were used to generate this plot. The first DMSO data point (Normalized Radial Position = 1) does not appear in the plot because the number of detected condensates was too small (< 10) to generate a statistically meaningful measurement. **(B, C)** Magnification of boxed regions from Figure 6 – figure supplement 1 of condensates containing LAT-mCherry (magenta in merge) and Nck-sfGFP (green in merge) during their trajectories across the IS in a cell treated with DMSO **(B)** or SMIFH2 **(C)**. Normalized radial position indicated above image panels and time below panels. Scale bar = 2 μm.

## Discussion

Compositional changes of biomolecular condensates in response to signals have been well documented (Chen et al., 2008; Dellaire et al., 2006; Markmiller et al., 2018; Salsman et al., 2017; Youn et al., 2018). However, the functional consequences of these compositional changes have generally not been elucidated. Here, we show that compositional changes alter the interactions of LAT condensates on membranes with active actomyosin networks. Our *in vitro* data demonstrate that Nck and N-WASP act as a clutch that mediates strong LAT condensate binding to actin filaments, thus coupling LAT condensate movement to actomyosin filament movement. LAT condensates lacking Nck and N-WASP can still be propelled by steric interactions with moving actomyosin filaments, but less efficiently than when the clutch is present. We also show that the composition of LAT condensates in activated Jurkat T cells changes as they move radially from the edge to the center of the IS. This change spatially parallels the change from the peripheral dendritic actin network to the circular actin arcs, and inhibition of the formin mDia1 prevents loss of Nck from LAT condensates. Mutations that add a basic sequence to Grb2, thus constitutively engaging LAT condensates with actin filaments, cause aberrant movement of the condensates with a higher tendency to deviate from a straight line.

These data suggest a model for movement of LAT condensates across the IS by navigating the distinct cortical actin networks. Condensates form in the outer region of the IS with the full complement of signaling molecules, including Grb2, Sos1, SLP-76, Nck, and WASP. Basic regions on Nck and WASP act as a molecular clutch, enabling the condensates to adhere tightly to the outer dendritic actin network, and to travel radially with the network as it moves by retrograde flow. Near the junction with the formin-generated actin arcs, a formin-dependent signal causes loss of Nck, and likely WASP (which is present in condensates through its interactions with Nck). We do not yet know the nature of this signal, but one possibility would be dephosphorylation of SLP-76, as Nck is known to join condensates primarily through binding SLP-76 phosphotyrosines (Barda-Saad et al., 2010; Pauker et al., 2012). More complex mechanisms involving appearance of competing Nck binding partners or mechanical disruptions are also possible. Loss of Nck/WASP decreases adhesion of the LAT condensates to actin. In this state, it appears that the condensates can still be moved by actin, but likely through repeated non-specific steric interactions. This is consistent with previous observations that in the actin arcs region of the IS, LAT condensates are repeatedly hit by arcs that move them briefly but then release (Murugesan et al., 2016). The circular movement of the telescoping actin arcs is randomly directed clockwise and counterclockwise, and repeated hits by arcs moving oppositely produce no net circular motion on the condensates. But the radial component is consistently directed toward the center of the IS, and thus repeated hits add constructively to produce a net movement in a radial direction. Thus, LAT condensates continue to move linearly toward the center of the IS in the arc region. If the condensates were to adhere tightly to actin in the arcs region (i.e. if they contained Nck and WASP there), they would no longer undergo repeated hits by arcs moving in opposite directions, and the circumferential force would not average to zero. In the simplest case, they would attach to the first arc they encountered and move circumferentially with it. Such effects likely account for the aberrant movement of LAT condensates containing the Grb2 mutant artificially equipped with a basic clutch, which should produce inappropriately strong adhesion to the telescoping actin arcs.

While we have examined Jurkat T cells here, existing data suggest that the mechanisms we have discovered for LAT condensate movement are likely similar to those in primary T cells. The IS formed by both Jurkat T cells and primary T cells is composed of a dendritically-branched actin network at the edge of the IS, followed immediately by a concentric actomyosin cable network near the center of the IS (Murugesan et al., 2016). Thus, condensates must be moved across two distinct actin networks in Jurkat and primary T cells. In primary T cells, condensate-associated actin polymerization localizes mostly (although not entirely) in the outer region of the IS and dissipates in the region adjacent to the branched actin network, where ICAM-1 localizes (Kumari et al., 2015), which would be consistent with loss of Nck and WASP toward the center of the IS. It is tempting to speculate that the engagement of the clutch mechanism has mechano-signaling consequences, since T cell receptor signaling appears to be mechanically gated (Chen and Zhu, 2013; Hui et al., 2014). Similarly, PLC-γ1 activation by WASP-promoted actin polymerization appears to localize mostly in the dSMAC where we observe strong Nck co-localization with LAT (Kumari et al., 2015). One difference between the two cell types is that WASP-promoted actin polymerization is much weaker in Jurkat T cells than in primary T cells (Kumari et al., 2015). In primary cells, this actin assembly may also play a role in the movement of condensates from the cell edge to the center, in addition to myosin-driven movement of the cortical actin. Future work addressing the modes of movement, and the precise signals that dictate compositional change, will elucidate the mechanisms by which LAT condensates move across the IS in primary T cells.

LAT condensates represent one particular type of biomolecular condensate. It is generally thought that the functions of condensates are intimately connected to their compositions, and that changes in composition could cause changes in function (Banani et al., 2016). Our data here demonstrate that when Nck and N-WASP are arrayed on membranes they can bind actin filaments efficiently, even though both bind filament sides only weakly in solution. This adhesion enables condensates containing the proteins to be moved over long distances in response to actomyosin contraction. Adhesion is lost when Nck and N-WASP depart. Thus, the composition of LAT condensates plays an important role in their coupling to actin and their mode of movement at the IS. These behaviors of the LAT system are produced by generalizable features of membrane-associated condensates - their high density and composition based on regulatable interactions. Analogous behaviors are likely to be widely observed as the biochemical and cellular activities of other condensates are explored.

## Materials and Methods

### Protein Reagents

Human Ezrin (aa 477-586) with an N-terminal His_10_-KCK tag, human Grb2 (aa 1-217), human Grb2-Double Basic (aa 1-217 of Grb2 fused with the human N-WASP basic region (x2) KEKKKGKAKKKRLTKGKEKKKGKAKKKRITK), human LAT (aa 48-233) with an N-terminal His_8_ tag, human Nck1 (aa 1-377), human Nck (FL) (aa 1-377) with an N-terminal His_8_-C(GGS)_4_ tag, human Nck (FLΔL1) with an N-terminal His_8_-C(GGS)_4_ tag, human Nck (L1-S2-L2-S3-L3-SH2) with an N-terminal His_8_-C(GGS)_4_ tag, human Nck (L1(K to E)-S2-L2-S3-L3-SH2) with an N-terminal His_8_-C(GGS)_4_ tag, human Nck (L1(HM)-S2-L2-S3-L3-SH2) with an N-terminal His_8_-C(GGS)_4_ tag, human Nck (S1-L1-S3-L3-SH2) with an N-terminal His_8_-C(GGS)_4_ tag, human Nck (S1-L1-S2-L2-S3) with an N-terminal His_8_-C(GGS)_4_ tag, human Nck (S2-L2-S3-L3-SH2) with an N-terminal His_8_-C(GGS)_4_ tag, human Nck (S3-L3-SH2) with an N-terminal His_8_-C(GGS)_4_ tag, human Nck (L3-SH2) with an N-terminal His_8_-C(GGS)_4_ tag, human Nck (S1) with an N-terminal His_8_-C(GGS)_4_ tag, human Nck (S2) with an N-terminal His_8_-C(GGS)_4_ tag, human Nck (S3) with an N-terminal His_8_-C(GGS)_4_ tag, human Nck (FL(L1 KtoE)) with an N-terminal His_8_-C(GGS)_4_ tag, human Nck (FL(L1 basic)) with an N-terminal His_8_-C(GGS)_4_ tag, human Nck (FL(L1 (GGSA)_10_)) with an N-terminal His_8_-C(GGS)_4_ tag, human SLP-76 (aa 101-420), human Sos1 (aa 1117-1319), human N-WASP (aa 451-485 of human WIP fused to aa 26-505 of human N-WASP), human N-WASP (aa 451-485 of human WIP fused to aa 26-505 of human N-WASP) with an N-terminal His_6_ tag, human N-WASP (aa 451-485 of human WIP fused to aa 26-185 and 201-505 of human N-WASP) with an N-terminal His_6_ tag and the basic region (KEKKKGKAKKKRLTK) doubled to (KEKKKGKAKKKRLTKGKEKKKGKAKKKRITK), human N-WASP (aa 451-485 of WIP fused to aa 26-185 and 201-505) with an N-terminal His_6_ tag and the basic region (KEKKKGKAKKKRLTK) replaced with a (GGS)_5_ linker, human N-WASP (aa 451-485 of human WIP fused to aa 26-185 and 201-505 of human N-WASP) with the basic region (KEKKKGKAKKKRLTK) doubled to (KEKKKGKAKKKRLTKGKEKKKGKAKKKRITK), and human N-WASP (aa 451-485 of WIP fused to aa 26-185 and 201-505) with the basic region (KEKKKGKAKKKRLTK) replaced with a (GGS)_5_ linker were expressed and purified from bacteria. Actin was purified from rabbit skeletal muscle. Myosin II was purified from chicken skeletal muscle. Rhodamine-labeled actin was purchased from Cytoskeleton, Inc. Details of constructs used in this study are listed in the Table S1.

### Protein Purification and Modification

#### Ezrin Purification

BL21(DE3) cells containing MBP-His_10_-Ezrin Actin Binding Domain (ABD) were collected by centrifugation and lysed by sonication in 50 mM NaH_2_PO_4_, 10 mM imidazole (pH 7.5), 150 mM NaCl, and 1 mM *β*ME. Centrifuge-cleared lysate was applied to a Ni-NTA column (GE Healthcare), washed with a gradient from 50 mM NaH_2_PO_4_, 10 mM imidazole (pH 7.5), 150 mM NaCl, and 1 mM *β*ME to 50 mM NaH_2_PO_4_, 50 mM imidazole (pH 7.5), 300 mM NaCl, and 1 mM *β*ME. MBP-His_10_-Ezrin(ABD) was eluted using a gradient of 10 mM➔500 mM imidazole (pH 7.5) in 50 mM NaH_2_PO_4_, 150 mM NaCl, and 1 mM *β*ME. Protein was concentrated using Amicon Ultra Centrifugal Filter units (Millipore) and MBP was cleaved by Factor Xa treatment for 10 hrs at 4°C. Cleaved protein was further purified by size exclusion chromatography using a Superdex 75 10/300 GL column in 50 mM Tris-HCl (pH 7.5), 300 mM NaCl, 1 mM DTT, and 10% glycerol.

#### Grb2 Purification

BL21(DE3) cells containing GST-Grb2 were collected by centrifugation and lysed by sonication in 25 mM Tris-HCl (pH 8.0), 200 mM NaCl, 2 mM EDTA (pH 8.0), 5 mM *β*ME, 1 mM PMSF, 1 *μ*g/ml antipain, 1 *μ*g/ml pepstatin, and 1 *μ*g/ml leupeptin. Centrifuge-cleared lysate was applied to Glutathione Sepharose 4B (GE Healthcare) and washed with 25 mM Tris-HCl (pH 8.0), 200 mM NaCl, and 1 mM DTT. GST was cleaved from protein by TEV protease treatment for 16 hrs at 4°C. Cleaved protein was applied to a Source 15 Q anion exchange column and eluted with a gradient of 0 mM ➔ 300 mM NaCl in 20 mM imidazole (pH 7.0) and 1 mM DTT followed by size exclusion chromatography using a Superdex 75 prepgrade column (GE Healthcare) in 25 mM HEPES (pH 7.5), 150 mM NaCl, 1 mM MgCl_2_, 1 mM *β*ME, and 10% glycerol.

#### Grb2_Basic_ Purification

BL21(DE3) cells containing GST-Grb2_Basic_ were collected by centrifugation and lysed by sonication in 25 mM Tris-HCl (pH 8.0), 200 mM NaCl, 2 mM EDTA (pH 8.0), 5 mM *β*ME, 1 mM PMSF, 1 *μ*g/ml antipain, 1 *μ*g/ml pepstatin, and 1 *μ*g/ml leupeptin. Centrifuge-cleared lysate was applied to Glutathione Sepharose 4B (GE Healthcare) and washed with 25 mM Tris-HCl (pH 8.0), 200 mM NaCl, and 1 mM DTT. GST was cleaved from protein by TEV protease treatment for 16 hrs at 4°C. Cleaved protein was applied to a Source 15 S cation exchange column and eluted with a gradient of 500 mM ➔ 1000 mM NaCl in 20 mM imidazole (pH 7.0) and 1 mM DTT followed by buffer exchange using a HiTrap 26/10 Desalting column (GE Healthcare) in 50 mM HEPES (pH 7.5), 250 mM NaCl, 2 mM *β*ME, and 10% glycerol.

#### LAT Purification and Modification

BL21(DE3) cells containing MBP-His_8_-LAT 48-233-His_6_ were collected by centrifugation and lysed by cell disruption (Emulsiflex-C5, Avestin) in 20 mM imidazole (pH 8.0), 150 mM NaCl, 5 mM *β*ME, 0.1% NP-40, 10% glycerol, 1 mM PMSF, 1 *μ*g/ml antipain, 1 *μ*g/ml pepstatin, and 1 *μ*g/ml leupeptin. Centrifugation-cleared lysate was applied to Ni-NTA agarose (Qiagen), washed with 10 mM imidazole (pH 8.0), 150 mM NaCl, 5 mM *β*ME, 0.01% NP-40, and 10% glycerol, and eluted with 500 mM imidazole (pH 8.0), 150 mM NaCl, 5 mM *β*ME, 0.01% NP-40, and 10% glycerol. The MBP tag and His_6_ tag were removed using TEV protease treatment for 16 hrs at 4°C. Cleaved protein was applied to a Source 15 Q anion exchange column and eluted with a gradient of 200 mM➔300 mM NaCl in 20 mM HEPES (pH 7.0) and 2 mM DTT followed by size exclusion chromatography using a Superdex 200 prepgrade column (GE Healthcare) in 25 mM HEPES (pH 7.5), 150 mM NaCl, 1 mM MgCl_2_, and 1 mM DTT. LAT was concentrated using Amicon Ultra Centrifugal Filter units (Millipore) to >400 *μ*M, mixed with 25 mM HEPES (pH 7.5), 150 mM NaCl, 15 mM ATP, 20 mM MgCl_2_, 2 mM DTT, and active GST-ZAP70 (SignalChem), and incubated for 24 hrs at 30°C. Phosphorylated LAT (pLAT) was resolved on a Mono Q anion exchange column using a shallow 250 mM ➔ 320 mM NaCl gradient in 25 mM HEPES (pH 7.5), 1 mM MgCl_2_, and 2 mM βME to separate differentially phosphorylated species of LAT. Complete LAT phosphorylation was confirmed by mass spectrometry. pLAT was then exchanged into 25 mM HEPES (pH 7.0), 150 mM NaCl, and 1 mM EDTA (pH 8.0) using a HiTrap Deslating Column (GE Healthcare). C_5_-maleimide Alexa488 was added in excess to pLAT in reducing agent-free buffer and incubated for 16 hrs at 4°C. Following the incubation 5 mM *β*ME was added to the labeling solution to quench the reaction. Excess dye was removed from Alexa488-labeled pLAT by size exclusion chromatography in 25 mM HEPES (pH 7.5), 150 NaCl, 1 mM MgCl_2_, 1 mM *β*ME, and 10% glycerol.

#### Nck and Nck Variant Purification

BL21(DE3) cells containing GST-Nck1 (or Nck variant, both His- and non-His-tagged) were collected by centrifugation and lysed by sonication in 25 mM Tris-HCl (pH 8.0), 200 mM NaCl, 2 mM EDTA (pH 8.0), 1 mM DTT, 1 mM PMSF, 1 μg/ml antipain, 1 *μ*g/ml pepstatin, and 1 *μ*g/ml leupeptin. Centrifuge-cleared lysate was applied to Glutathione Sepharose 4B (GE Healthcare) and washed with 25 mM Tris-HCl (pH 8.0), 200 mM NaCl, and 1 mM DTT. GST was cleaved from protein by TEV protease treatment for 16 hrs at 4°C. Cleaved protein was applied to a Source 15 Q anion exchange column and eluted with a gradient of 0 ➔ 200 mM NaCl in 20 mM imidazole (pH 7.0) and 1 mM DTT. Eluted protein was pooled and applied to a Source 15 S cation exchange column and eluted with a gradient of 0 ➔ 200 mM NaCl in 20 mM imidazole (pH 7.0) and 1 mM DTT. Eluted protein was concentrated using Amicon Ultra 10K concentrators and further purified by size exclusion chromatography using a Superdex 75 prepgrade column (GE Healthcare) in 25 mM HEPES (pH 7.5), 150 mM NaCl, 1 mM βME, and 10% glycerol.

#### N-WASP Purification

BL21(DE3) cells containing His_6_-N-WASP (or N-WASP variants) expressing cells were collected by centrifugation and lysed by cell disruption (Emulsiflex-C5, Avestin) in 20 mM imidazole (pH 7.0), 300 mM KCl, 5 mM βME, 0.01% NP-40, 1 mM PMSF, 1 μg/ml antipain, 1 mM benzamidine, and 1 μg/ml leupeptin. The cleared lysate was applied to Ni-NTA agarose (Qiagen), washed with 50 mM imidazole (pH 7.0), 300 mM KCl, 5 mM βME, and eluted with 300 mM imidazole (pH 7.0), 100 mM KCl, and 5 mM βME. The eluate was further purified over a Source 15 Q column using a gradient of 250 ➔ 450 mM NaCl in 20 mM imidazole (pH 7.0), and 1 mM DTT. The His_6_-tag was removed by TEV protease at 4°C for 16 hr (for His-tagged variants, no TEV treatment occurred). Cleaved N-WASP (or uncleaved for His-tagged variants) was then applied to a Source 15 S column using a gradient of 110 ➔ 410 mM NaCl in 20 mM imidazole (pH 7.0), 1 mM DTT. Fractions containing N-WASP were concentrated using an Amicon Ultra 10K concentrator (Millipore) and further purified by size exclusion chromatography using a Superdex 200 prepgrade column (GE Healthcare) in 25 mM HEPES (pH 7.5), 150 mM KCl, 1 mM βME, and 10% glycerol.

#### SLP-76 Purification and Modification

BL21(DE3) cells containing MBP-SLP-76 Acidic and Proline Rich region-His_6_ were collected by centrifugation and lysed by cell disruption (Emulsiflex-C5, Avestin) in 20 mM imidazole (pH 8.0), 150 mM NaCl, 5 mM *β*ME, 0.01% NP-40, 10% glycerol, 1 mM PMSF, 1 *μ*g/ml antipain, 1 *μ*g/ml pepstatin, and 1 *μ*g/ml leupeptin. Centrifuge-cleared lysate was applied to Ni-NTA Agarose (Qiagen), washed first with 20 mM imidazole (pH 8.0), 150 mM NaCl, 5 mM *β*ME, 0.01% NP-40, 10% glycerol, and 1 mM benzamidine, then washed with 50 mM imidazole (pH 8.0), 300 mM NaCl, 5 mM *β*ME, 0.01% NP-40, 10% glycerol, 1 mM benzamidine, and eluted with 500 mM imidazole (pH 8.0), 150 mM NaCl, 5 mM *β*ME, 0.01% NP-40, 10% glycerol, and 1 mM benzamidine. MBP cleaved by TEV protease treatment for 16 hrs at 4°C or for 2 hrs at room temperature. His_6_ was concurrently cleaved by PreScission protease treatment for 16 hrs at 4°C or for 2 hrs at room temperature. Cleaved protein was applied to a Source 15 Q anion exchange column and eluted with a gradient of 200 mM➔350 mM NaCl in 20 mM HEPES (pH 7.5) and 2 mM βME followed by size exclusion chromatography using a Superdex 200 prepgrade column (GE Healthcare) in 25 mM HEPES (pH 7.5), 150 mM NaCl, 1 mM MgCl_2_, and 1 mM DTT. To phosphorylate SLP-76, Purified SLP-76 was incubated in 50 mM HEPES (pH 7.0), 150 mM NaCl, 20 mM MgCl_2_, 15 mM ATP, 2 mM DTT, and 20 nM Active GST-ZAP70 (SignalChem). Phosphorylated SLP-76 was resolved on a Mono Q anion exchange column (GE Healthcare) using a shallow gradient from 300 mM ➔ 400 mM in 25 mM HEPES (pH 7.5) and 1 mM βME to separate differentially phosphorylated species of SLP-76. The phosphorylation state of pSLP-76 was confirmed by mass spectrometry analysis. pSLP-76 was then exchanged into a buffer containing 25 mM HEPES (pH 7.5), 150 mM NaCl, 1 mM MgCl_2_, 1 mM *β*ME, and 10% glycerol over a size exclusion column (Superdex 200 prepgrade column).

#### Sos1 Purification

BL21(DE3) cells containing GST-Sos1 were collected by centrifugation and lysed by sonication in 25 mM Tris-HCl (pH 8.0), 200 mM NaCl, 2 mM EDTA (pH 8.0), 1 mM DTT, 1 mM PMSF, 1 *μ*g/ml antipain, 1 *μ*g/ml pepstatin, and 1 *μ*g/ml leupeptin. Centrifuge-cleared lysate was applied to Glutathione Sepharose 4B (GE Healthcare) and washed with 50 mM HEPES (pH 7.0), 150 mM NaCl, 10% glycerol and 1 mM DTT. GST was cleaved from protein by PreScission protease treatment overnight at 4°C. Cleaved protein was purified by size exclusion chromatography using a Superdex 200 prepgrade column (GE Healthcare) in 50 mM HEPES (pH 7.5), 150 mM NaCl, 1 mM *β*ME, and 10% glycerol.

### Small Unilamellar Vesicle Preparation

Synthetic 1,2-dioleyl-*sn*-glycero-3-phosphocholine (POPC), 1,2-dioleoyl-*sn*-glycero-3-[(*N*-(5-amino-1-carboxypentyl)iminodiacetic acid)succinyl] (nickel salt, DGS-NTA-Ni), L-α-phosphatidylserine (Brain PS), 1,2-distearoyl-*sn*-glycero-3-phosphoethanolamine-N-[biotinyl(polyethylene glycol)-2000] (ammonium salt) (DSPE-PEG 2000 Biotin), and 1,2-dioleoyl-*sn*-glycero-3-phosphoethanolamine-N-[methoxy(polyethyleneglycol)-5000] (ammonium salt) (PEG5000 PE) were purchased from Avanti Polar Lipids. Phospholipids for reconstitution assays (93% POPC, 5% PS, 2% DGS-NTA-Ni and 0.1% PEG 5000 PE) or (98% POPC, 1% DSPE-PEG Biotin, 1% DGS-NTA-Ni, and 0.1% PEG 5000 PE) were dried under a stream of Argon, desiccated over 3 hrs, and resuspended in PBS (pH 7.3). To promote the formation of small unilamellar vesicles (SUVs), the lipid solution was repeatedly frozen in liquid N_2_ and thawed using a 37°C water bath until the solution cleared. Cleared SUV-containing solution was centrifuged at 33,500g for 45 min at 4°C. Supernatant containing SUVs was collected and stored at 4°C covered with Argon.

### Steady-State Reconstitution Assays

These methods are adapted from previously published methods (Köster et al., 2016; Su et al., 2017, 2016). Supported lipid bilayers (SLBs) containing 93% POPC, 5% PS, 2% DGS-Ni-NTA, and trace PEG 5000 PE were formed in 96-well glass-bottomed plates (Matrical). Glass was washed with Hellmanex III (Hëlma Analytics) for 4 hrs at 50°C, thoroughly rinsed with MilliQ H_2_O, washed with 6M NaOH for 30 min at 45°C two times, and thoroughly rinsed with MilliQ H_2_O followed by equilibration with 50 mM HEPES (pH 7.3), 150 mM KCl, and 1 mM TCEP. SUVs were added to cleaned wells covered by 50 mM HEPES (pH 7.3), 150 mM KCl, and 1 mM TCEP and incubated for 1 hr at 40°C to allow SUVs to collapse on glass and fuse to form the SLB. SLBs were washed with 50 mM HEPES (pH 7.3), 150 mM KCl, and 1 mM TCEP to remove excess SUVs. SLBs were blocked with 50 mM HEPES (pH 7.3), 150 mM KCl, 1 mM TCEP, and 1 mg/mL BSA for 30 min at room temperature. 20 nM His_8_-pLAT-Alexa488 and 5 nM His_10_-Ezrin(ABD) were premixed and incubated with SLBs in 50 mM HEPES (pH 7.3), 150 mM KCl, 1 mM TCEP, and 1 mg/ml BSA for 30 min. SLBs were then washed with 50 mM HEPES (pH 7.3), 75 mM KCl, 1 mM MgCl_2_, 1 mM TCEP, and 1 mg/ml BSA to remove unbound His-tagged proteins. Polymerized, 10% rhodamine-labeled actin was added to the SLB and allowed to bind to membrane-attached Ezrin for 15 min. 100 nM myosin II and 1 mM ATP was added to the membrane-bound actin network to induce steady-state movement of actin filaments by myosin II activity. Indicated amounts of soluble proteins were added to His-tagged protein- and actomyosin-bound SLBs and imaged using TIRF microscopy. Time-lapse images were captured every 15 seconds for 15 minutes. To obtain replicates for each time-lapse, 5 fields of view were captured by sequential, multi-point image acquisition. Microscopy experiments were performed in the presence of a glucose/glucose oxidase/catalase O_2_-scavenging system. Identical image acquisition settings were used for each steady-state biochemical experiment. See Table S2 for more information.

### Actomyosin Contraction Assays

These methods are adapted from previously published methods (Köster et al., 2016; Su et al., 2017, 2016). Supported lipid bilayers (SLBs) containing 93% POPC, 5% PS, 2% DGS-Ni-NTA, and trace PEG 5000 PE were formed in 96-well glass-bottomed plates (Matrical). Glass was washed with Hellmanex III (Hëlma Analytics) for 4 hrs at 50°C, thoroughly rinsed with MilliQ H_2_O, washed with 6M NaOH for 30 min at 45°C two times, and thoroughly rinsed with MilliQ H_2_O followed by equilibration with 50 mM HEPES (pH 7.3), 150 mM KCl, and 1 mM TCEP. SUVs were added to cleaned wells covered by 50 mM HEPES (pH 7.3), 150 mM KCl, and 1 mM TCEP and incubated for 1 hr at 40°C to allow SUVs to collapse on glass and fuse to form the SLB. SLBs were washed with 50 mM HEPES (pH 7.3), 150 mM KCl, and 1 mM TCEP to remove excess SUVs. SLBs were blocked with 50 mM HEPES (pH 7.3), 150 mM KCl, 1 mM TCEP, and 1 mg/mL BSA for 30 min at room temperature. 20 nM His_8_-pLAT-Alexa488 and 5 nM His_10_-Ezrin(ABD) were premixed and incubated with SLBs in 50 mM HEPES (pH 7.3), 150 mM KCl, 1 mM TCEP, and 1 mg/ml BSA for 30 min. SLBs were then washed with 50 mM HEPES (pH 7.3), 50 mM KCl, 1 mM MgCl_2_, 1 mM TCEP, and 1 mg/ml BSA to remove unbound His-tagged proteins. Polymerized, 10% rhodamine-labeled actin was added to the SLB and allowed to bind to membrane-attached Ezrin for 15 min. Indicated amounts of soluble proteins were added to His-tagged protein- and actin-bound SLBs and incubated for 15 min to allow for the formation of phase-separated condensates. TIRF microscopy was then used to capture actin contraction and associated condensate movement when 100 nM myosin II was added to the membrane-bound actin network to induce contraction of actin filaments by myosin II activity. Time-lapse images were captured every 5 seconds for up to 10 minutes, until contraction was completed. To obtain replicates for each time-lapse, 5 fields of view were captured by sequential, multi-point image acquisition. Microscopy experiments were performed in the presence of a glucose/glucose oxidase/catalase O_2_-scavenging system. Identical image acquisition settings were used for each actomyosin contraction biochemical experiment. See Table S2 for more information. These experiments were especially sensitive to light-induced artifacts, so images of nearby regions were captured outside of the field of view after the completion of actin contraction to ensure that time-lapse images were representative of condensate and actin movement rather than the result of an artifact (Figure 2 – figure supplement 1).

### Actin Binding Assays on SLBs

Supported lipid bilayers (SLBs) containing 93% POPC, 5% PS, 2% DGS-Ni-NTA, and trace PEG 5000 PE were formed in 96-well glass-bottomed plates (Matrical). Glass was washed with Hellmanex III (Hëlma Analytics) for 4 hrs at 50°C, thoroughly rinsed with MilliQ H_2_O, washed with 6M NaOH for 30 min at 45°C two times, and thoroughly rinsed with MilliQ H_2_O followed by equilibration with 50 mM HEPES (pH 7.3), 150 mM KCl, and 1 mM TCEP. SUVs were added to cleaned wells covered by 50 mM HEPES (pH 7.3), 150 mM KCl, and 1 mM TCEP and incubated for 1 hr at 40°C to allow SUVs to collapse on glass and fuse to form the SLB. SLBs were washed with 50 mM HEPES (pH 7.3), 150 mM KCl, and 1 mM TCEP to remove excess SUVs. SLBs were blocked with 50 mM HEPES (pH 7.3), 150 mM KCl, 1 mM TCEP, and 1 mg/mL BSA for 30 min at room temperature. Varied concentrations of His-tagged pLAT, Nck fragments, Nck full-length variants, or N-WASP variants were incubated with SLBs in 50 mM HEPES (pH 7.3), 150 mM KCl, 1 mM TCEP, and 1 mg/ml BSA for 30 min. SLBs were then washed with 50 mM HEPES (pH 7.3), 50 mM KCl, 1 mM MgCl_2_, 1 mM TCEP, and 1 mg/ml BSA to remove unbound His-tagged proteins. Polymerized, 10% rhodamine-labeled actin was added to the SLB and allowed to bind to membrane-bound proteins. TIRF microscopy was used to capture images. Microscopy experiments were performed in the presence of a glucose/glucose oxidase/catalase O_2_-scavenging system. Identical image acquisition settings were used for each actin-binding biochemical experiment. Images were analyzed using ImageJ (FIJI). The same brightness and contrast were applied to images within the same panels. Camera background was subtracted before calculating mean fluorescence intensities. Data from image analysis within FIJI was analyzed and graphed using GraphPad Prism v.7.

### Actin Co-sedimentation Assays

G-actin was allowed to polymerize for 1 hour at room temperature in a buffer containing 50 mM HEPES pH 7.3, 50 mM KCl, 1 mM MgCl_2_, 1 mM EGTA pH 8.0, and 1 mM ATP. 2 *μ*M F-actin and 2 *μ*M *α*-actinin, 2 *μ*M Grb2, 2 *μ*M Nck, or 2 *μ*M N-WASP were incubated at room temperature for 1 hour. Solutions containing F-actin and potential binding partners were centrifuged at 100,000 g for 20 minutes at 20°C. Supernatant and pellet components were analyzed using SDS-PAGE gels stained with coomassie blue.

### Cell Culture and Transduction

Jurkat T cells were grown in RPMI-1640 supplemented with 10% FBS, 100 U/mL penicillin, and 100 mg/mL streptomycin. Lentiviral transduction was used to make WT Jurkat T cells (E6.1) stably expressing combinations of LAT, Grb2, Gads, Nck, Grb2_Basic_, or LifeAct. LAT was inserted into either the pHR-mCitrine-tWPRE or pHR-mCherry-tWPRE backbone vector, Grb2 and Grb2_Basic_ was inserted into the pHR-mCherry-tWPRE backbone vector, Nck was inserted into the pHR-sfGFP-tWPRE backbone vector, and Gads and LifeAct were inserted into the pHR-BFP-tWPRE backbone vector.

### Activation of Jurkat T Cells

Supported lipid bilayers (SLBs) containing 98% POPC, 1% DSPE-PEG 2000 Biotin, 1% DGS-Ni-NTA, and trace PEG 5000 PE were formed in 96-well glass-bottomed plates (Matrical). Glass was washed with Hellmanex III (Hëlma Analytics) for 4 hrs at 50°C, thoroughly rinsed with MilliQ H_2_O, washed with 6M NaOH for 30 min at 45°C two times, and thoroughly rinsed with MilliQ H_2_O followed by equilibration with PBS (pH 7.3). SUVs were added to cleaned wells covered by PBS (pH 7.3) and incubated for 1 hr at 40°C to allow SUVs to collapse on glass and fuse to form the SLB. SLBs were washed with PBS (pH 7.3) to remove excess SUVs. 1 *μ*g / ml streptavidin-Alexa647 (Thermo Fisher Scientific) in PBS (pH 7.3) was added to the SLB and incubated at room temperature for 30 min. SLBs were thoroughly washed with PBS (7.3) to remove excess streptavidin-Alexa647. 5 *μ*g / ml OKT3-Biotin (Thermo Fisher Scientific) in PBS (pH 7.3) was added to the SLB and incubated at room temperature for 30 min. SLBs were thoroughly washed with PBS (pH 7.3) to remove excess OKT3-Biotin. 1 *μ*g / ml His-ICAM-1 (Thermo Fisher Scientific) in PBS (pH 7.3) was added to the SLB and incubated at room temperature for 30 min. SLBs were thoroughly washed with PBS (pH 7.3) to remove excess OKT3-biotin. PBS (pH 7.3) was exchanged to RPMI 1640 supplemented with 20 mM HEPES (pH 7.4) by washing SLBs three times. WT Jurkat T Cells expressing either LAT-mCitrine, Grb2-mCherry, and Gads-BFP, LAT-mCherry, Nck-sfGFP, and LifeAct-BFP, or LAT-mCitrine and Grb2_Basic_-mCherry were dropped onto the bilayers. Condensate mobility and composition was imaged by TIRF microscopy at 37°C. Images were captured every 5 seconds for up to 5 minutes. Identical image acquisition settings were used for each cellular experiment.

### TIRF Microscopy

TIRF images were captured using a TIRF/iLAS2 TIRF/FRAP module (Biovision) mounted on a Leica DMI6000 microscope base equipped with a Hamamatsu ImagEMX2 EM-CCD camera with a 100X 1.49 NA objective. Images were acquired using MetaMorph.software. Experiments using TIRF microscopy are described in detail in the following methods sections: Steady State Reconstitution Assays, Actomyosin Contraction Assays, Actin Binding Assays on SLBs, and Activation of Jurkat T Cells.

### Polarization Microscopy and Analysis

We used a Nikon TE-2000E microscope with custom-built polarization optical systems on an optical table. The laser beam (25 mW, OBIS 561 nm, Coherent) was routed via custom optics and focused on the back focal plane of a 100 × 1.49 NA TIRF objective (Nikon). The laser beam was circularly polarized using a combination of a half wave plate and quarter wave plate (Meadowlark Optics). The circularly polarized beam was rotated at 300 – 400 Hz in the back focal plane of the objective to achieve total internal reflection at the specimen plane to achieve isotropic excitation within the focal plane. A quadrant imaging system, as described in Mehta et al (Mehta et al., 2016) was used for instantaneous image capture along four polarization orientations at 45° increments (I0, I45, I90, I135). Time-lapse images were captured using an EMCCD camera iXon+ (Andor Technology) every 5 seconds for up to 10 minutes at 37°C. Micro – Manager (version 1.4.15) software was used to acquire images. A detailed description of the analysis can be found in Mehta et al. (Mehta et al., 2016) and Nordenfelt et al (Nordenfelt et al., 2017). Briefly, the first ten frames following IS formation from each time-lapse image set were analyzed using custom code developed in MATLAB 2014a. Calculated polarizations for each detected SiR-Actin speckle were visualized on time-lapse image panels using FIJI and plotted on a radial line from the synapse center to the synapse edge and determined to be within 45° of perpendicular or parallel form the synapse edge.

### Confocal Microscopy of Activated Jurkat T cells

Confocal images were captured using a Yokogawa spinning disk (Biovision) mounted on a Leica DMI6000 microscope base equipped with a Hamamatsu ImagEMX2 EM-CCD camera with a 100X 1.49 NA objective. Images were acquired using MetaMorph.software. SLBs were prepared for cellular activation as described above. Jurkat T cells expressing LAT-mCherry, Nck-sfGFP, and LifeAct-BFP were activated on the SLB for 5 minutes to allow the IS to form. Confocal slices were then captured with a 0.25 *μ*m step-size. 3-dimensional images were reconstructed using Matlab and the position of the dense actin ring (LifeAct-BFP) and membrane (as indicated by LAT-mCherry fluorescence) were measured and analyzed for spatial orientation using Matlab.

### Data Analysis and Display

#### 1. Drift Correction for In Vitro Movies

Due to imaging multiple time points at multiple stage positions for all *in vitro* reconstitution assay imaging, movies were subject to drift artifacts. To correct for drift, we aligned the frames in the pLAT channel by the maximum pixel-level cross correlation between adjacent frames. We then applied the shift to all channels to ensure identical corrections.

There were no drift artifacts from cellular imaging data because a single stage position was used.

#### 2. pLAT Condensates in vitro within No Actin / Actin / Actomyosin Networks at Steady State

##### 2a. Detection of pLAT Condensates

To detect pLAT condensates, we first pre-processed the pLAT condensate images by (i) subtracting inhomogeneous background, where the background image was estimated by filtering the pLAT image with a large Gaussian kernel (σ = 10 pixels), and (ii) suppressing noise by filtering the background subtracted image with a small Gaussian kernel (σ = 1). Next we detected pLAT condensates using a combination of local maxima detection (Jaqaman et al., 2008) to handle diffraction-limited condensates, which tended to be dim, and intensity-based segmentation (Otsu threshold) to handle larger condensates, which tended to be brighter. Applying these two algorithms resulted in three detection scenarios: (1) A segmented region containing one local maximum within it, taken to represent one condensate. (2) A local maximum not enclosed within a segmented region, representing a dim, diffraction-limited condensate. In this case, a circular area approximating the two-dimensional point spread function (PSF σ = 74 nm = 0.46 pixels, => circle radius = 3σ = 222 nm = 1.4 pixels) centered at the local maximum was taken in lieu of a segmented condensate area. (3) A segmented region containing multiple local maxima. This scenario could arise from overlapping nearby condensates or from fluctuating intensity within one large condensate (possibly due to incomplete mixing following the fusion of two or more individual condensates). To distinguish between these two cases, we compared the intensity variation along the line connecting each pair of local maxima in a segmented region to the intensity variation between the center and the edge of isolated condensates (scenario 1, i.e. one local maximum within one condensate). Specifically, for each pair of local maxima in a segmented region, we averaged their peak intensities and then calculated the ratio of this average to the minimum intensity along the line between them. Similarly, for isolated condensates (scenario 1), we calculated the ratio of their peak intensity to the minimum intensity at the edge (Figure 1 – figure supplement 1). This provided us with a reference distribution of peak/minimum ratios for truly separate condensates. In particular, we took the 1^st^ percentile of this distribution as a threshold to distinguish between pairs of local maxima with a segmented region belonging to separate condensates (pair peak/minimum ratio ≥ threshold) and pairs of local maxima belonging to the same condensate (pair peak/minimum ratio < threshold). If a pair of local maxima was deemed to belong to the same condensate, the local maximum with lower peak intensity was discarded. After eliminating superfluous local maxima with this procedure, we finalized the condensate segmentation and center estimation by using watershed on the originally segmented region with the remaining local maxima as seeds for the watershed algorithm (Figure 1 – figure supplement 1).

##### 2b. Tracking of pLAT Condensates

To track the detected pLAT condensates, we employed our previously developed multiple-particle tracking software “u-track” (Jaqaman et al., 2008), using a search radius of 5 pixels, a gap closing time window of 3 frames, and with the possibility of merging and splitting (all other parameters had default values). The search radius and gap closing time window were optimized by inspecting the distributions of frame-to-frame displacements and gap durations output by the software, and by visual inspection of the tracked condensates.

##### 2c. Motion Analysis of pLAT Condensates

To characterize the movement of pLAT condensates, we used previously developed moment scaling spectrum (MSS) analysis of the condensate tracks (Ewers et al., 2005; Ferrari et al., 2001; Jaqaman et al., 2011). Apart from employing new MSS slope thresholds to distinguish between immobile and confined tracks (Vega et al., 2018), all analysis procedures and parameters were as described in (Jaqaman et al., 2011).

##### 2d. Analysis of Actin Enrichment at pLAT Condensates

To quantify the enrichment of actin at pLAT condensates (Figure 1C), we employed our previously developed point-to-continuum colocalization analysis algorithm (Githaka et al., 2016). Briefly, actin enrichment in each movie was defined as the ratio of actin intensity within pLAT condensates to the actin intensity outside condensates, averaged over all condensates, at the last frame of each movie.

#### 3. pLAT Condensates In Vitro within Contractile Actomyosin Networks

##### 3a. STICS Analysis of pLAT and Actomyosin Movement

To capture the overall movement of the actomyosin network and pLAT condensates, we first applied SpatioTemporal Image Correlation Spectroscopy (STICS) analysis to each channel separately (Ashdown et al., 2014). STICS was preferred over particle tracking to analyze the pLAT channel in this assay for two reasons: 1) Experimental conditions (described above) resulted in many pLAT ➔ N-WASP condensates strongly wetting actin filaments (when compared with steady-state experiments), thus hindering accurate detection of individual condensates; and 2) using STICS on both channels allowed direct comparisons of speed and directionality at specific image sub-regions. To perform STICS analysis, we used the STICS Fiji plug-in provided by the Stowers Institute (http://research.stowers.org/imagejplugins/index.html), and the java package Miji (https://imagej.net/Miji) to call Fiji from MATLAB via a short script. Analysis was performed on image sub-regions of size 16 × 16 pixels (16 pixels = 2.56 μm) with a step size of 8 pixels across the image, and with a temporal correlation shift of 3 frames (i.e. 15 s). This was found to be the smallest shift possible to capture both condensate and actin movement without noise dominating the measurement. To capture movements relevant to contraction, STICS analysis was applied to movies only within the time interval of contraction, which lasted 10-15 frames (i.e. 50 - 75 s).

##### 3b. Analysis of pLAT and Actomyosin Co-movement

After acquiring STICS movement vector fields for the two channels (over the contraction time interval), we compared the vector magnitudes (i.e. speeds) and angles at each image sub-region between the two channels. The angle distribution (e.g. Figure 2D; taken from the *histogram* function and plotted using the *stairs* function) and speed comparison (e.g. Figure 2C; plotted using the MATLAB function *histogram2*) for any condition were comprised of all the measurements from all sub-regions of all movies representing that condition. Angle distributions were compared between conditions (or between a condition and its randomized control) via a Kolmogorov-Smirnov (KS) test. For this, due to the hyper-sensitivity of the KS test when the distributions comprise many data points (roughly >1000 data points), we performed the KS test on pairs of subsamples from the distributions (down to 500 data points each), repeated 100 times, and the average p-value of the 100 repeats was reported as the KS test p-value (e.g. in Figure 2D).

#### 4. Live Cell Data

##### 4a. Synapse and cSMAC Segmentation

To identify the location of LAT condensates relative to the synapse edge and cSMAC center, we first segmented the synapse and the cSMAC. Both were done in a semi-automated manner. For synapse segmentation, the actin channel (LifeAct – BFP) was used when available for a condition, as this channel provided the clearest synapse edge for segmentation. In conditions where actin was not labeled, the LAT channel was used. Images were first smoothed using a Gaussian kernel (σ = 2 pixels), and then an Otsu threshold was applied to separate the image into 2 levels, i.e. synapse and background (using the MATLAB function *multithresh*, which generalizes the Otsu method to determine thresholds for a multimodal distribution). We retained only the synapse at the center of the image for further analysis, identified as the largest thresholded object in the image.

For cSMAC segmentation, the LAT channel was always used. In this case, *multithresh* was tasked to determine two thresholds that would separate the LAT image into three levels. The cSMAC was taken as the largest segmented area at the highest intensity level within the segmented synapse. For early frames in which the cSMAC had not yet formed, instead of segmenting the cSMAC we used the point that would eventually become the cSMAC center as an alternative reference. Specifically, we applied the above process but on the average of all the time-lapse frames, and through this the center of the eventual cSMAC was determined and used in those early frames (area = 0 in this case since cSMAC had not formed yet).

We then visually inspected all segmentation results (synapse and cSMAC) and in some cases manually refined the segmentation using in-house software.

##### 4b. Detection and Tracking of LAT Condensates

Due to the majority of condensates being diffraction limited and a lower SNR in our cellular data, thresholding as described above in 2a for *in vitro* condensates was not appropriate for cellular analysis, as it lacked the sensitivity to detect individual condensates in cells. Instead, we detected pLAT condensates solely using local maxima detection (Jaqaman et al., 2008).

After detection, we tracked the LAT condensates in the same way and with the same parameters described in section 2b for *in vitro* condensates.

##### 4c. Defining a Normalized Radial Position Between Synapse Edge and cSMAC Center

Because synapse and cSMAC size differed between cells, and additionally synapses were not circular, we sought a unitless, normalized measure of position that allowed us to pool measurements between cells, and even from different parts of the same cell. To this end, for any point in the synapse, we drew a straight line from the cSMAC center to the synapse edge going through that point, and then defined the point’s normalized radial position as the ratio of the distance between the point and the cSMAC center to the total line length. With this, the normalized radial position ranged from zero at the cSMAC center to one at the synapse edge. All LAT condensate and actin trends were then measured vs. this normalized radial position (e.g. Figure 4C).

##### 4d. LAT Condensate Composition Analysis

To analyze the composition of LAT condensates as they moved from the synapse edge toward the cSMAC, we selected condensates based on their track duration, track geometry, track start and end position, and initial Nck/Grb2 content, as explained next in detail:

###### (1) Track duration

Only tracks lasting a minimum of 5 frames were used for any analysis, to obtain enough information per track.

###### (2) Track geometry

We reasoned that LAT condensates moving toward the cSMAC should be overall linear (asymmetric). Therefore, for this analysis we selected tracks that were approximately linear, as assessed by measuring the degree of anisotropy of the scatter of condensate positions along the track (Huet et al., 2006; Jaqaman et al., 2008).

###### (3) Track start and end position

In addition to being approximately linear, tracks used for composition analysis were filtered by their start and end positions. To this end, we used the rise and fall of actin intensity as one traversed the synapse from the edge to the cSMAC to define a position threshold (described next), such that LAT condensate tracks used for composition analysis had to start between the synapse edge and this threshold, and had to end between this threshold and the cSMAC center.

To define this position threshold, we took for every cell at each frame eight actin intensity profiles from the synapse edge to the cSMAC center, using straight lines with a 45° angle between each line and the next (Figure 4 – figure supplement 2) (using the MATLAB function *improfile*). Pooling the intensity profiles from all time points for all cells showed that the actin intensity peak had a median normalized radial position of ~0.6 (Figure 4 – figure supplement 2). Therefore, we used this as the threshold dividing the start and end position of tracks used for composition analysis.

###### (4) Initial Nck/Grb2 content

As explained in the next paragraph, for composition analysis the amount of Nck/Grb2 in a LAT condensate at any time point was normalized by its amount when the condensate first appeared (specifically the average of the first three frames to account for fluctuations). To exclude condensates that had low amounts of Nck or Grb2 to begin with, only condensates that contained an average Nck or Grb2 protein intensity (as defined next) in their first three time points greater than the average standard deviation of the background intensity in the first three time points were included in the analysis (on average 88% of the tracks surviving conditions 1-3 above).

To quantify protein (i.e. Nck, Grb2, or LAT) content in a condensate, we subtracted local background from the protein intensity inside the condensate and then took the average background subtracted intensity as a measure of protein content. To estimate local background, we determined which condensates were in each other’s proximity (referred to as “condensate aggregates”) and then calculated the average and standard deviation of intensity in a 2 pixel thick perimeter around each condensate aggregate (thus all condensates within one aggregate had the same local background level). For composition analysis over the lifetime of each track, the protein content at each of its time points was normalized by its average protein content at its first three time points. The normalized protein content was then pooled for all condensates based on their frame-by-frame normalized radial position, specifically 0.9-1 (closest to the synapse edge), 0.8-0.9, 0.7-0.8, etc. The ratio of pooled normalized protein content (either Grb2: LAT or Nck: LAT) was plotted as a minimal boxplot showing only the median and notches, indicating the 95% confidence interval of each median (e.g. Figure 4C).

##### 4e. Measuring Deviation from Straight Path for Condensate Tracks

In order to measure whether condensates retaining a molecular clutch via Grb2_Basic_ displayed aberrant movement as they traversed the two actin networks, we measured the deviation of each condensate track from a straight path towards the cSMAC. Before measuring track deviation from a straight path, it was necessary to remove the following two parts of each condensate track, as these parts were irrelevant to the question of interest: (1) the initial time points of a track where the condensate was moving along the synapse edge instead of toward the cSMAC; and (2) the late time points of a track where the condensate had entered the cSMAC (Figure 4 – figure supplement 2).

To identify the initial part of a track, i.e. condensate movement along the synapse edge rather than toward the cSMAC, we measured both the frame-to-frame change in distance between the condensate and the cSMAC center and the frame-to-frame change in angle between consecutive displacements taken by the condensate. The rationale was that before a condensate began to move towards the cSMAC, the change in distance should be low, while the change in angle should be high. On the other hand, once a condensate began to move towards the center, the change in distance should be high, while the change in angle should be low. Therefore, to detect the transition between moving along the synapse edge and moving toward the cSMAC, we took the ratio of the change in distance to change in angle at every track time point and chose the first time point with a value in the top 10% of all ratios within the track as the transition point. In the majority of cases (>90% of tracks), the condensate position at this time point was before the actin position threshold previously discussed. However, in some cases, primarily in the case of shorter tracks, the condensate position at the transition time point chosen in this manner was after the actin position threshold. In this case, the transition time point was taken as the first time point with a change in distance-to-change in angle ratio in the top 25%. Manual inspection verified that with this strategy all transition time points were indeed before the actin position threshold.

After the initial and final track parts were removed, the remaining track segment was transformed such that the beginning and end were set to zero on the y-axis. Thus any non-zero y-position along the track could be directly measured as a deviation from a straight path. Additionally, tracks were flipped if necessary so that the majority of their deviations were in the positive direction. The entire deviation distribution from all time points of all tracks was then taken from the *histogram* function and plotted using the *stairs* function (Figure 5F). Example tracks were plotted using the function *scatter* and colored to represent the deviation value at each position (Figure 5E).

##### 4f. Measuring Extent of Photobleaching in Live Cell Data

In order to determine whether photobleaching might contribute to measured changes in condensate composition, we measured the change in cell background intensity over time. The mean intensity within the segmented synapse, but outside the segmented cSMAC and detected condensate areas, was taken for each cell for every frame. All mean intensity measurements were normalized by the mean intensity of the first frame for each cell. The ratio of pooled normalized mean intensity (either Grb2: LAT or Nck: LAT) was plotted as a minimal boxplot showing only the median and notches, indicating the 95% confidence interval of each median (Figure 4 – figure supplement 3).

**Table 1.**
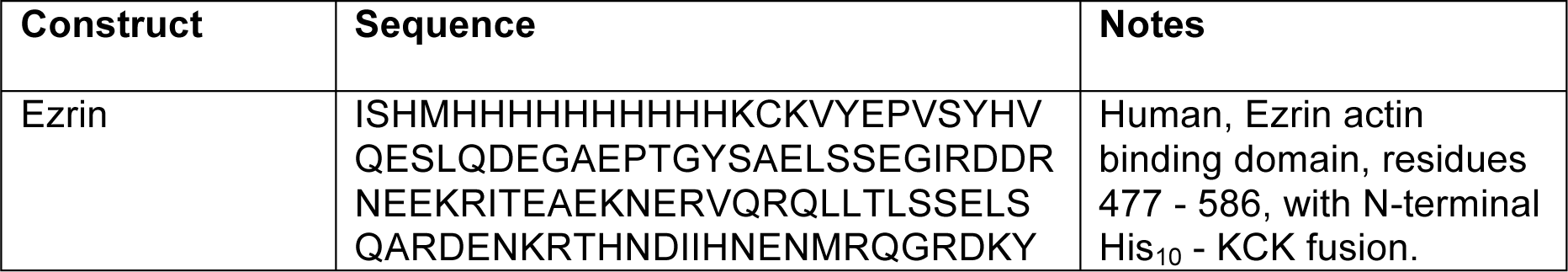
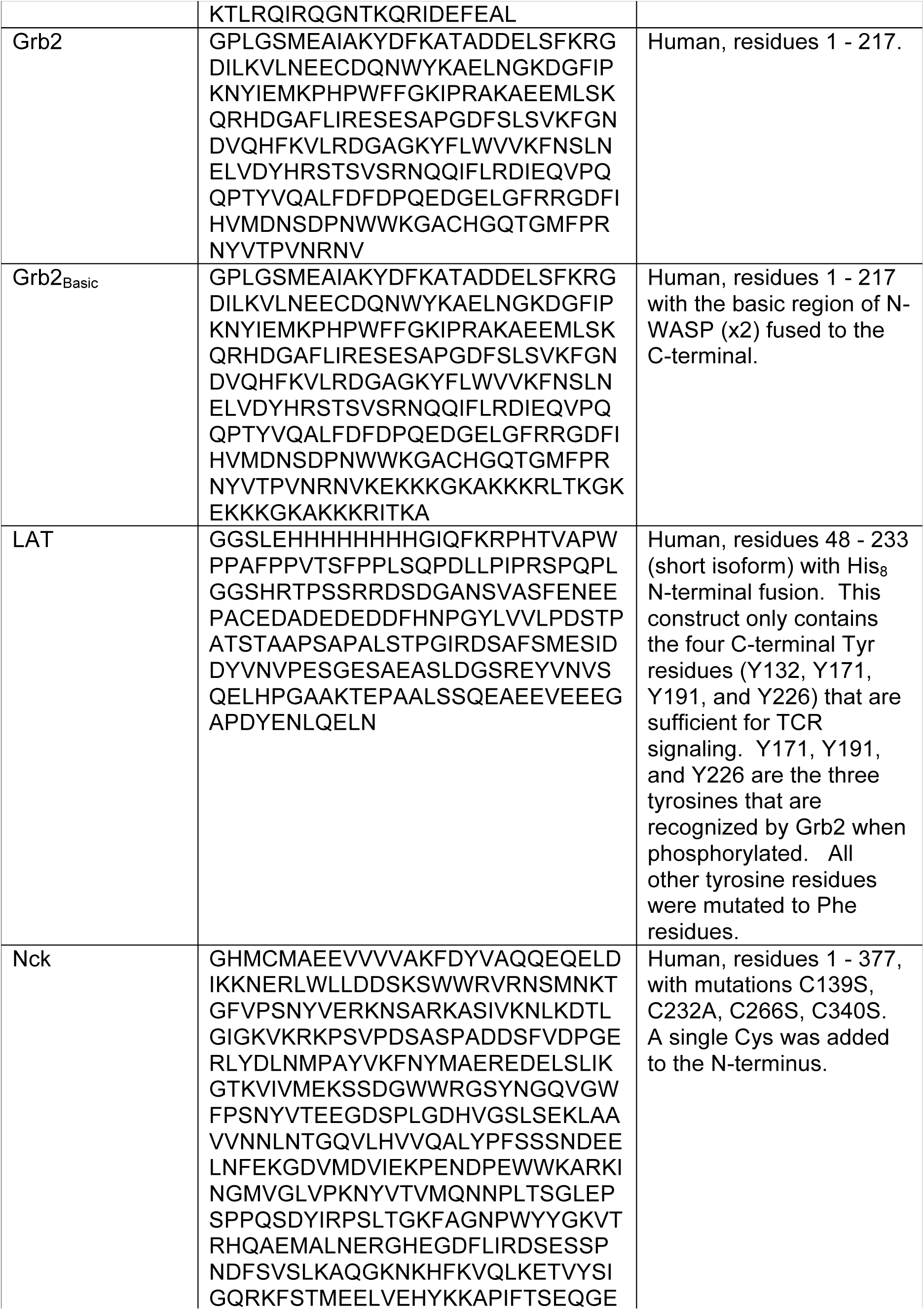
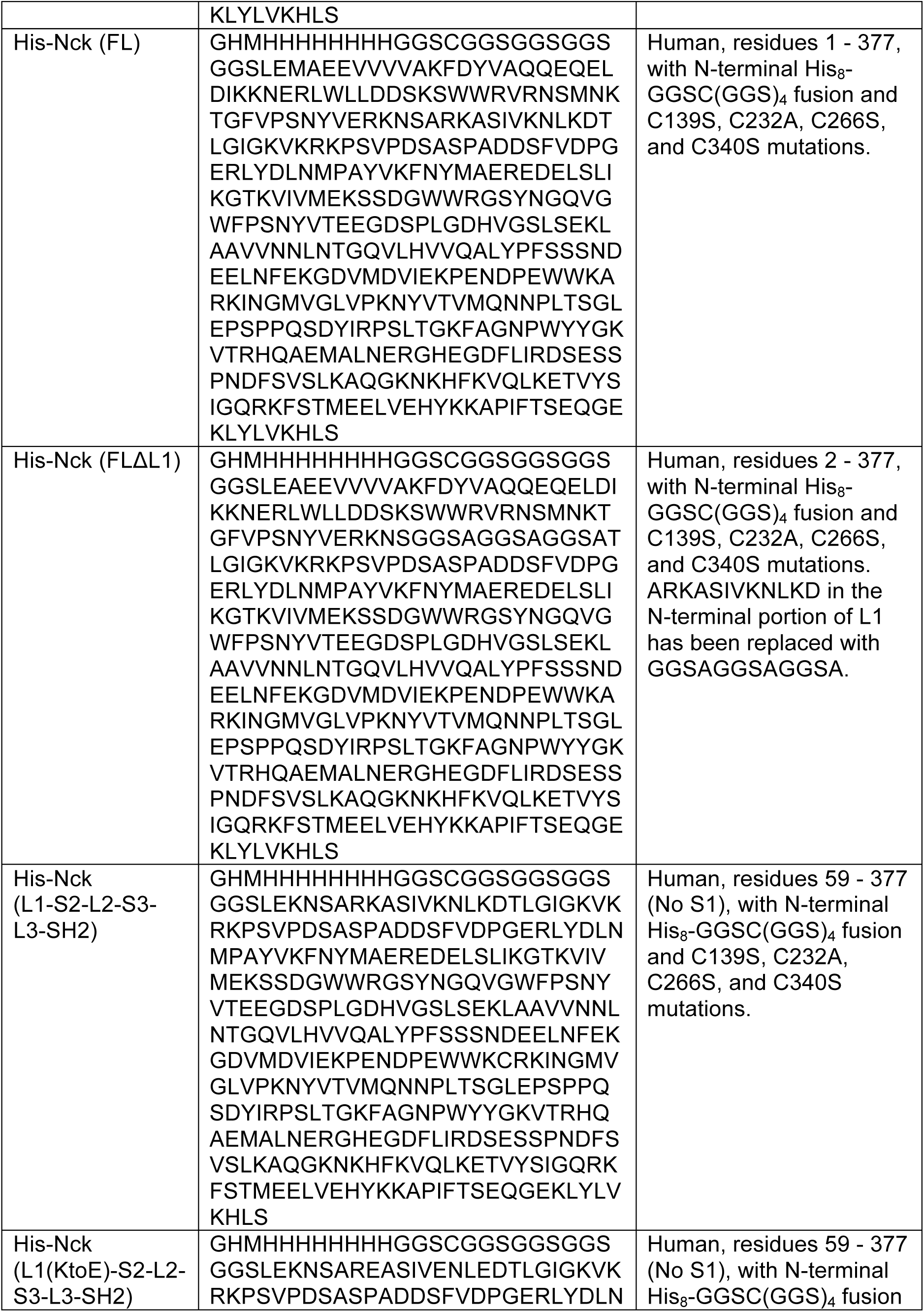
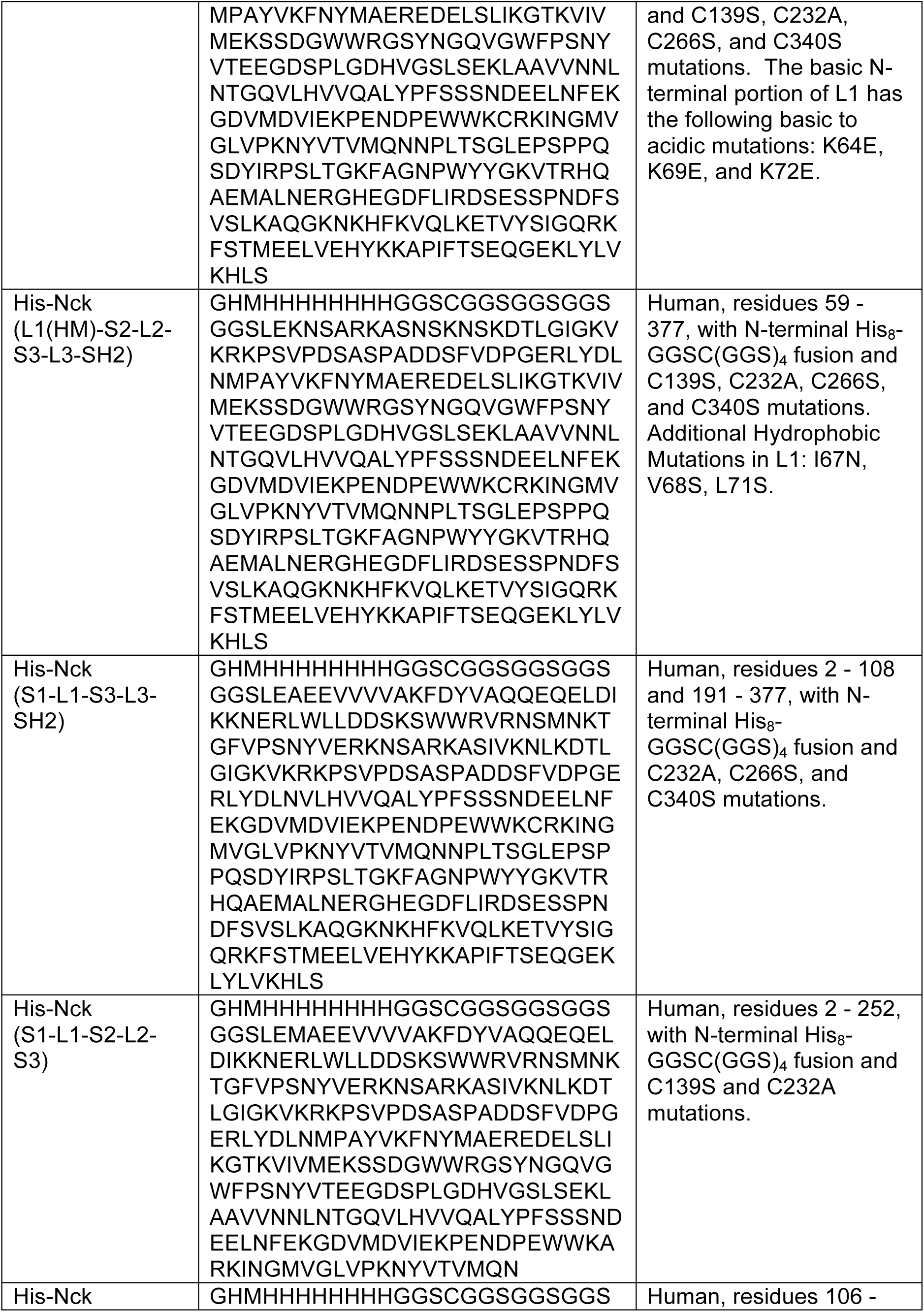
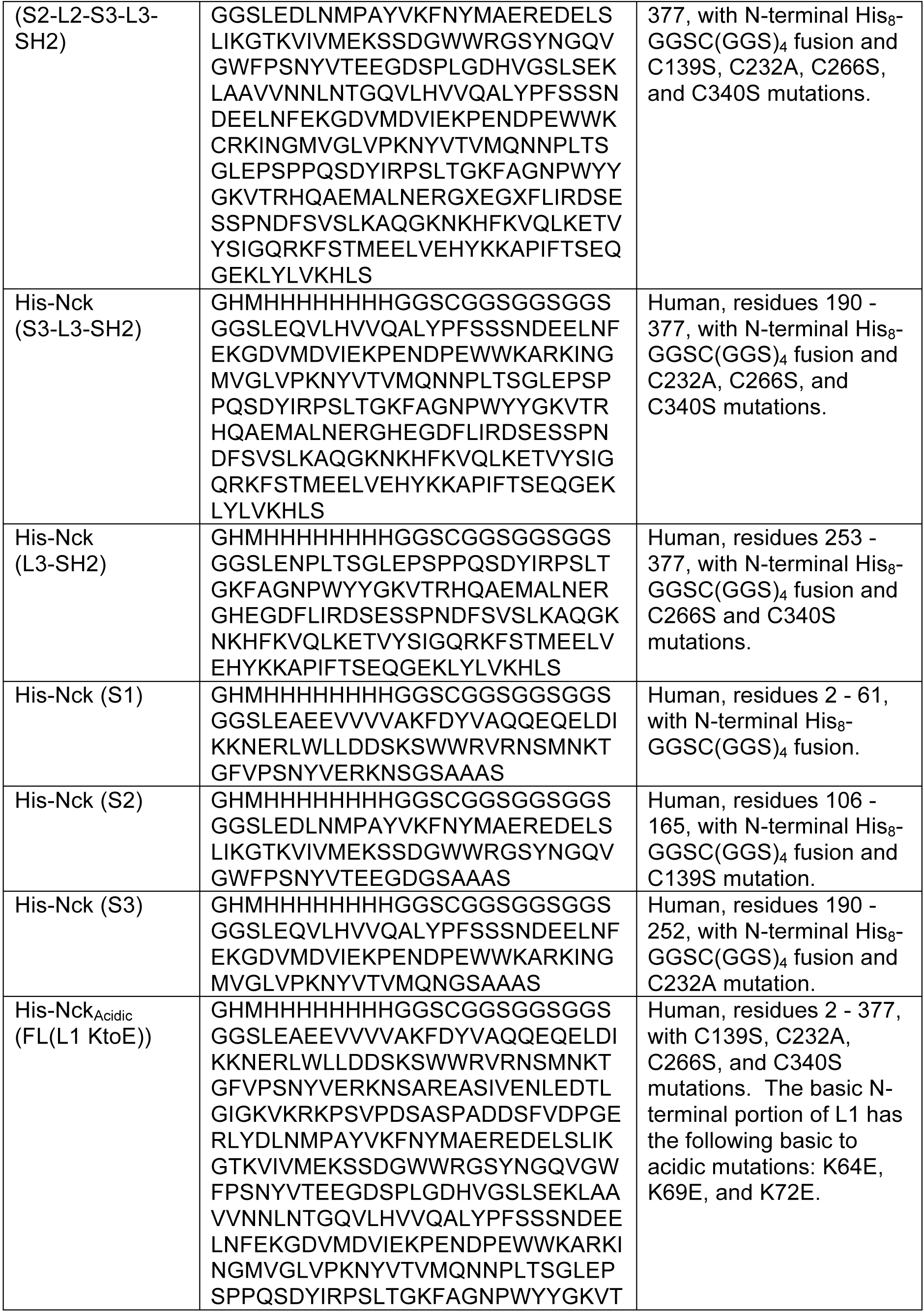
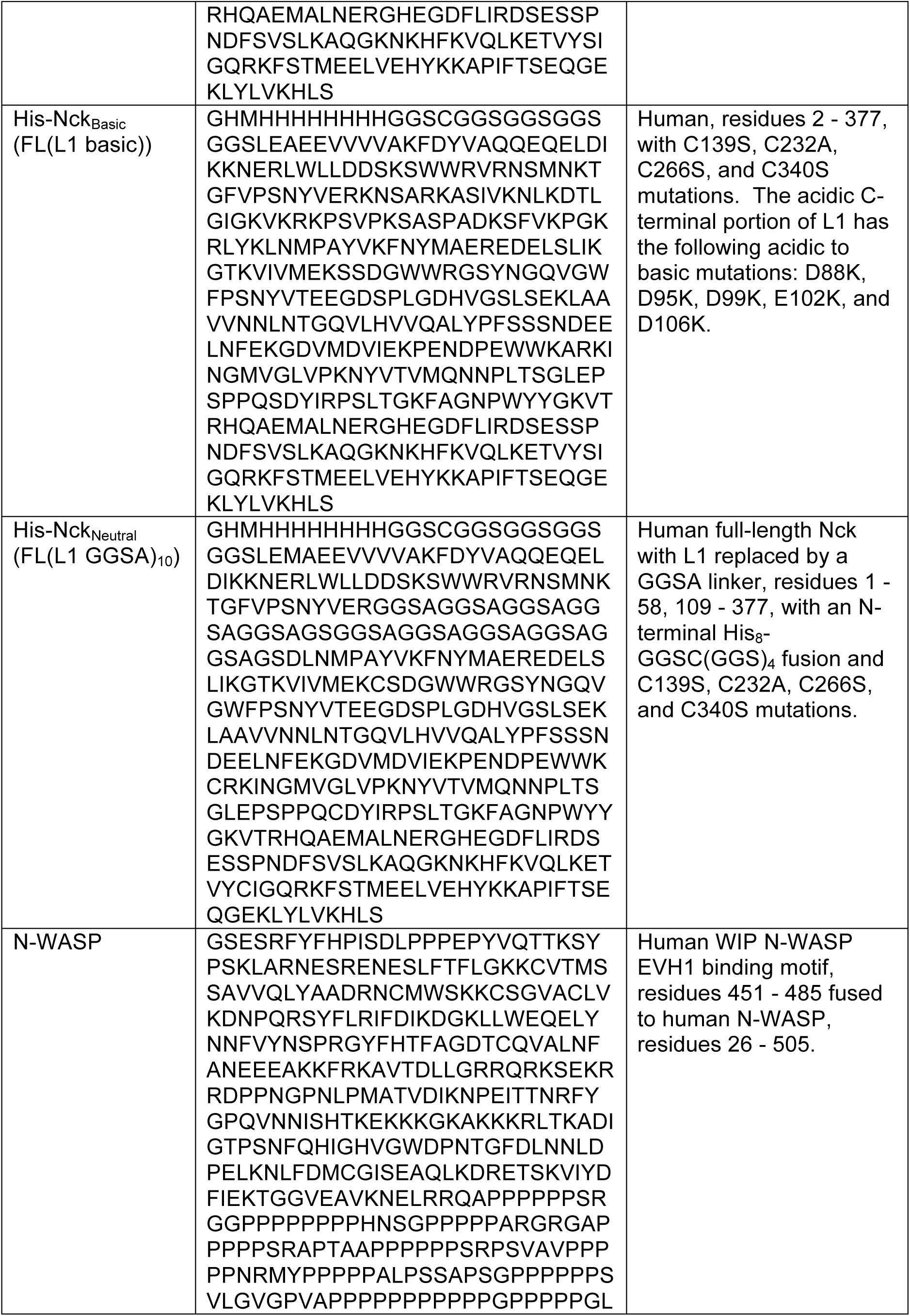
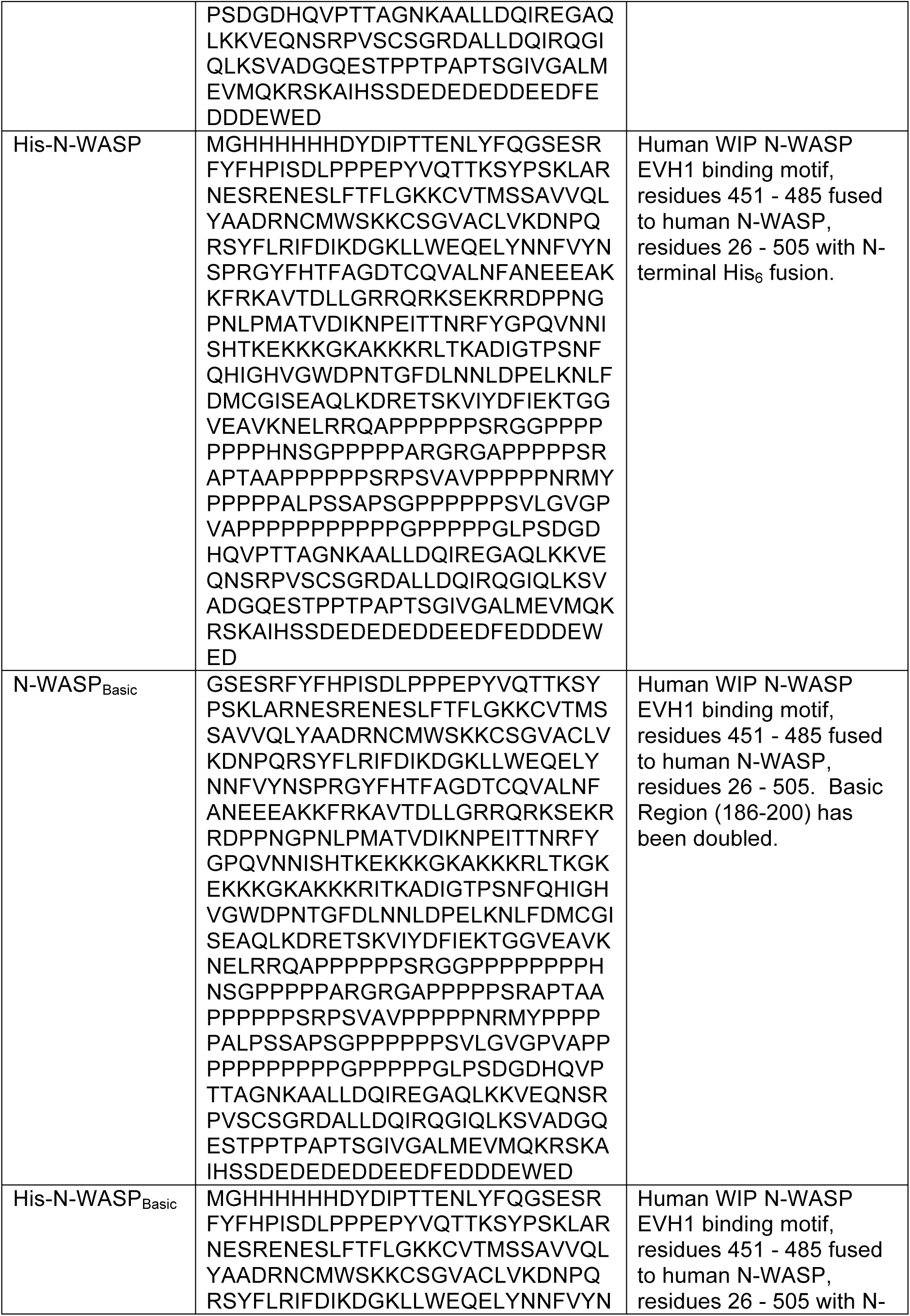
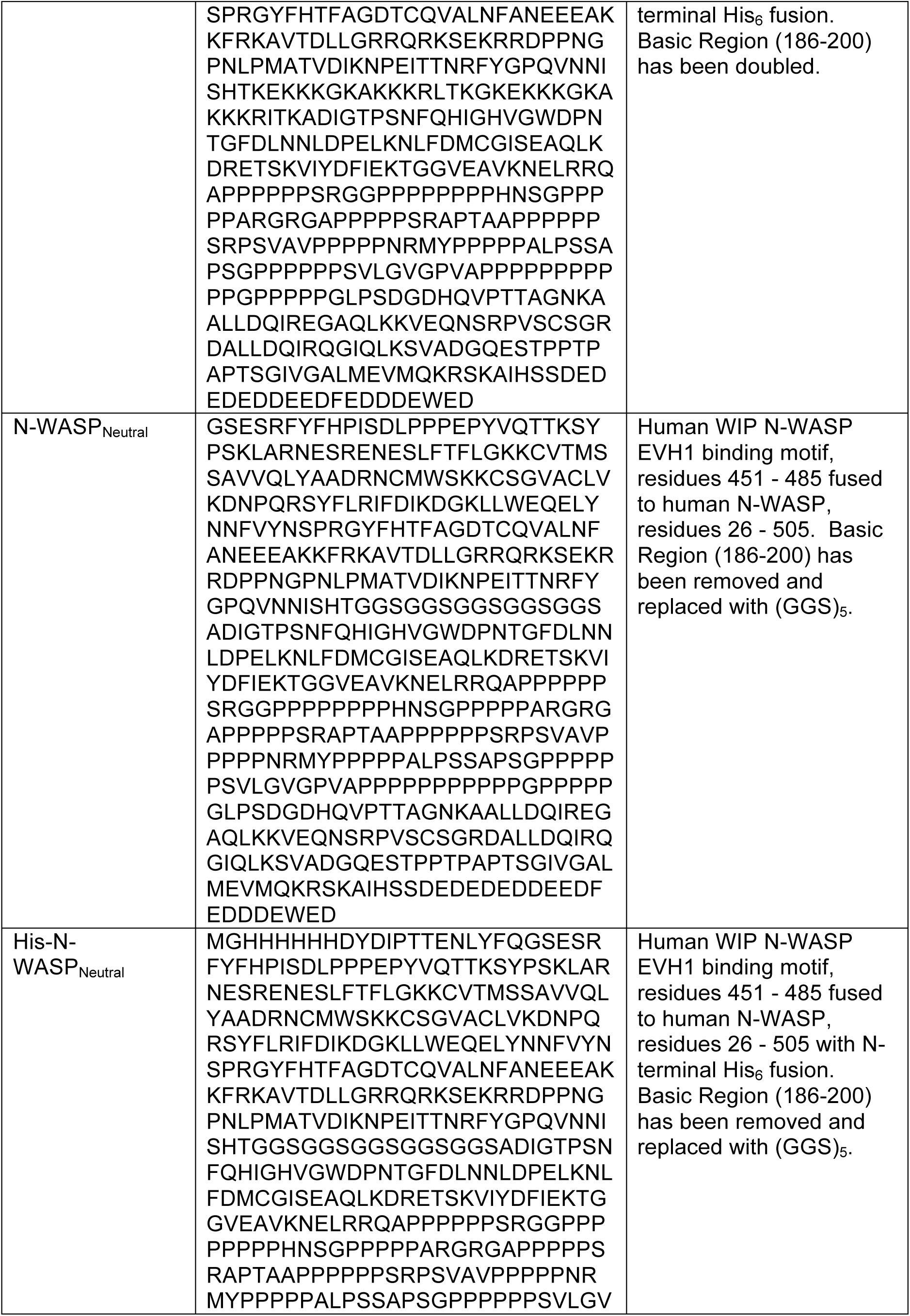
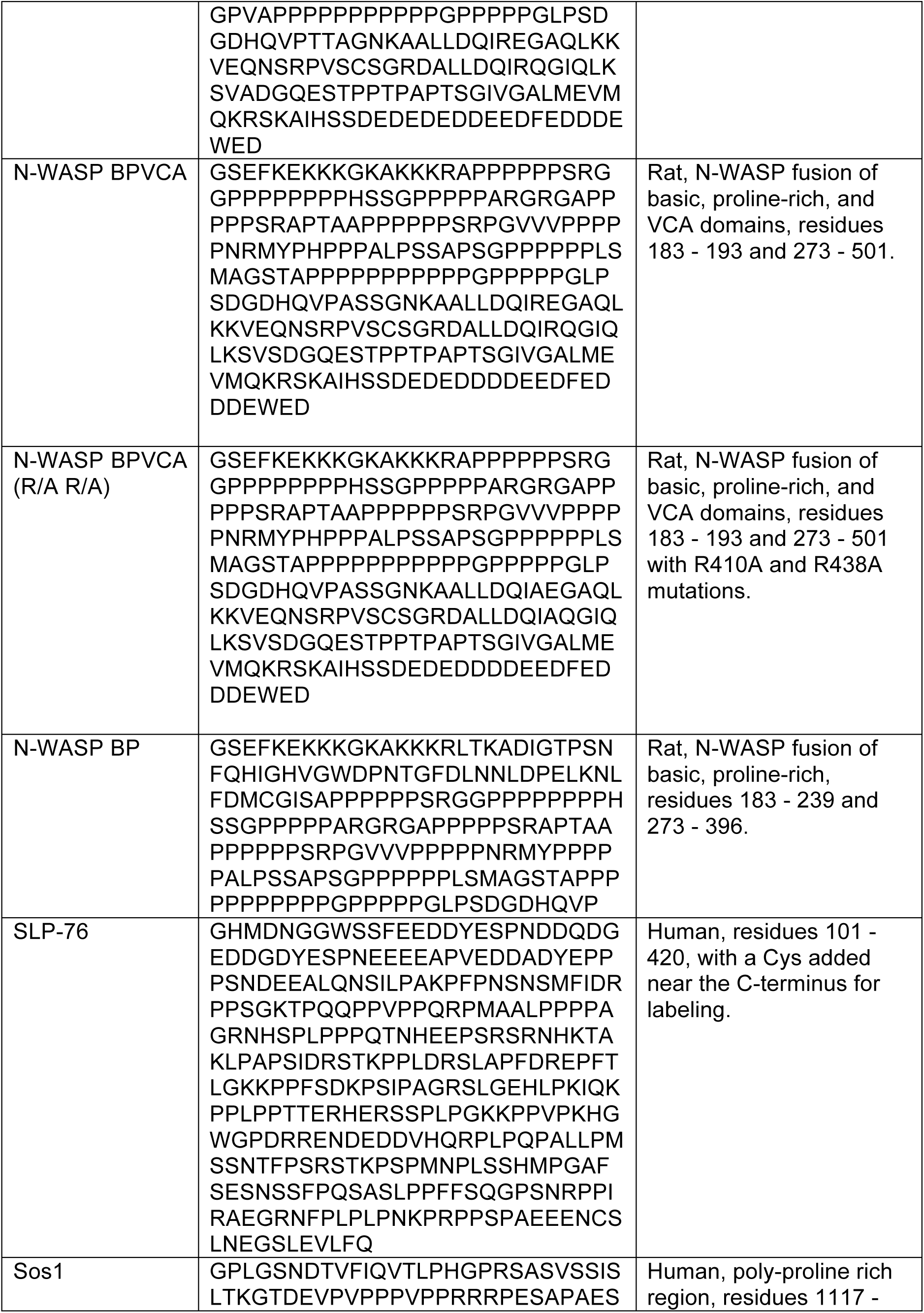
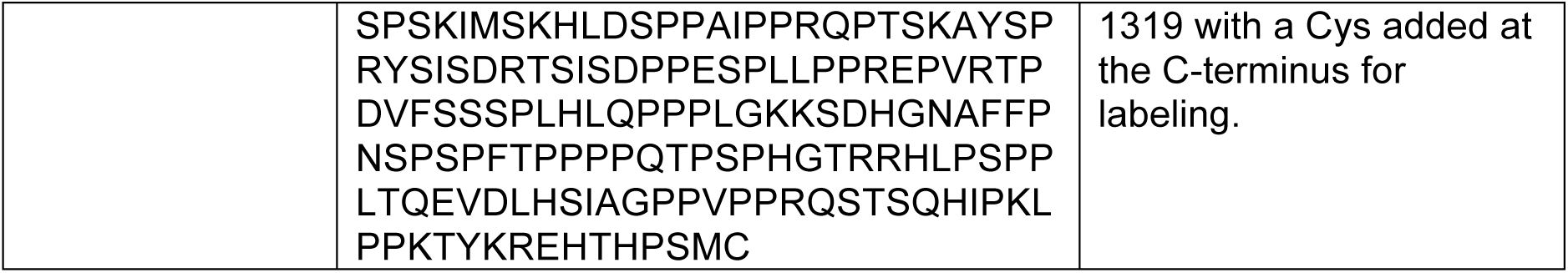
Sequences of constructs used in the study.

**Table 2.**
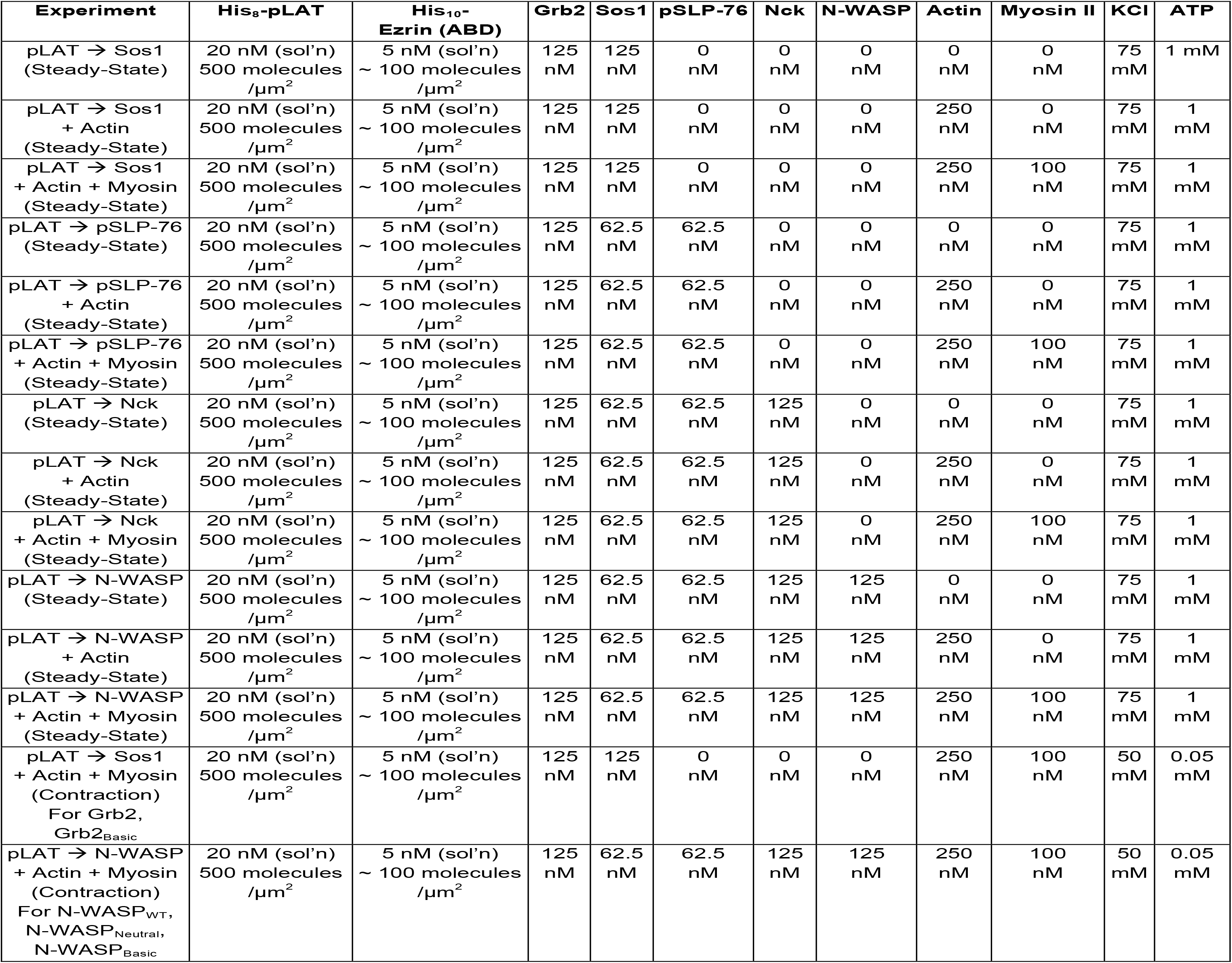
Experimental conditions used in steady-state and actomyosin contraction assays. 50 mM HEPES pH 7.3, 1 mM MgCl_2_, 1 mM TCEP, 1 mg / ml BSA, and 0.2 mg / ml glucose oxidase + 0.035 mg / ml catalase + 1 mM glucose (oxygen scavenging system) were included in every assay.

| Experiment | His<sub>8</sub>-pLAT | His<sub>10</sub>-<br>Ezrin (ABD) | Grb2 | Sos1 | pSLP-76 | Nck | N-WASP | Actin | Myosin II | KCl | ATP |
| --- | --- | --- | --- | --- | --- | --- | --- | --- | --- | --- | --- |
| pLAT → Sos1<br>(Steady-State) | 20 nM (sol'n)<br>500 molecules<br>/μm<sup>2</sup> | 5 nM (sol'n)<br>~ 100 molecules<br>/μm<sup>2</sup> | 125<br>nM | 125<br>nM | 0<br>nM | 0<br>nM | 0<br>nM | 0<br>nM | 0<br>nM | 75<br>mM | 1 mM |
| pLAT → Sos1<br>+ Actin<br>(Steady-State) | 20 nM (sol'n)<br>500 molecules<br>/μm<sup>2</sup> | 5 nM (sol'n)<br>~ 100 molecules<br>/μm<sup>2</sup> | 125<br>nM | 125<br>nM | 0<br>nM | 0<br>nM | 0<br>nM | 250<br>nM | 0<br>nM | 75<br>mM | 1<br>mM |
| pLAT → Sos1<br>+ Actin + Myosin<br>(Steady-State) | 20 nM (sol'n)<br>500 molecules<br>/μm<sup>2</sup> | 5 nM (sol'n)<br>~ 100 molecules<br>/μm<sup>2</sup> | 125<br>nM | 125<br>nM | 0<br>nM | 0<br>nM | 0<br>nM | 250<br>nM | 100<br>nM | 75<br>mM | 1<br>mM |
| pLAT → pSLP-76<br>(Steady-State) | 20 nM (sol'n)<br>500 molecules<br>/μm<sup>2</sup> | 5 nM (sol'n)<br>~ 100 molecules<br>/μm<sup>2</sup> | 125<br>nM | 62.5<br>nM | 62.5<br>nM | 0<br>nM | 0<br>nM | 0<br>nM | 0<br>nM | 75<br>mM | 1<br>mM |
| pLAT → pSLP-76<br>+ Actin<br>(Steady-State) | 20 nM (sol'n)<br>500 molecules<br>/μm<sup>2</sup> | 5 nM (sol'n)<br>~ 100 molecules<br>/μm<sup>2</sup> | 125<br>nM | 62.5<br>nM | 62.5<br>nM | 0<br>nM | 0<br>nM | 250<br>nM | 0<br>nM | 75<br>mM | 1<br>mM |
| pLAT → pSLP-76<br>+ Actin + Myosin<br>(Steady-State) | 20 nM (sol'n)<br>500 molecules<br>/μm<sup>2</sup> | 5 nM (sol'n)<br>~ 100 molecules<br>/μm<sup>2</sup> | 125<br>nM | 62.5<br>nM | 62.5<br>nM | 0<br>nM | 0<br>nM | 250<br>nM | 100<br>nM | 75<br>mM | 1<br>mM |
| pLAT → Nck<br>(Steady-State) | 20 nM (sol'n)<br>500 molecules<br>/μm<sup>2</sup> | 5 nM (sol'n)<br>~ 100 molecules<br>/μm<sup>2</sup> | 125<br>nM | 62.5<br>nM | 62.5<br>nM | 125<br>nM | 0<br>nM | 0<br>nM | 0<br>nM | 75<br>mM | 1<br>mM |
| pLAT → Nck<br>+ Actin<br>(Steady-State) | 20 nM (sol'n)<br>500 molecules<br>/μm<sup>2</sup> | 5 nM (sol'n)<br>~ 100 molecules<br>/μm<sup>2</sup> | 125<br>nM | 62.5<br>nM | 62.5<br>nM | 125<br>nM | 0<br>nM | 250<br>nM | 0<br>nM | 75<br>mM | 1<br>mM |
| pLAT → Nck<br>+ Actin + Myosin<br>(Steady-State) | 20 nM (sol'n)<br>500 molecules<br>/μm<sup>2</sup> | 5 nM (sol'n)<br>~ 100 molecules<br>/μm<sup>2</sup> | 125<br>nM | 62.5<br>nM | 62.5<br>nM | 125<br>nM | 0<br>nM | 250<br>nM | 100<br>nM | 75<br>mM | 1<br>mM |
| pLAT → N-WASP<br>(Steady-State) | 20 nM (sol'n)<br>500 molecules<br>/μm<sup>2</sup> | 5 nM (sol'n)<br>~ 100 molecules<br>/μm<sup>2</sup> | 125<br>nM | 62.5<br>nM | 62.5<br>nM | 125<br>nM | 125<br>nM | 0<br>nM | 0<br>nM | 75<br>mM | 1<br>mM |
| pLAT → N-WASP<br>+ Actin<br>(Steady-State) | 20 nM (sol'n)<br>500 molecules<br>/μm<sup>2</sup> | 5 nM (sol'n)<br>~ 100 molecules<br>/μm<sup>2</sup> | 125<br>nM | 62.5<br>nM | 62.5<br>nM | 125<br>nM | 125<br>nM | 250<br>nM | 0<br>nM | 75<br>mM | 1<br>mM |
| pLAT → N-WASP<br>+ Actin + Myosin<br>(Steady-State) | 20 nM (sol'n)<br>500 molecules<br>/μm<sup>2</sup> | 5 nM (sol'n)<br>~ 100 molecules<br>/μm<sup>2</sup> | 125<br>nM | 62.5<br>nM | 62.5<br>nM | 125<br>nM | 125<br>nM | 250<br>nM | 100<br>nM | 75<br>mM | 1<br>mM |
| pLAT → Sos1<br>+ Actin + Myosin<br>(Contraction)<br>For Grb2,<br>Grb2<sub>Basic</sub> | 20 nM (sol'n)<br>500 molecules<br>/μm<sup>2</sup> | 5 nM (sol'n)<br>~ 100 molecules<br>/μm<sup>2</sup> | 125<br>nM | 125<br>nM | 0<br>nM | 0<br>nM | 0<br>nM | 250<br>nM | 100<br>nM | 50<br>mM | 0.05<br>mM |
| pLAT → N-WASP<br>+ Actin + Myosin<br>(Contraction)<br>For N-WASP<sub>WT</sub>,<br>N-WASP<sub>Neutral</sub>,<br>N-WASP<sub>Basic</sub> | 20 nM (sol'n)<br>500 molecules<br>/μm<sup>2</sup> | 5 nM (sol'n)<br>~ 100 molecules<br>/μm<sup>2</sup> | 125<br>nM | 62.5<br>nM | 62.5<br>nM | 125<br>nM | 125<br>nM | 250<br>nM | 100<br>nM | 50<br>mM | 0.05<br>mM |

## Acknowledgements

We thank L. Rice, J. Hammer III, and our fellow HCIA Summer Institute scientists for stimulating discussions about this study. This work was supported by a Howard Hughes Medical Institute Collaborative Innovation Award, the Welch Foundation (I-1544 to M.K.R.), a J.C. Bose Fellowship from the Department of Science and Technology, government of India (S.M.), a Margadarshi Fellowship from the Wellcome Trust – Department of Biotechnology, India Alliance (IA/M/15/1/502018 to S.M.), NIH (R01 GM100160 to T.T.) (R35 GM119619 to K.J.), a CPRIT Recruitment Award (R1216 to K.J.), and the UT Southwestern Endowed Scholars Program (K.J.). Research in the Rosen lab is supported by the Howard Hughes Medical Institute. J.A.D. was supported by a National Research Service Award F32 (F32 DK101188). A.R.V. was supported by a CPRIT Training Grant (RP140110, PI: Michael White). D.V.K. was supported by fellowships of the AXA Research Fund and the National Centre for Biological Sciences, Tata Institute for Fundamental Research. X.S. was supported by a Cancer Research Institute Irvington postdoctoral fellowship.

---

**Figure 1 – figure supplement 1.**
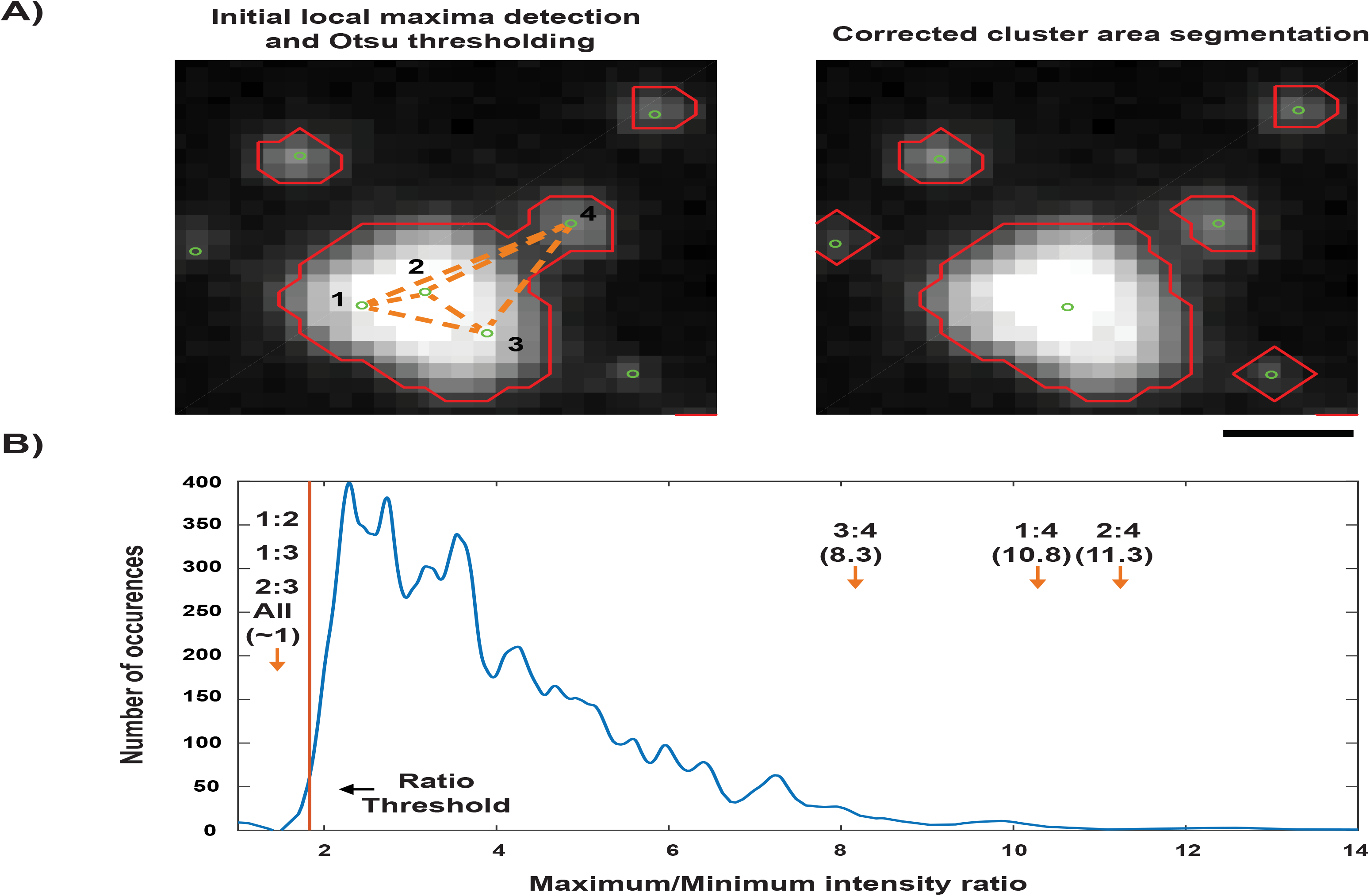
Detection process for LAT condensates *in vitro*. **(A)** (left) An example of the initial thresholding and local maxima detection for LAT condensates. Orange dotted lines indicate pair-wise comparisons that are made to remove superfluous local maxima. (right) Thresholded areas and local maxima after correction steps. **(B)** The distribution of maximum/minimum intensity ratios taken from segmented areas with a single local maximum. The threshold (1^st^ percentile of distribution) used to discard local maxima is shown as an orange line. The maximum/minimum intensity ratios of the pairs from **(A)** are shown with orange arrows.

**Figure 2 – figure supplement 1.**
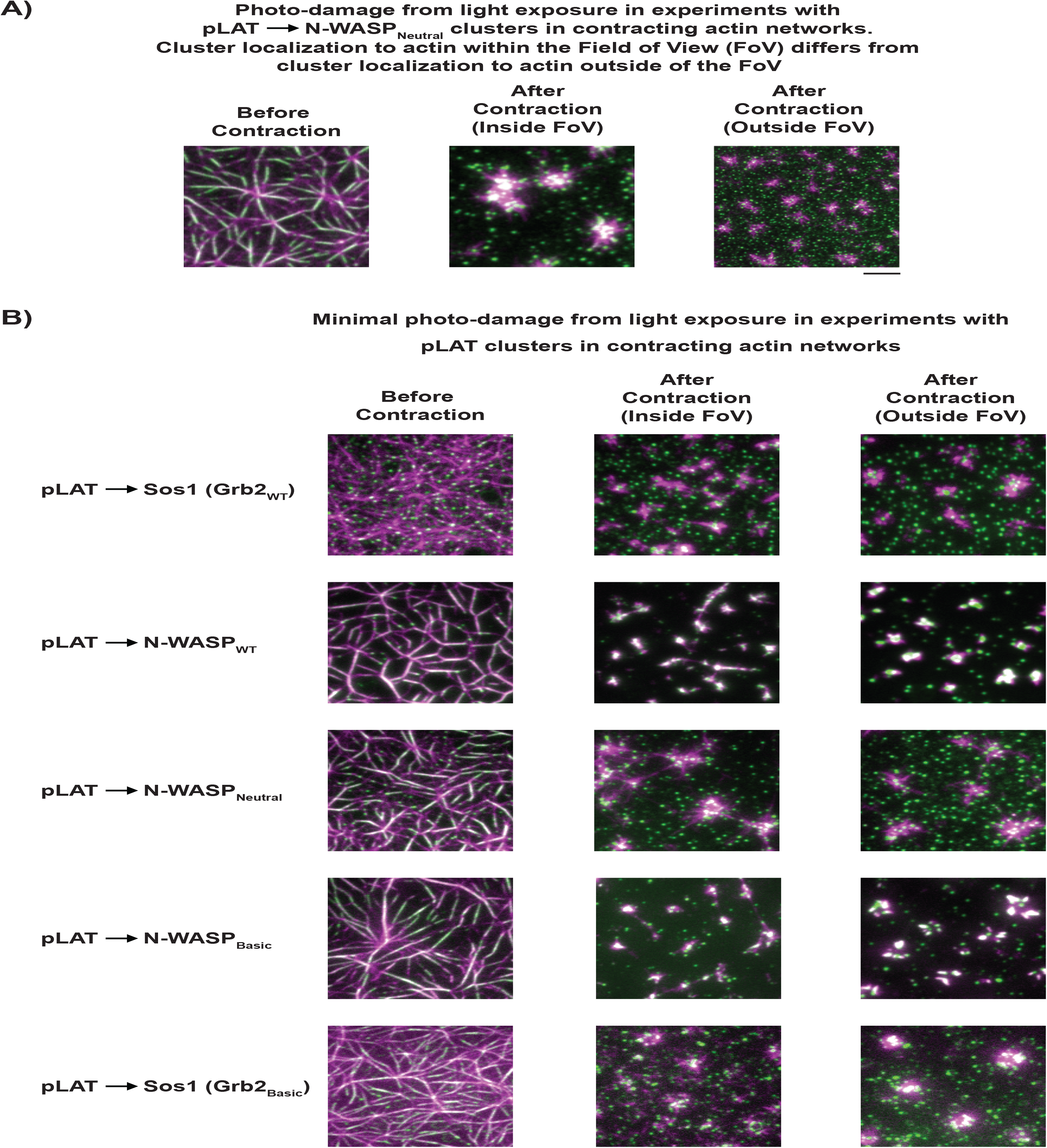
In actomyosin contractile assays, imaging conditions were chosen such that photo-damage was minimized to avoid artifactual results. **(A)** TIRF microscopy images of LAT condensate movement following myosin II-induced actin contraction inside or outside the field of view (FOV) when care was not taken to minimize photo-damage. (left) Representative image of pLAT ➔ N-WASP_Neutral_ condensate (green) wetting of rhodamine-actin filaments (magenta) prior to induced actin contraction. (center) Image of same FOV after imaging during myosin II-induced actin contraction. (right) Image of region outside of FOV (i.e. not imaged during contraction) obtained immediately following actin contraction. Comparing center and right images indicates that photo-damage can increase interactions between condensates and actin, leading to artifactual co-localization following contraction. **(B)** When imaging is performed with low laser power (as done in our experiments), regions inside and outside of the FOV during actin contraction show similar co-localization of condensates with actin asters. Figure shows representative images for a variety of pLAT condensates types captured in the FOV prior to myosin II-induced actin contraction (left), in the FOV after myosin II-induced contraction (center), and outside of the FOV after contraction (right).

**Figure 3 – figure supplement 1.**
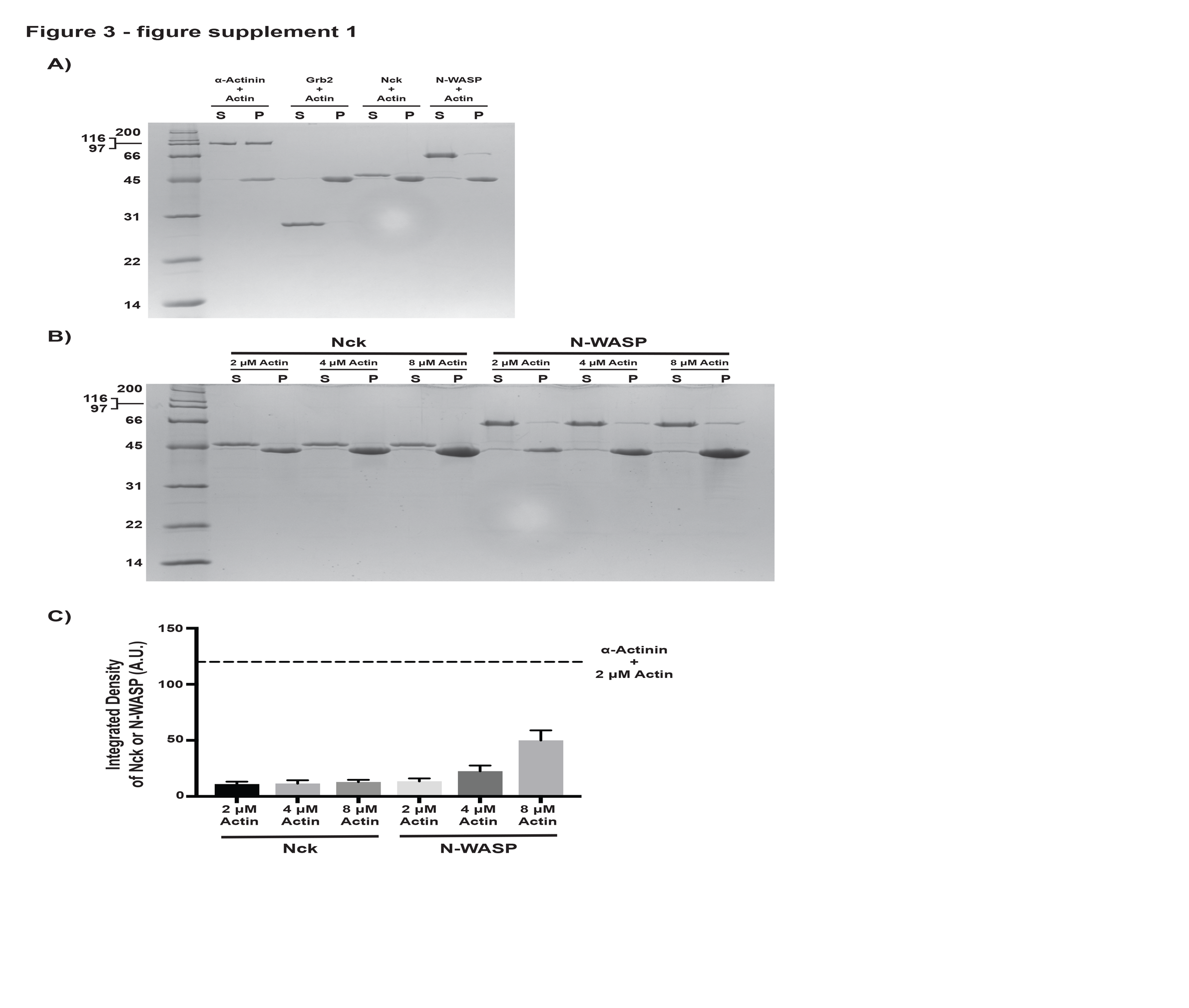
Nck does not co-sediment with actin filaments. N-WASP co-sediments with actin filaments, but not to the same degree as α-actinin, a known actin filament binding protein. (**A)** Coomassie blue-stained SDS-PAGE gel showing co-sedimentation of α-actinin (positive control), Grb2 (negative control), Nck, and N-WASP with actin filaments. **(B)** Coomassie blue-stained SDS-PAGE gel showing co-sedimentation of Nck or N-WASP with increasing actin filament concentration. **(C)** Quantification of Nck and N-WASP bands shown in **(B)**. Error bars report +/- s.d. from N = 3 independent co-sedimentation assays and gels.

**Figure 3 – figure supplement 2.**
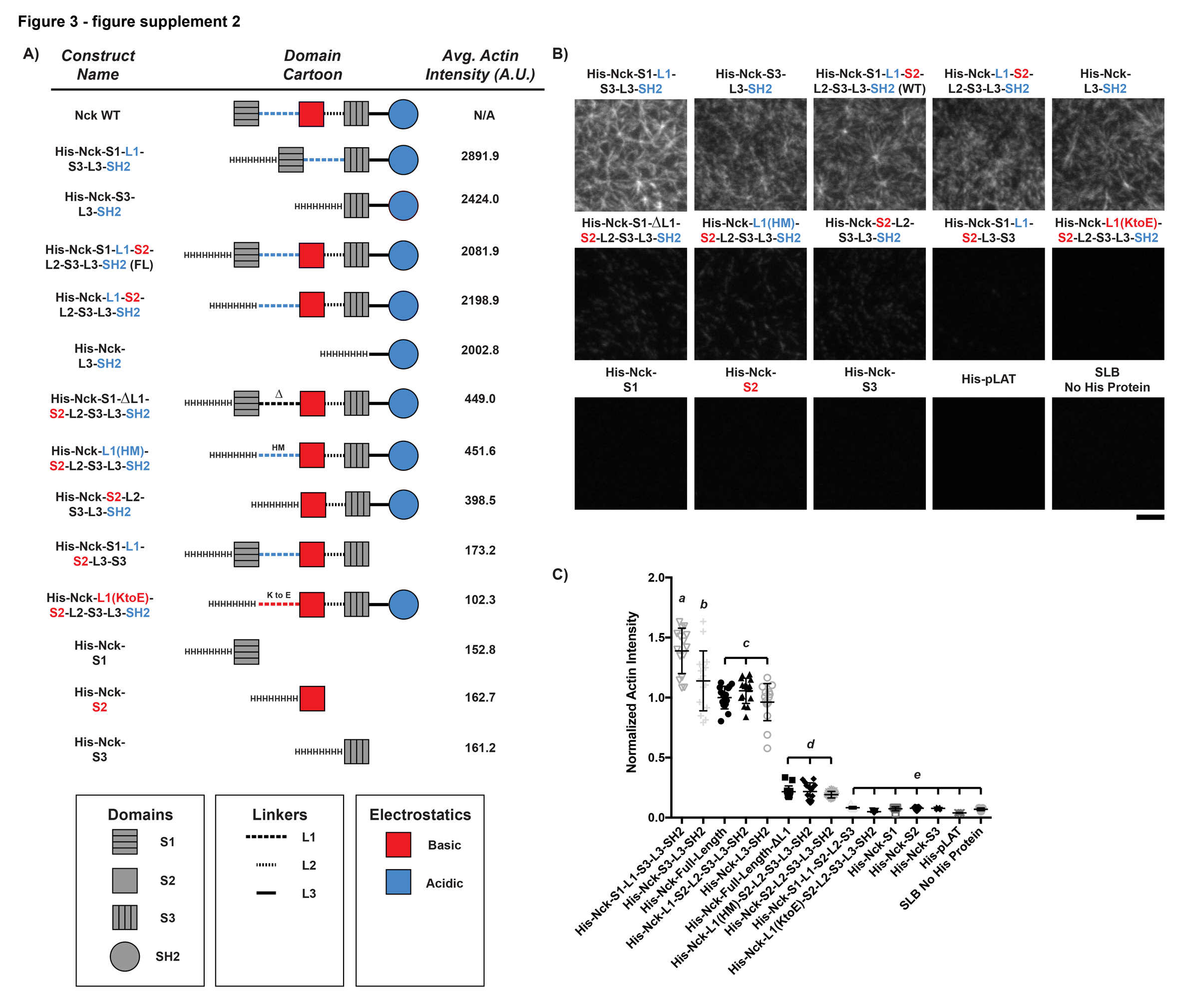
The ability of His-tagged Nck variants to recruit actin filaments to an SLB depends on the number of basic regions (concentrated in L1 and the SH2 domain) vs. the number of acidic regions (concentrated in the second SH3 domain). If the number of basic regions > the number of acidic regions, the Nck variant will recruit actin filaments. **(A)** Names (left), domain schematics (center), and average actin intensity recruited to the bilayer (right) for each Nck variant. **(B)** TIRF microscopy images of actin recruited to SLBs coated with His-tagged Nck variants or His-tagged pLAT as a control. Scale Bar = 5 μm. All images set to same intensity range. **(C)** Normalized fluorescence intensity of rhodamine-actin (normalized to the average actin filament intensity of His-Nck-L3-SH2 experiments) recruited to SLBs coated with His-tagged Nck variants. Shown are individual data points and their mean +/- s.d. from N = 15 fields of view from 3 independent experiments (5 FOV per experiment). Significance was determined using analysis of variance with Tukey’s multiple comparison test and found that p < 0.0332 for b vs. c, p < 0.0021 for d vs. e, and p < 0.0001 for a vs. b, a vs. c, a vs. d, a vs. e, b vs. d, b vs. e, c vs. d, and c vs. e.

**Figure 3 – figure supplement 3.**
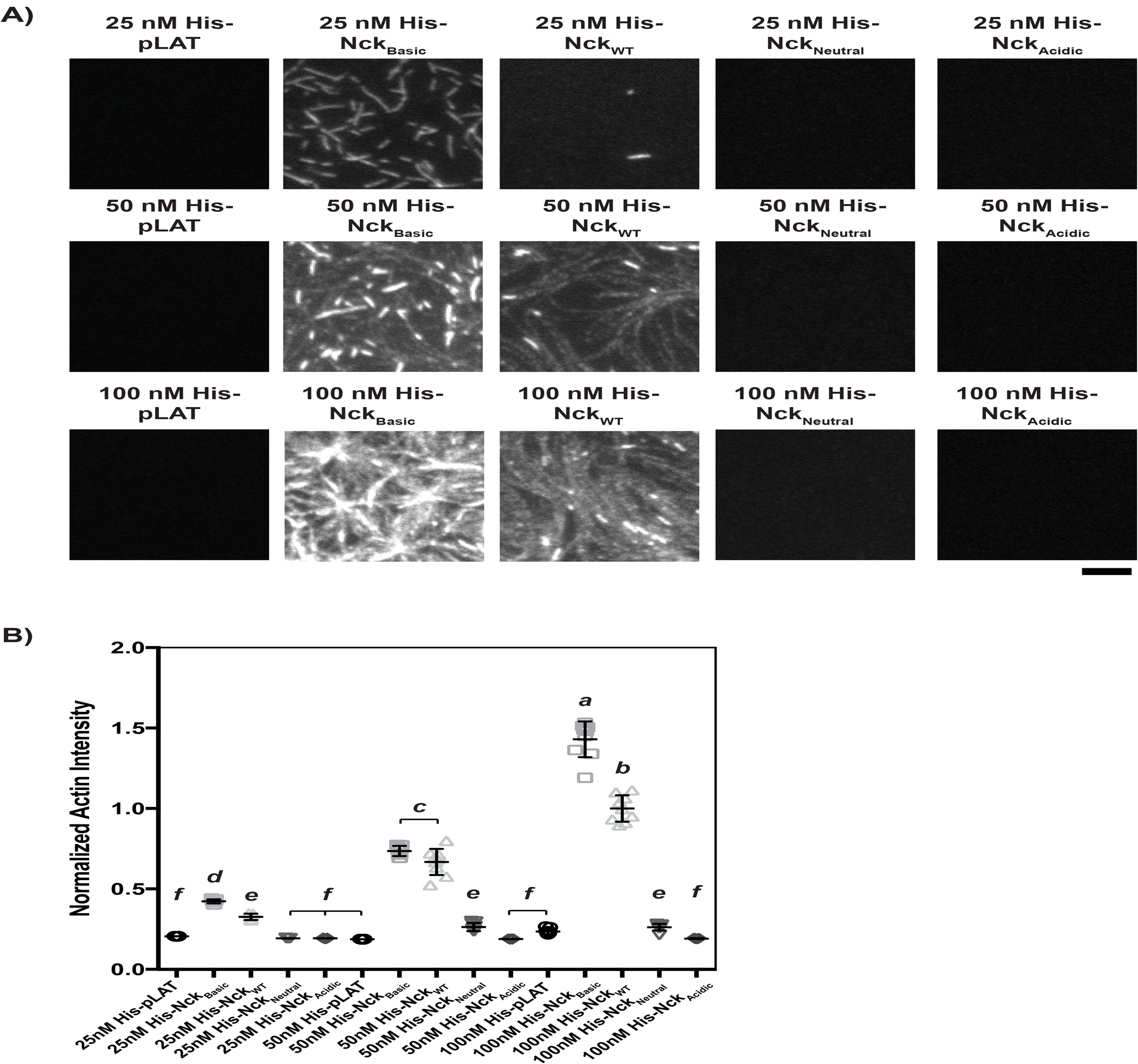
His-tagged Nck variants containing mutations to L1 recruit actin to the bilayer in a density-dependent manner when L1 is WT or contains basic mutations. **(A)** TIRF microscopy images of actin recruited to SLBs coated with His-tagged full-length Nck variants. Concentrations refer to the concentration of His-tagged protein in solution used to coat the bilayer; for pLAT-Alexa488, with 2 % Ni-NTA lipid and His-pLAT-Alexa488 protein in the 0-100 nM range, the density of protein recruited to the bilayer scales approximately linearly with protein concentration in solution (not shown). Scale bar = 5 μm. All images set to same intensity range. **(B)** Normalized fluorescence intensity of rhodamine-actin (normalized to the average actin filament intensity of 100 nM His-Nck_WT_ experiments) recruited to SLBs coated with His-tagged full-length Nck variants. Shown are individual data points and their mean +/- s.d. from N = 15 fields of view from 3 independent experiments (5 FOV per experiment). Significance was determined using analysis of variance with Tukey’s multiple comparison test and found that p < 0.0332 for e vs. f, p < 0.0002 for d vs. e, and p < 0.0001 for a vs. b, a vs. c, a vs. d, a vs. e, a vs. f, b vs. c, b vs. d, b vs. e, b vs. f, c vs. d, c vs. e, and c vs. f.

**Figure 3 – figure supplement 4.**
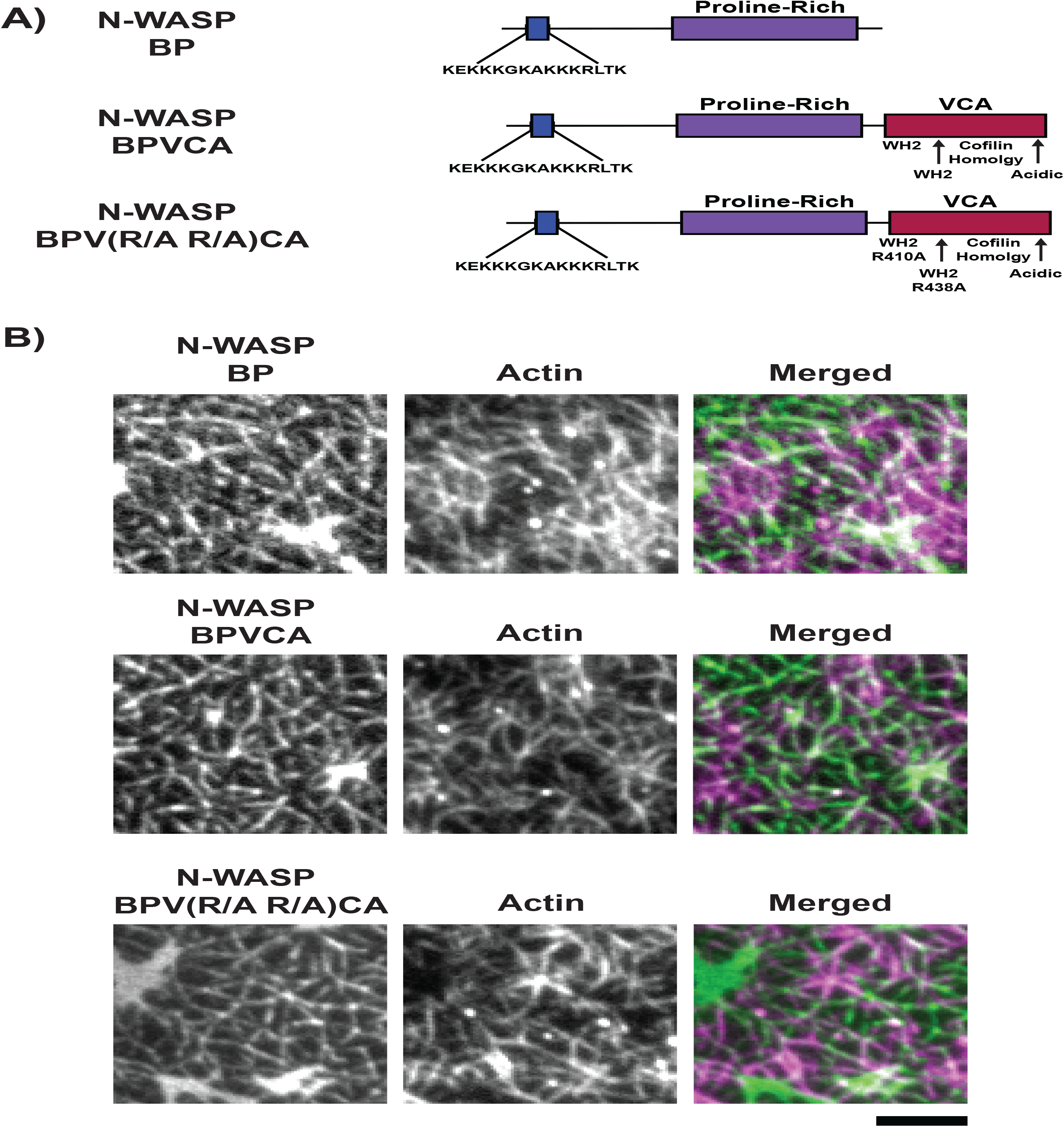
LAT condensates containing N-WASP fragments recruit actin to SLBs. **(A)** Schematics of N-WASP variants used to generate LAT condensates. **(B)** TIRF microscopy images of pLAT ➔ N-WASP variant condensates incubated with rhodamine-labeled actin filaments. Condensates containing N-WASP variants strongly recruited actin filaments to SLBs. Images are representative of 3 independent experiments. Scale bar = 5 μm. All images set to same intensity range.

**Figure 3 – figure supplement 5.**
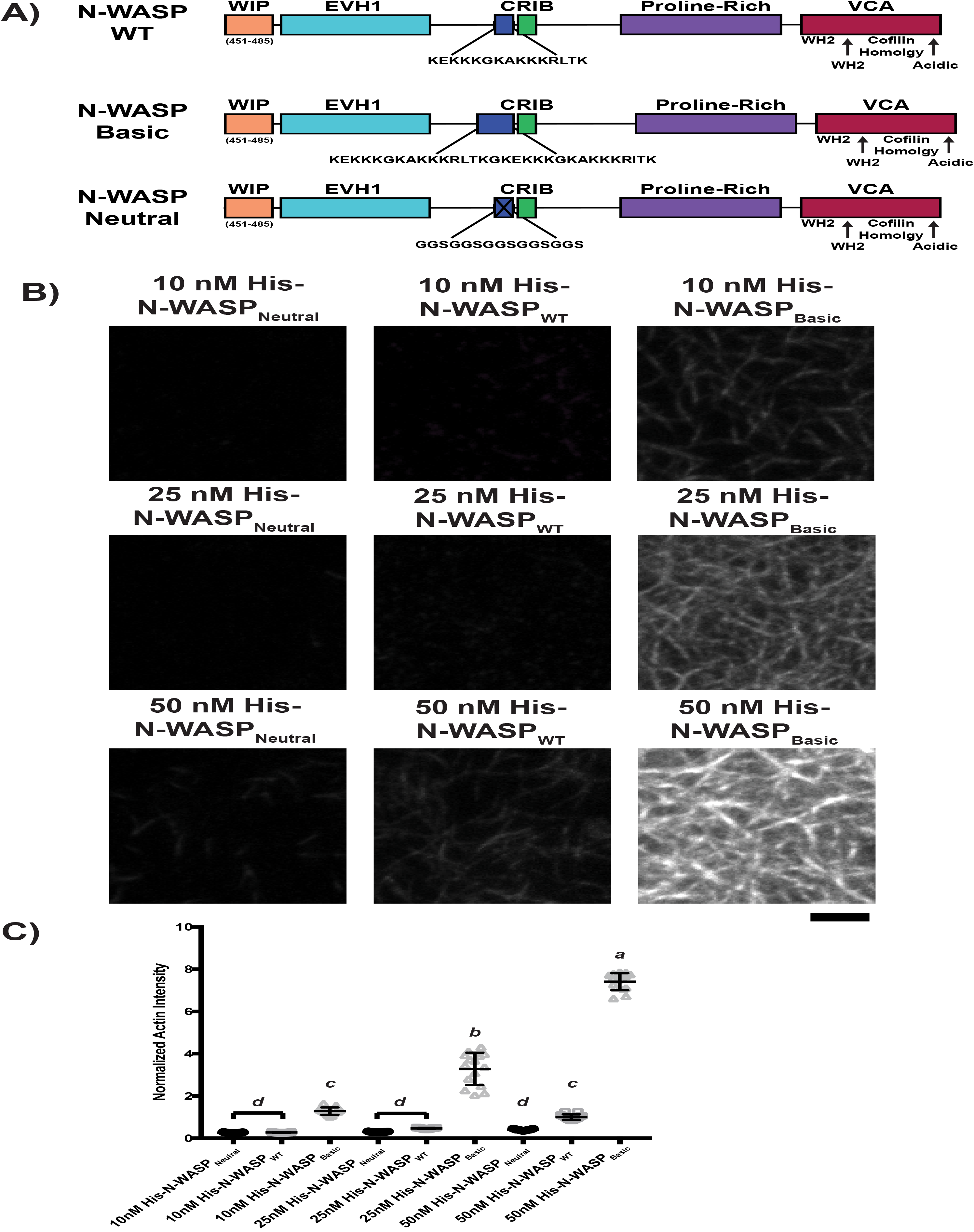
His-tagged N-WASP_Neutral_, N-WASP_WT_, or N-WASP_Basic_ recruit actin filaments to SLBs in a density-dependent manner. **(A)** Schematics of full-length N-WASP variants. **(B)** TIRF microscopy images of rhodamine-labeled actin filaments recruited to SLBs coated with His-tagged full-length N-WASP variants. Concentrations refer to the concentration of His-tagged protein in solution used to coat the bilayer; for pLAT-Alexa488, with 2 % Ni-NTA lipid and His-pLAT-Alexa488 protein in the 0-100 nM range, the density of protein recruited to the bilayer scales approximately linearly with protein concentration in solution. Scale bar = 5 μm. All images set to same intensity range. **(C)** Normalized fluorescence intensity of rhodamine-actin (normalized to the average actin filament intensity of 50 nM His-N-WASP_WT_ experiments) recruited to SLBs coated with His-tagged full-length N-WASP variants. Shown are individual data points and their mean +/- s.d. for N = 15 fields of view from 3 independent experiments (5 FOV per experiment). Significance was determined using analysis of variance with Tukey’s multiple comparison test and found that p < 0.0001 for a vs. b, a vs. c, a vs. d, b vs. c, b vs. d, and c vs. d.

**Figure 4 – figure supplement 1.**
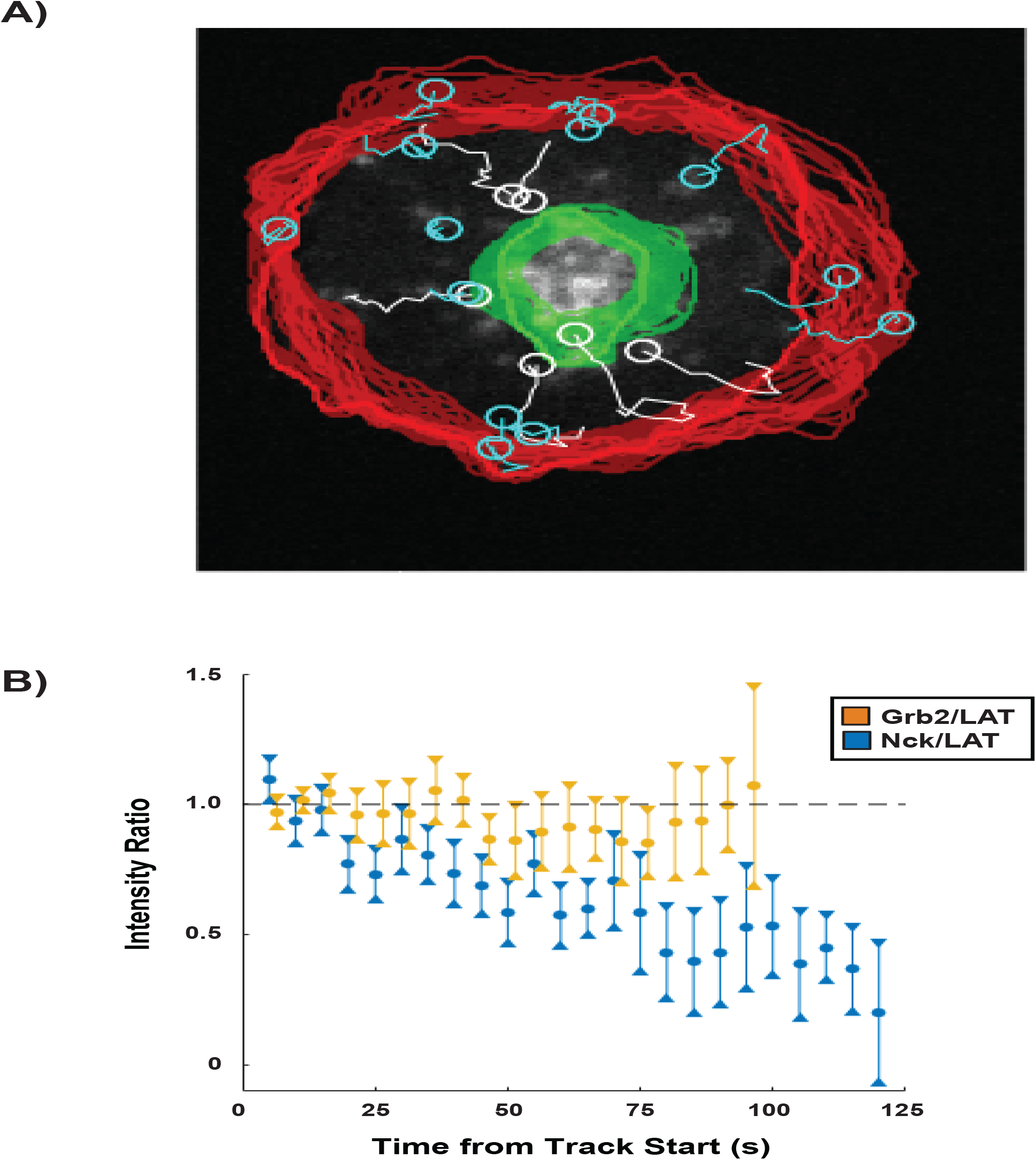
Nck dissipation from condensates is spatially regulated within the first 5 minutes of synapse formation. **(A)** Example of analysis overlayed on a cell showing tracks of individual LAT condensates that move radially over the course of the time-lapse (white), LAT condensates tracks that do not move radially over the course of the time-lapse (cyan), the synapse edge over the course of the time-lapse (red), and the cSMAC over the course of the time-lapse (green). For information regarding detection of the synapse edge and cSMAC see section 4A of the Supplemental Methods. Tracks that did not move radially (cyan) were discarded from the analysis. **(B)** Nck / LAT (blue) and Grb2 / LAT (orange) ratios in condensates plotted against time. The ratio of Grb2: LAT remains constant over time while the ratio of Nck: LAT steadily decreases over time. Plot displays median and its 95% confidence interval from N = 125 condensates from 25 cells expressing Nck-sfGFP and LAT-mCherry from 5 independent experiments and 82 condensates from 11 cells expressing Grb2-mCherry and LAT-mCitrine from 5 independent experiments.

**Figure 4 – figure supplement 2.**
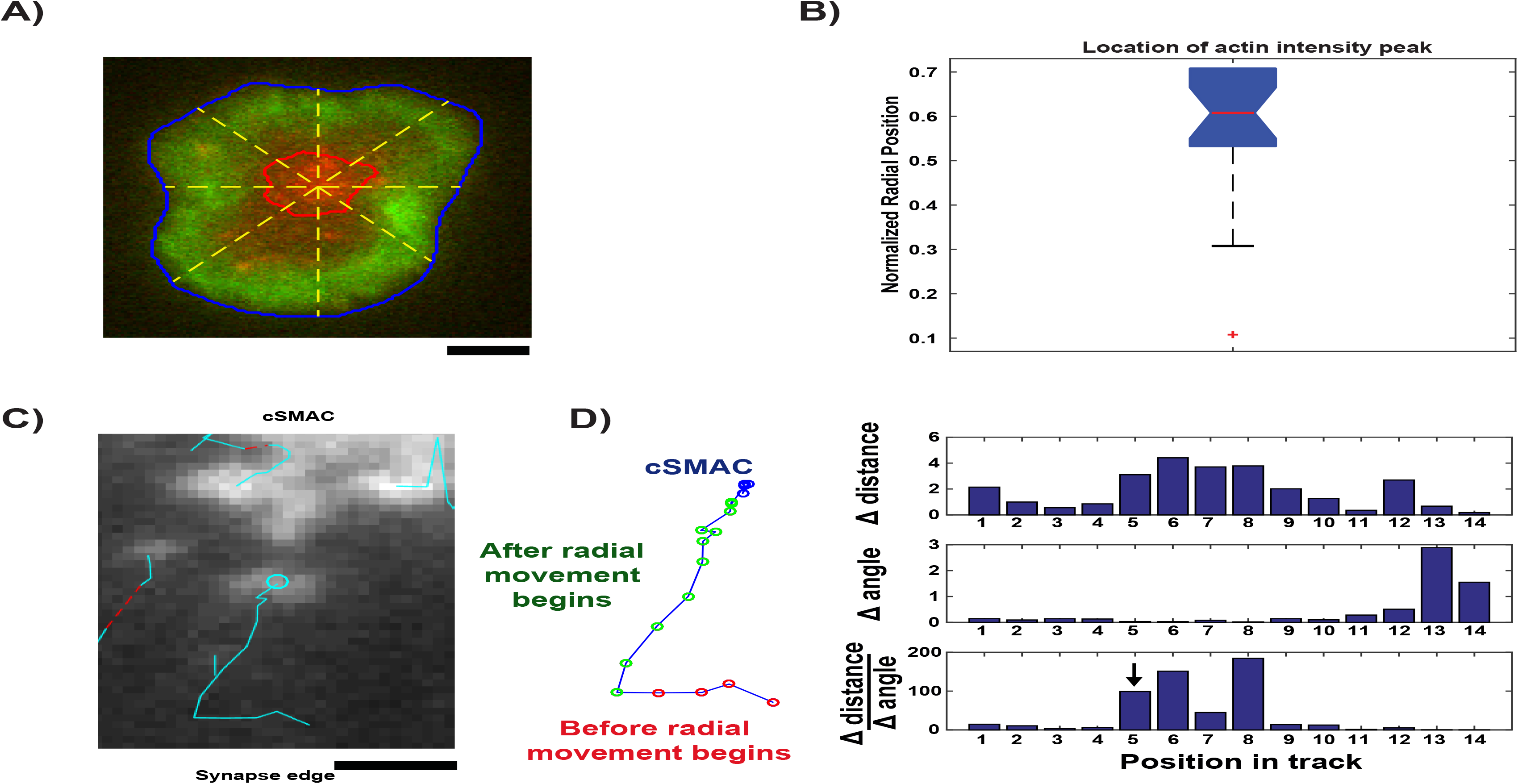
Image analysis for live cell data. **(A)** Example cell (LAT channel: red, LifeAct (actin) channel: green) overlaid with segmented synapse edge (blue line), segmented cSMAC edge (red line), and lines from cSMAC center to synapse edge (yellow dashed lines, 45° angles between neighboring lines) to measure actin intensity radial profile. **(B)** Box plot of the normalized radial position at which the actin intensity profile along each line (yellow dashed lines in **(A)**) reaches its maximum, for each time point in each cell. Box plot description: red central mark shows median; box edges show 25th and 75th percentiles; dashed whiskers extend to the most extreme data points not designated as “outliers”; and notch emanating from median indicates the 95% confidence interval around the median, shown for visual aid. **(C)** Example of condensate that is tracked before and after radial movement. **(D)** (left) Example track that is segmented into 3 distinct regions: before actin engagement (red), after actin engagement (green), and inside cSMAC (blue). (right) Measurements used to detect when condensates begin to move radially: frame-to-frame change in distance (top), frame-to-frame change in angle between consecutive displacements (middle), ratio of previous two measurements (bottom). Arrow indicates position chosen based on the first ratio value in the top 10^th^ percentile.

**Figure 4 – figure supplement 3.**
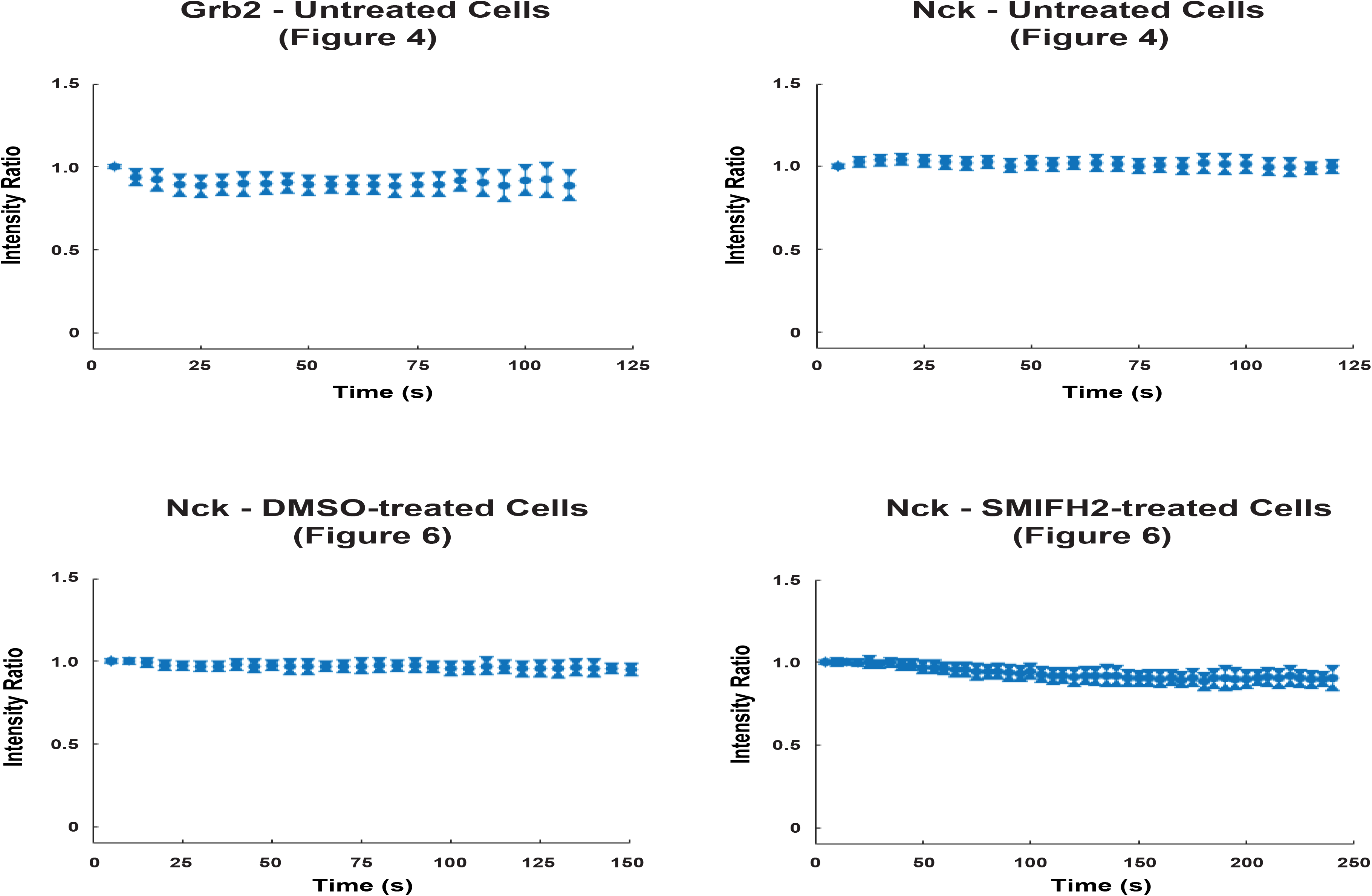
Photobleaching analysis of live-cell data. The mean intensity within the segmented synapse, but outside the segmented cSMAC and detected condensate areas, was taken for each cell for every frame. All mean intensity measurements were normalized by the mean intensity of the first frame for each cell. The ratio of pooled normalized mean intensity (either Grb2: LAT or Nck: LAT) was plotted as a minimal boxplot showing only the median and notches, indicating the 95% confidence interval of each median. This analysis demonstrates that the two channels in live-cell time lapses exhibited similar photobeaching rates (which were overall minimal), and thus variable photobleaching did not contribute to the observed condensate composition changes over time.

**Figure 4 – figure supplement 4.**
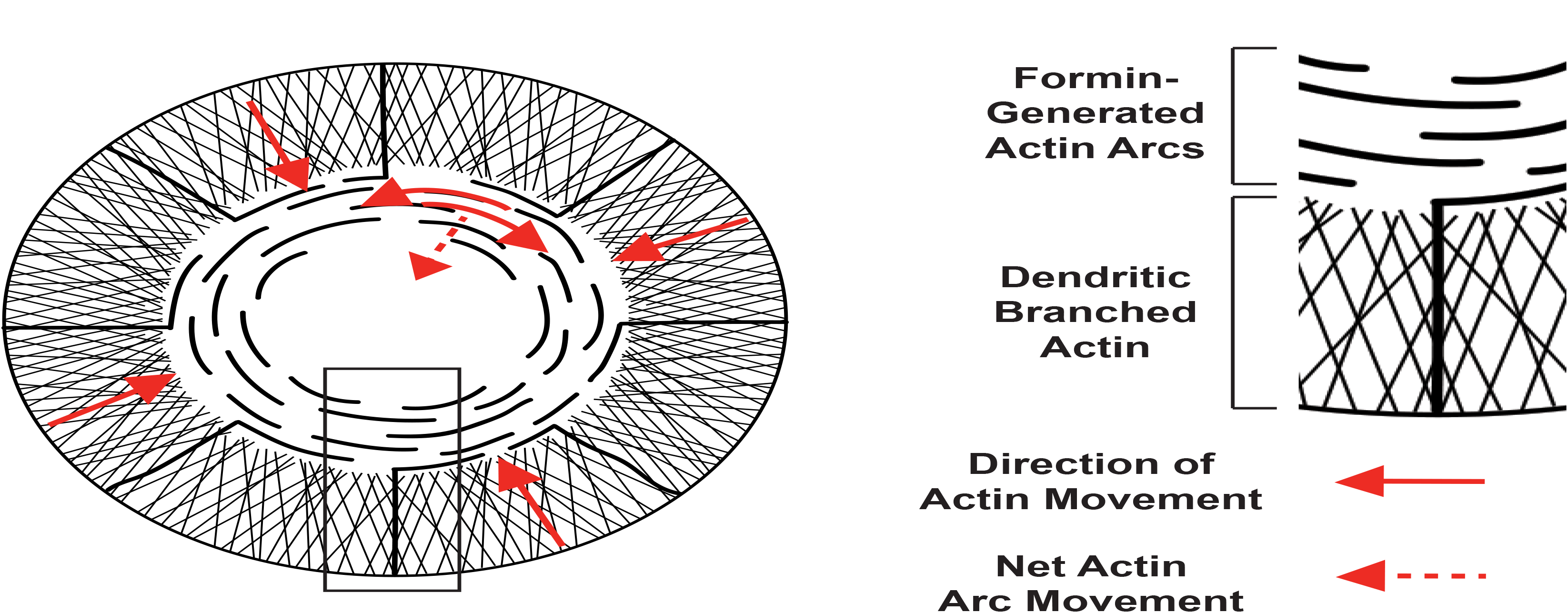
Two actin networks at the IS, with different architecture and movement. (Left) Schematic of actin filament networks and direction of movement at the IS. In the outer region, a dendritic branched actin network undergoes retrograde flow approximately perpendicular to the edge of the synapse (solid red arrows at synapse edge). In the medial region, contractile actin arcs sweep towards the center of the synapse. Actin arcs move parallel to the synapse edge (solid red arrows; filaments sliding against each other through the action of myosin II). Parallel movement results in net movement that is perpendicular to the edge via a telescoping mechanism in which filament motion ‘closes’ the circular architecture (dotted red arrow; much like the diaphragm of a camera). (Right) Enlarged view of boxed region from schematic on left with dendritic branched actin network and formin-generated actin arcs.

**Figure 4 – figure supplement 5.**
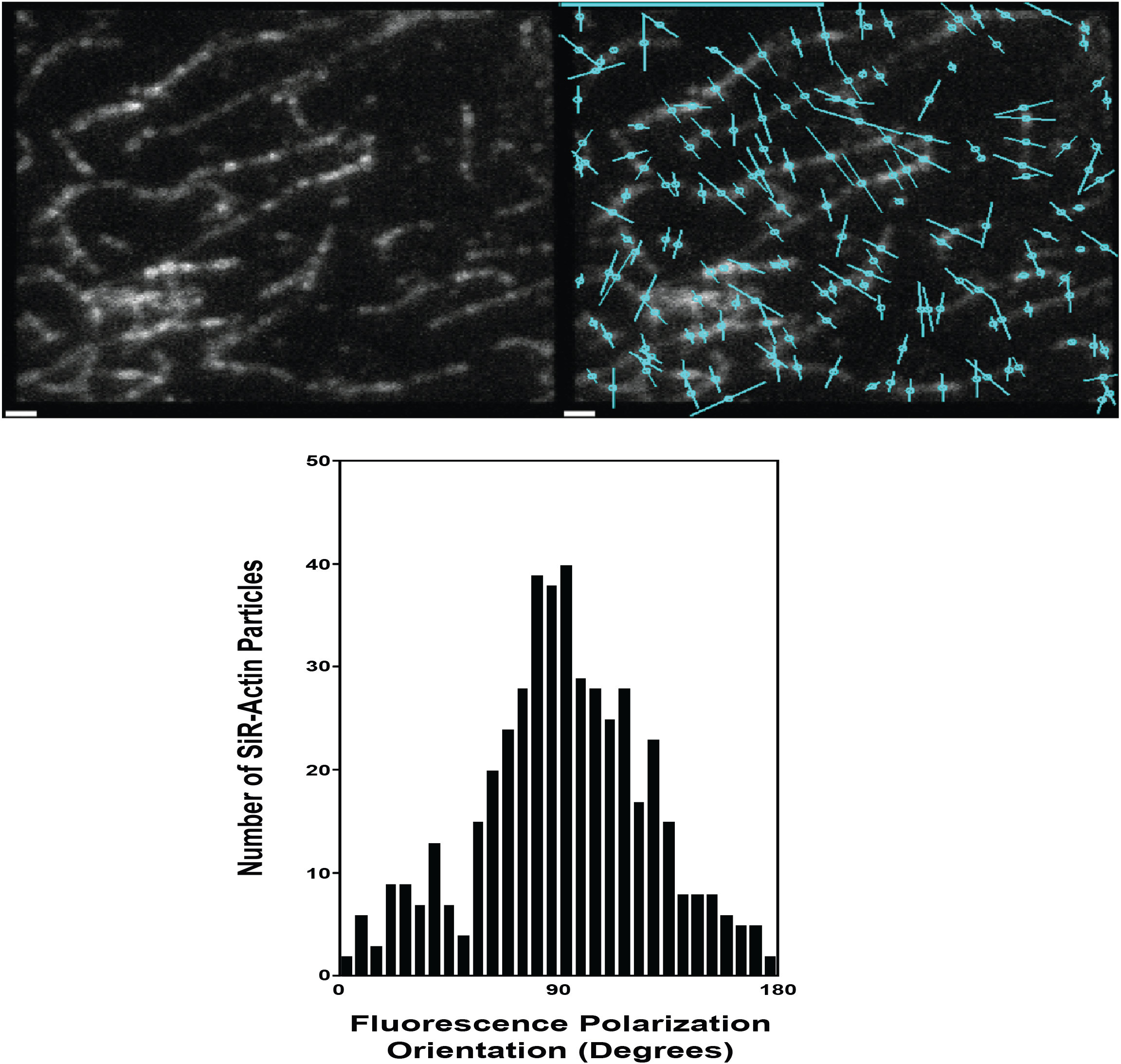
SiR-Actin polarity is perpendicular to actin filaments. (top, left) SiR-Actin labeling of immobilized actin filaments attached to a glass coverslip. (top, right) The polarity of SiR-Actin bound to immobilized actin filaments is shown by circles transversed by a line to indicate the dye polarity (cyan). (bottom) Histogram showing the orientation of SiR-Actin bound to actin filaments with respect to the alignment of the actin filament. Scale Bar = 1 μm.

**Figure 4 – figure supplement 6.**
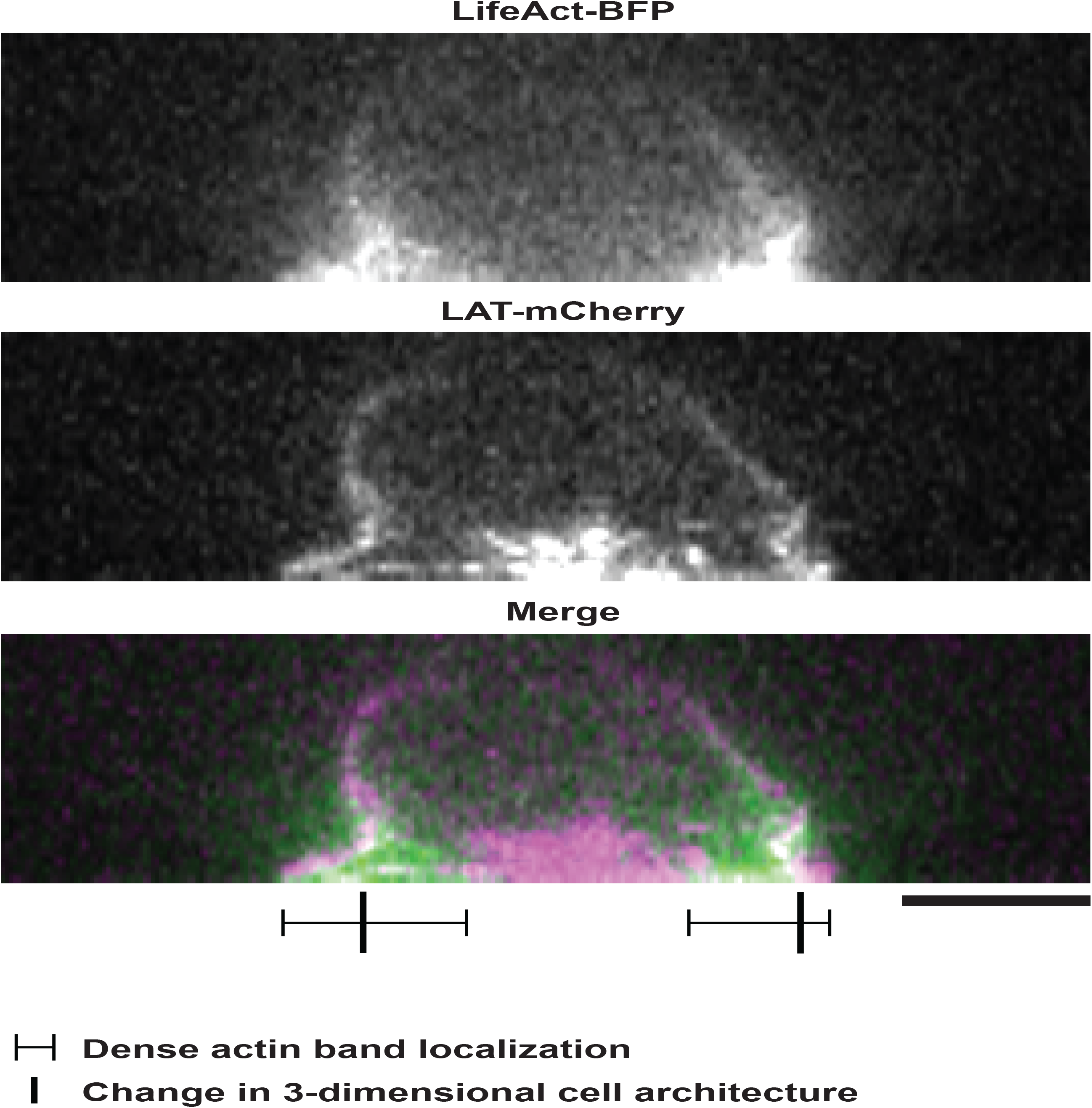
The two-dimensional extension of the dense actin network at the IS in Jurkat T cells activated on SLBs is independent of the three dimensional shape of the activated Jurkat T cell. Scale Bar = 5 μm.

**Figure 5 – figure supplement 1.**
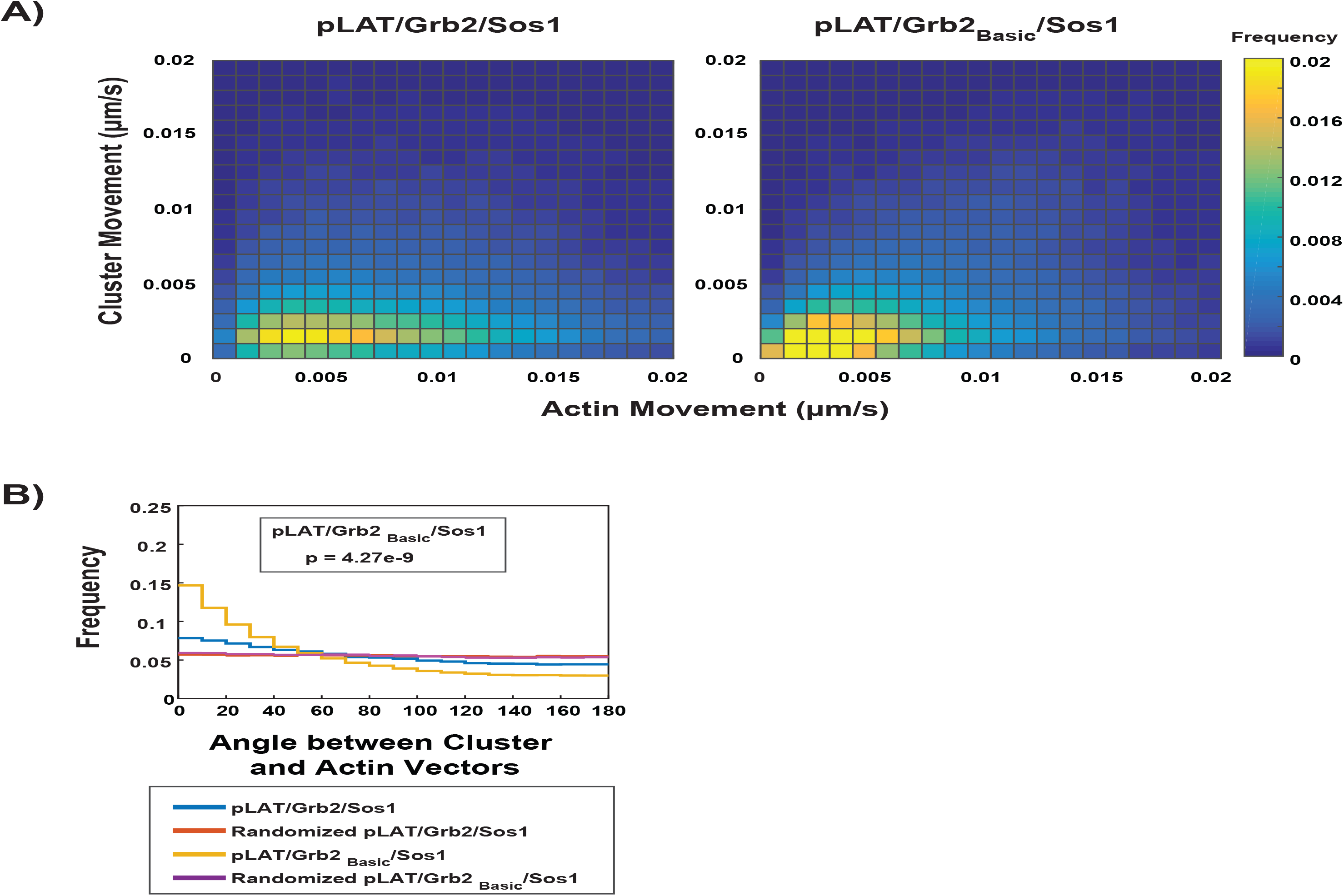
Adding a doubled N-WASP basic domain to Grb2 (Grb2_Basic_) enables pLAT / Grb2_Basic_ / Sos1 condensates to move with actin to a better degree than condensates formed with WT Grb2. **(A)** Condensate speed vs. actin speed for pLAT / Grb2 / Sos1 (from Fig. 3) or pLAT / Grb2_Basic_ / Sos1 condensates in contracting actin networks from STICS analysis. Type of condensate indicated above each heat map. Heat map indicates frequency in each bin, i.e. counts in each bin normalized by total number of counts. **(B)** Distribution of the angle between actin and condensate movement vectors for pLAT / Grb2 / Sos1 (blue, from Fig. 3), pLAT / Grb2_Basic_ / Sos1 (gold), randomized pLAT / Grb2 / Sos1 (red, from Fig. 3), or randomized pLAT / Grb2_Basic_ / Sos1 (purple). Data in **(A)** and **(B)** were pooled from N = 15 fields of view from 3 independent experiments (5 FOV per experiment).

**Figure 6 – figure supplement 1.**
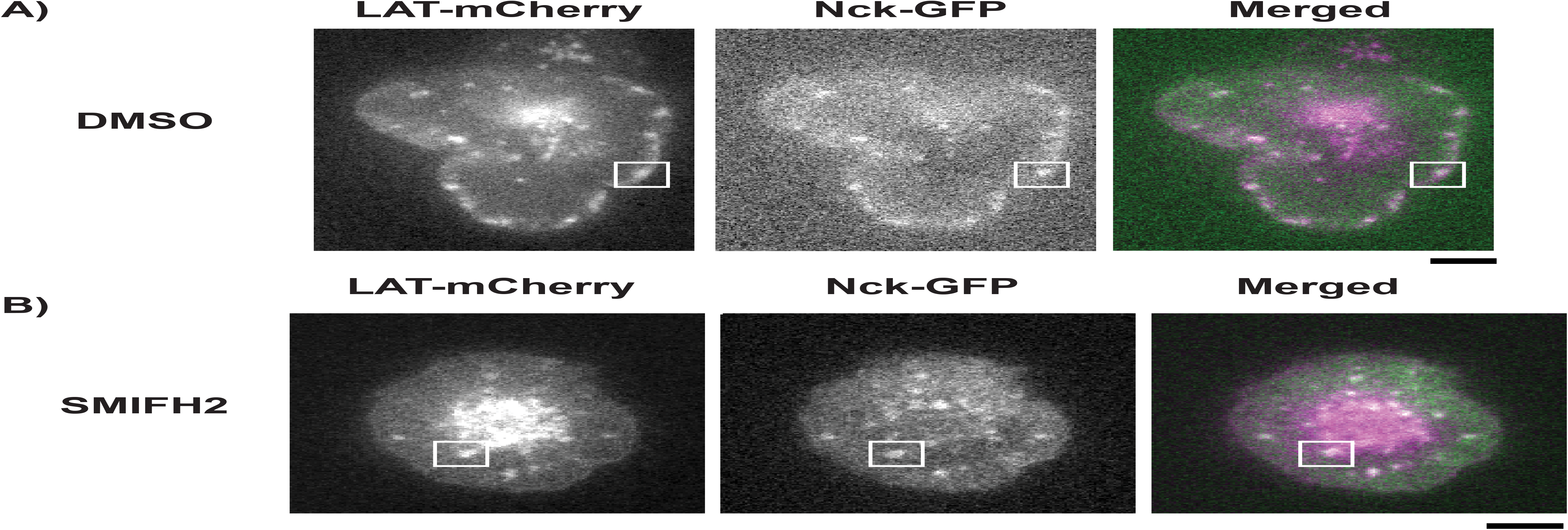
Example images showing DMSO- or SMIFH2-treated Jurkat T cells activated on SLBs. **(A, B)** TIRF microscopy images of Jurkat T cells expressing Nck-sfGFP and LAT-mCherry and treated with DMSO **(A)** or the formin inhibitor SMIFH2 **(B)** for 5 minutes prior to activation on SLBs. Scale Bars = 5 μm.

**Video 1. Reconstitution of LAT condensate formation and movement using different component mixtures.** TIRF microscopy revealed LAT condensate formation and movement on supported lipid bilayers. His_8_-pLAT-Alexa488 was attached to Ni-functionalized SLBs at 500 molecules / μm^2^. Condensate component mixtures are indicated at the top of each movie panel. All reconstitutions were performed in 75 mM KCl. Condensates were formed by adding 125 nM Grb2 and 125 nM Sos1 (pLAT ➔ Sos1), 125 nM Grb2, 62.5 nM Sos1, and 62.5 nM pSLP-76 (pLAT ➔ pSLP-76), 125 nM Grb2, 62.5 nM Sos1, 62.5 nM pSLP-76, and 125 nM Nck (pLAT ➔ Nck), or 125 nM Grb2, 62.5 nM Sos1, 62.5 nM pSLP-76, 125 nM Nck, and 125 nM N-WASP (pLAT ➔ N-WASP). After formation, most condensates were either immobile or displayed confined movement regardless of their composition. Movie shows a 32 μm x 32 μm field of view for each movie panel. The movie is played at 9 fps with frame intervals of 15 seconds.

**Video 2. Reconstitution of LAT condensate formation and movement using different component mixtures in actin networks.** TIRF microscopy revealed LAT condensate formation and movement on supported lipid bilayers in an actin network. His_8_-pLAT-Alexa488 (green) was attached to Ni-functionalized SLBs at 500 molecules / μm^2^. Actin filaments (magenta) were attached to the same bilayers via His_10_-ezrin actin binding domains. Condensate component mixtures are indicated at the top of each movie panel. All reconstitutions were performed in 75 mM KCl. Condensates were formed by adding 125 nM Grb2 and 125 nM Sos1 (pLAT ➔ Sos1), 125 nM Grb2, 62.5 nM Sos1, and 62.5 nM pSLP-76 (pLAT ➔ pSLP-76), 125 nM Grb2, 62.5 nM Sos1, 62.5 nM pSLP-76, and 125 nM Nck (pLAT ➔ Nck), or 125 nM Grb2, 62.5 nM Sos1, 62.5 nM pSLP-76, 125 nM Nck, and 125 nM N-WASP (pLAT ➔ N-WASP). After formation, pLAT ➔ Sos1 and pLAT ➔ pSLP-76 condensates displayed increased mobility (compared to condensates formed in the absence of actin filaments) while pLAT ➔ Nck and pLAT ➔ N-WASP showed decreased mobility (compared to condensates formed in the absence of actin networks) while binding to and wetting actin filaments. Movie shows a 32 μm x 32 μm field of view for each movie panel. The movie is played at 9 fps with frame intervals of 15 seconds.

**Video 3. Reconstitution of LAT condensate formation and movement using different component mixtures in steady-state active actomyosin networks.** TIRF microscopy revealed LAT condensate formation and movement on supported lipid bilayers in an active actomyosin network. His_8_-pLAT-Alexa488 (green) was attached to Ni-functionalized SLBs at 500 molecules / μm^2^. Actin filaments (magenta) were attached to the same bilayers via His_10_-ezrin actin binding domains. 100 nM myosin II and 1 mM ATP were added to actin filaments prior to LAT condensate formation to induce actin filament movement. Condensate component mixtures are indicated at the top of each movie panel. All reconstitutions were performed in 75 mM KCl. Condensates were formed by adding 125 nM Grb2 and 125 nM Sos1 (pLAT ➔Sos1), 125 nM Grb2, 62.5 nM Sos1, and 62.5 nM pSLP-76 (pLAT ➔ pSLP-76), 125 nM Grb2, 62.5 nM Sos1, 62.5 nM pSLP-76, and 125 nM Nck (pLAT ➔ Nck), or 125 nM Grb2, 62.5 nM Sos1, 62.5 nM pSLP-76, 125 nM Nck, and 125 nM N-WASP (pLAT ➔ N-WASP). After formation, all condensate types displayed increased condensate mobility (compared to condensates formed in the absence of actin filaments or in an actin network). Movie shows a 32 μm × 32 μm field of view for each movie panel. The movie is played at 9 fps with frame intervals of 15 seconds.

**Video 4. Reconstitution of LAT condensate movement using different component mixtures in contracting actomyosin networks.** TIRF microscopy revealed actin filament and LAT condensate movement on supported lipid bilayers in a contracting actomyosin network. His_8_-pLAT-Alexa488 (green) was attached to Ni-functionalized SLBs at 500 molecules / μm^2^. Actin filaments (magenta) were attached to the same bilayers via His_10_-ezrin actin binding domains. Condensate component mixtures are indicated at the top of each movie panel. All reconstitutions were performed in 50 mM KCl. Condensates were formed by adding 125 nM Grb2 and 125 nM Sos1 (pLAT ➔ Sos1) or 125 nM Grb2, 62.5 nM Sos1, 62.5 nM pSLP-76, 125 nM Nck, and 125 nM N-WASP (pLAT ➔ N-WASP). After formation, pLAT ➔ Sos1 condensates were randomly distributed within the actin network while pLAT à N-WASP condensates bound to and wet actin filaments. Actin filament contraction was induced by adding 100 nM myosin II. pLAT➔ Sos1 condensates did not efficiently move with contracting actin filaments while pLAT ➔ N-WASP condensates efficiently moved with contracting actin filaments. Movie shows a 52 μm × 52 μm field of view for each movie panel. The movie is played at 5 fps with frame intervals of 5 seconds.

**Video 5. Reconstitution of LAT condensate movement using different component mixtures containing N-WASP_Neutral_ or N-WASP_Basic_ in contracting actomyosin networks.** TIRF microscopy revealed actin filament and LAT condensate movement on supported lipid bilayers in a contracting actomyosin network. His_8_-pLAT-Alexa488 (green) was attached to Ni-functionalized SLBs at 500 molecules / μm^2^. Actin filaments (magenta) were attached to the same bilayers via His_10_-ezrin actin binding domains. Condensate component mixtures are indicated at the top of each movie panel. All reconstitutions were performed in 50 mM KCl. Condensates were formed by adding 125 nM Grb2, 62.5 nM Sos1, 62.5 nM pSLP-76, 125 nM Nck, and 125 nM N-WASP_Neutral_ (pLAT ➔ N-WASP_Neutral_) or 125 nM Grb2, 62.5 nM Sos1, 62.5 nM pSLP-76, 125 nM Nck, and 125 nM N-WASP_Basic_ (pLAT ➔ N-WASP_Basic_). After formation, both types of condensates bound to and wet actin filaments, although pLAT ➔ N-WASP_Neutral_ condensates did not wet filaments to the same degree as either pLAT ➔ N-WASP_WT_ condensates (compare with Movie S6) or pLAT ➔ N-WASP_Basic_. Actin filament contraction was induced by adding 100 nM myosin II. pLAT➔ N-WASP_Neutral_ condensates did not efficiently move with contracting actin filaments (although they moved with actin filaments to a greater degree than pLAT ➔ Sos1 condensates [compare with Movie S6]) while pLAT ➔ N-WASP_Basic_ condensates efficiently moved with contracting actin filaments. Movie shows a 52 μm × 52 μm field of view for each movie panel. The movie is played at 5 fps with frame intervals of 5 seconds.

**Video 6. Single particle tracking of LAT condensates in activated Jurkat T cells.** TIRF microscopy of an activated Jurkat T cell expressing LAT-mCherry on a SLB coated with ICAM-1 and OKT3. LAT condensates form at the synapse edge (cyan perimeter) and are tracked, using uTrack, as they move towards the cSMAC. Track colors: Cyan-track segment between adjacent frames; Red-gap closing between non-adjacent frames (maximum of 3 frames between end of a track segment and the beginning of another). Showing a maximum track tail length of 10 frames for visual clarity. The movie is played at 5 fps with frame intervals of 5 seconds.

**Video 7. Grb2 is maintained as LAT condensates move across the IS in activated Jurkat T cells.** TIRF microscopy of an activated Jurkat T cell expressing Grb2-mCherry (magenta) and LAT-mCitrine (green) on a SLB coated with ICAM-1 and OKT3 revealed that Grb2 co-localizes with LAT condensates as they move from the edge of the synapse to the cSMAC. Movie shows a 22 μm × 22 μm field of view. The movie is played at 7 fps with frame intervals of 5 seconds.

**Video 8. Nck dissipates from LAT condensates as they move across the IS in activated Jurkat T cells.** TIRF microscopy of an activated Jurkat T cell expressing LAT-mCherry (magenta) and Nck-sfEGFP (green) on a SLB coated with ICAM-1 and OKT3 revealed that Nck dissipates from LAT condensates as they move from the edge of the synapse to the cSMAC. Movie shows a 24 μm × 24 μm field of view. The movie is played at 5 fps with frame intervals of 5 seconds.

**Video 9. Reconstitution of movement in LAT condensates containing Grb2_Basic_ in contracting actomyosin networks.** TIRF microscopy revealed actin filament and LAT condensate movement on supported lipid bilayers in a contracting actomyosin network. His_8_-pLAT-Alexa488 (green) was attached to Ni-functionalized SLBs at 500 molecules / μm^2^. Actin filaments (magenta) were attached to the same bilayers via His_10_-ezrin actin binding domains. Reconstitution was performed in 50 mM KCl. Condensates were formed by adding 125 nM Grb2_Basic_ and 125 nM Sos1. After formation, condensates bound to and wet actin filaments. Actin filament contraction was induced by adding 100 nM myosin II. LAT condensates composed of Grb2_Basic_ moved with contracting actin filaments, although not to the same degree as pLAT ➔ N-WASP_WT_ condensates (Compare with Movie S6). Movie shows a 52 μm × 52 μm field of view. The movie is played at 5 fps with frame intervals of 5 seconds.

**Video 10. LAT condensates containing Grb2_Basic_ deviate tend to deviate from a radial trajectory across the IS.** TIRF microscopy of an activated Jurkat T cell expressing Grb2_Basic_-mCherry and LAT-mCitrine on a SLBcoated with ICAM-1 and OKT3 revealed that condensates containing Grb2_Basic_-mCherry tend to deviate from a radial trajectory as they move across the IS (Compare with Movie S1). Movie shows a 35 μm × 35 μm field of view. The movie is played at 6 fps with frame intervals of 5 seconds.

**Video 11. Nck dissipates from LAT condensates as they move across the IS in activated Jurkat T cells treated with DMSO.** TIRF microscopy of an activated Jurkat T cell expressing LAT-mCherry (magenta) and Nck-sfGFP (green) on a SLB coated with ICAM-1 and OKT3 revealed that Nck dissipates from LAT condensates as they move from the edge of the synapse to the cSMAC following treatment with DMSO for 5 minutes prior to activation. Movie shows a 31 μm × 31 μm field of view. The movie is played at 6 fps with frame intervals of 5 seconds.

**Video 12. Nck is maintained in LAT condensates as they move across the IS in activated Jurkat T cells treated with the formin inhibitor SMIFH2.** TIRF microscopy of an activated Jurkat T cell expressing LAT-mCherry (magenta) and Nck-sfGFP (green) on a SLB coated with ICAM-1 and OKT3 revealed that Nck is maintained in LAT condensates as they move from the synapse of the cell to the cSMAC following treatment with SMIFH2 for 5 minutes prior to activation. Movie shows a 27 μm × 27 μm field of view. The movie is played at 10 fps with frame intervals of 5 seconds.

